# Bayesian Network Structure Learning: The New Calibrated Minimum Uncertainty Criterion and Discriminative Power Evaluation Framework

**DOI:** 10.1101/2025.10.13.681946

**Authors:** Grigoriy Gogoshin, Andrei S Rodin

## Abstract

Bayesian network (BN) structure learning (BNSL) from heterogeneous data is a classical problem in probabilistic machine learning and knowledge discovery. A variety of computational methods exist for score-based BNSL, but all inherit limitations imposed by the underlying model selection paradigms. With finite, often scarce data, imperfect conversion of observations into structural evidence can lead to miscalibration, structural bias, amplification of statistical errors, and unstable or poorly interpretable learned networks. The recently introduced Minimum Uncertainty (MU) principle was proposed to mitigate these problems for discrete BNSL, but its scoring criterion remained a preliminary prototype requiring generalization and calibration. Here, we derive a resolution-aware, calibrated MU criterion under broad perturbation assumptions and develop a framework for evaluating the discriminative power of BN scoring criteria. Across parameter regimes and dependence modes, comparison with MDL, BDeu, and K2 shows that the calibrated MU criterion maintains balanced operating characteristics and improves structure recovery performance, with negligible computational overhead; these gains are also confirmed on real biomedical data. The evaluation framework additionally provides edge-level specificity, sensitivity, and evidence-quality estimates, supporting both practical interpretation of learned networks and principled comparison of scoring criteria. Together, the calibrated MU criterion and the discriminative power evaluation framework provide a general approach for more robust, interpretable, and statistically calibrated score-based structure learning across diverse settings and domains.

## 1 Background

Bayesian networks (BNs) have become important tools for knowledge discovery, elucidating the intricate web of interactions that govern complex multiscale systems across many diverse domains. These probabilistic graphical models represent conditional dependencies among the variables of interest, facilitating diagnostic and predictive inference, modularization of knowledge, and surveying evidence propagation flows. In contemporary biomedicine, BNs enable the development of more effective approaches to dissecting and manipulating biological systems *in silico* and, ultimately, the identification of novel therapy targets and strategies. In computer vision and econometrics, they offer a way to deal with uncertain measurements and factors when making inferences about the global state of the system. Whatever the domain, BN modeling transforms a complex multivariate dependence into a transparent yet principled representation.

Unlike approaches that rely solely on pre-existing knowledge or curated knowledge graphs, reconstructing BN models directly from the data is primarily a machine learning and data mining task. Unlike many machine learning/deep learning methods, however, BN modeling offers native interpretability, enabling researchers to move beyond predictive accuracy and descriptive statistics toward hypothesizing and delineating underlying mechanisms (Hammond and Smith 2025; Canonaco et al. 2025), with a principled bridge to granular causal inference.

The value of this methodology is substantial — and growing — across diverse application domains. In biomedicine, recent applications range from multimodal oncology, nephrology, critical care, and mental health research to population mapping, health monitoring, and adverse outcome pathway modeling, to name just a few (Zhang et al. 2025; Briganti et al. 2024; Hong et al. 2025; Wang et al. 2025; Jeong et al. 2025; Zhang et al. 2026; Petousis et al. 2026; Tong et al. 2026). However, reconstructing BN models from data is far from straightforward. Limited sample sizes and the non-uniqueness of models that equally explain the observations (parameter/structure equivalence) can yield inaccurate or unstable reconstructions. This inherent non-identifiability introduces uncertainty about the system’s true underlying structure. Recent work has further shown that BN modeling can be surprisingly sensitive to even seemingly incidental aspects of the learning procedure, such as variable ordering (Kitson and Constantinou 2024). Hence, rigorous assessment and improvement of accuracy, precision, and recall are essential to broaden adoption and enhance the practical utility of BN modeling across domains and disciplines.

Among the BN design choices, a principled, consistent model selection criterion is pivotal to achieving models that balance fidelity, parsimony, and usability. However, many model selection criteria, despite their relative effectiveness, have limitations and irregularities that can hinder reliable model selection (Suzuki 2016; Scutari 2018, 2016; de Campos et al. 2009; Gogoshin and Rodin 2025). A given scoring criterion may be too inflexible with respect to symmetry or asymptotic constraints in settings where such abstract requirements may be unattainable or irrelevant due to a strictly limited sample size. Or it may have a tendency to over-prioritize a single ad hoc aspect like parsimony, undervaluing subtle but critical details hidden under the more obvious layer of relationships. By construction, unguided generic criteria are usually too abstract to integrate concrete contextual information and, therefore, often lack adequate empirical discriminative resolution. These limitations can lead to model selection that performs well superficially yet has low precision, creates epistemic friction, and is ultimately misleading. More fundamentally, existing scoring paradigms do not explicitly calibrate model selection to the empirical statistical resolution at which dependencies can actually be distinguished from sampling uncertainty.

Nevertheless, some criteria have been quite successful in various domains and real-world applications. The popular Minimum Description Length (MDL) criterion is one such practical, “plug-and-play” score with a strong track record in performance and computational efficiency. It seeks an optimal balance between data fidelity and parsimony.

The classical statistical variant of MDL is formulated in terms of the appropriate trade-off between the description length of the data given the model and the description length of the model itself (de L.M. Campos 2006). The core components of this variant are the log-likelihood *LL* that serves as a measure of the description length of the data given the model, and a measure of model complexity *C* proportional to the degrees of freedom required for the representation of the factorization of the joint probability

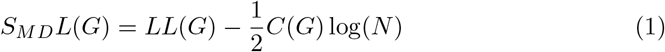

where *G* is a graph defined on the set of variables 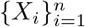 and *N* is the sample size of the associated dataset. The log-likelihood can be conveniently written in terms of the information-theoretic entropy *H*

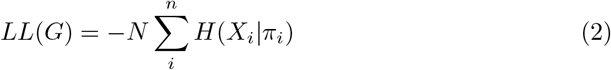

where *π*_*i*_ is the ancestor sets of *X*_*i*_, and N is the size of the sample. If an *r*_*i*_-variate *X*_*i*_ has *q*_*i*_-variate parent set *π*_*i*_, the complexity of the network is given by

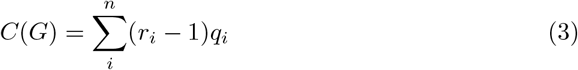

Like many other criteria, MDL addresses the BN structure learning (BSNL) by solving an optimization problem, seeking the solution *G*^∗^ that maximizes the score

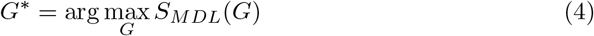

Rescaling *S*_*MDL*_ by −1*/N* strips the first term from the sample size dependence, yielding an elegant and convenient information-theoretic criterion

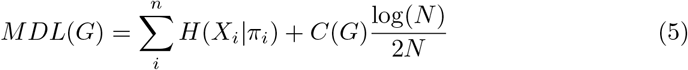

and reconfigures the MDL-based BNSL into the equivalent minimization problem

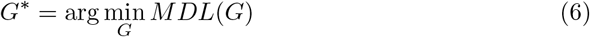

It’s worth noting that in the BNSL context MDL score is asymptotically equivalent to the Bayesain Information Criterion (BIC) even though they are derived from independent foundational principles, targeting maximum compression and maximum posterior probability respectively.

Despite its elegance, MDL — like many well-established criteria — exhibits specific shortcomings which become particularly noticeable in the context of cross-comparison of homologous interactions across a variety of datasets. For example, MDL exhibits strong dependence on the sample size for nonlinear (non-pairwise) configurations (Theorem 1 in de Campos et al. (2009)). A notable and undesirable consequence of this is its complexity-dependent miscalibration of the cost term that tends to over-penalize variables with higher cardinality, setting strict parent set bounds and signalling a specificity-sensitivity imbalance (see Section 2.2 for a concrete example).

This and other similar difficulties motivated the work in Gogoshin and Rodin (2025) which proposed the *Minimum Uncertainty* (MU) model selection principle as a way to derive the components of the objective function naturally. There, contextual statistical resolution considerations are integrated directly into the selection principle; a preliminary formulation of the corresponding scoring criterion demonstrates a proof of concept in BNSL with the desired properties.

Here, we give the MU principle a firmer footing by deriving a new, generalized and calibrated scoring criterion under broad perturbation assumptions that reflect the majority of empirically reasonable scenarios. Furthermore, we develop a dedicated dependency model for evaluating the discriminative power profile of the new score. Finally, we contrast the operating characteristics of several conventional scoring functions and the MU criterion across varying parameter regimes, and show that the resulting framework supports edge-level specificity, sensitivity, and evidence-quality estimates in practical BNSL. Thus, beyond the MU criterion itself, the present work establishes a general empirical framework for assessing and interpreting score-based structure learning across diverse settings.

For completeness, Sections 2.1–2.3 briefly recapitulate the conventional scoring paradigms and the original MU principle formulation; the methodological novelties of the present work begin in Sections 2.4–2.6

## 2 Methods

### 2.1 MDL and BD as Decision Tests

BN reconstruction from data utilizing the MDL score attempts to solve the optimization problem *G*^∗^ = arg max_*G*_ *MDL*(*G*).

To do so the process usually evaluates the improvement obtained by a test network structure *G*^*′*^ over the accepted structure *G* as scored by the MDL criterion

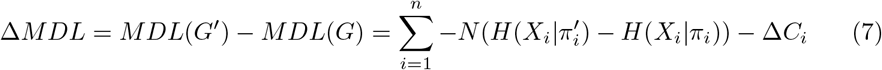

where

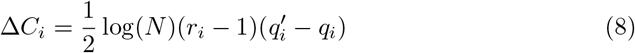

The BN structure is updated to *G*^*′*^ whenever Δ*MDL >* 0. Assuming that *G*^*′*^ adds an edge to *G*, the local conditional independence criterion for a new member of the ancestor set 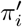 of *X*_*i*_ can be formulated in terms of entropy as

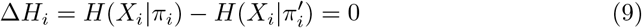

Note that if 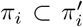 then 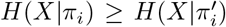. Hence, the condition triggering the update from *G* to *G*^*′*^ is that there is at least one such *i* for which

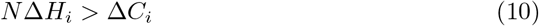

i.e. for at least one new candidate edge the deviation from conditional independence must be above the threshold given by Δ*C*_*i*_.It follows that, for many information-theoretic model selection criteria with an additive penalty, finding the optimal structure is a matter of identifying and ordering conditional independence specifications of the model that can be framed as decision-theoretic task.

As will be shown below, a similar claim can be made about many posterior-probability derived criteria as well. Consider the BD family of scores which targets the solution *G*^∗^ of the optimization problem

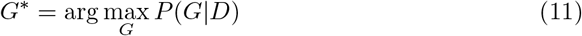

that maximizes the posterior probability, given some dataset *D*. Since *P* (*G*|*D*) = *P* (*G, D*)*/P* (*D*) and the data prior *P* (*D*) is assumed to be the same for all structures it is sufficient to evaluate *P* (*G, D*), i.e.

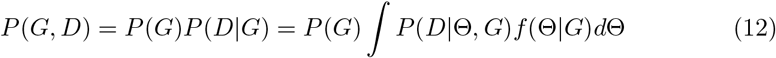

where Θ are conditional probabilities associated with *G, P* (*D*|Θ, *G*) is the likelihood of the data given the network configuration and *f* (Θ|*G*) is the prior density of its parameters. Assuming the structure prior *P* (*G*) is uniform, *P* (*G, D*) can be stated as

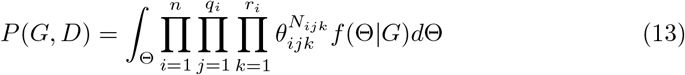

where *n* is the number of *r*_*i*_-variate variables *X*_*i*_ in the data, *q*_*i*_ is the number of unique instantiations *w*_*ij*_ of the ancestors *π*_*i*_, *N*_*ijk*_ is the count corresponding to occurence of the event {*X*_*i*_ = *x*_*ik*_} ∩ {*π*_*i*_ = *w*_*ij*_} in the dataset, and 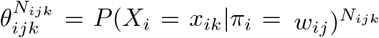 is the conditional probability associated with that event. If the prior density *f* (Θ|*G*) of possible conditional probabilities is assumed to be of the form

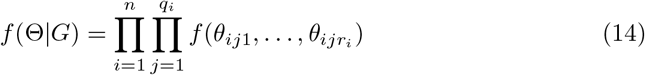

evaluating the *P* (*G, D*) integral with the uniform prior *f* (Θ|*G*) = (*r* − 1)! leads to the expression

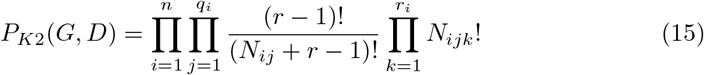

which is known as the K2 score (Cooper and Herskovits 1992). In practice, the actual score is taken to be the natural log of the above expression to simplify computation of nested products and factorials, yielding

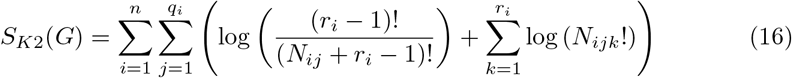

The Dirichlet prior *f* (Θ|*G*) ∼ **Dir**(*α*_*ijk*_) with strictly positive hyper-parameters *α*_*ijk*_ which depend on *G* gives rise to a more general expression for *P* (*G, D*) known as the BD score (see Heckerman et al. (1995)):

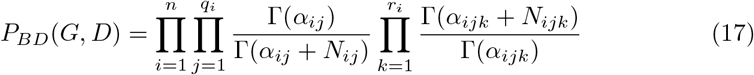

where *α*_*ij*_ = ∑_*k*_ *α*_*ijk*_. This formula yields K2 when *α*_*ijk*_ = 1, and the *Bayesian Dirichlet equivalent-uniform* score (BDeu) when *α*_*ijk*_ = *α/*(*r*_*i*_*q*_*i*_) for some *α* (e.g. *α* = 1).

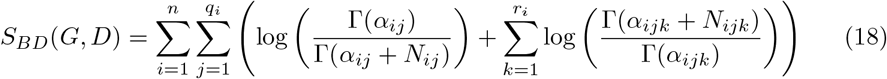

It is possible to establish a relationship between the BD score and information-theoretic entropy via the asymptotic expansion of log (Γ(*z* + 1)) with Stirling’s series (Arfken and Weber 1995):

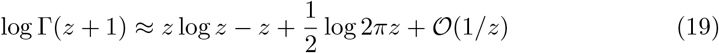

Factoring the terms of *S*_*BD*_ that contain *N*_*ijk*_ and *N*_*ij*_

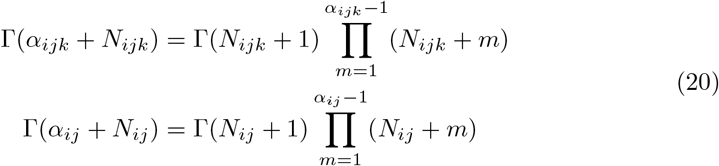

and expanding the resulting log(Γ(*N*_*ijk*_ + 1)) and log(Γ(*N*_*ij*_ + 1)) factors with Stirling’s series leads to the following approximation:

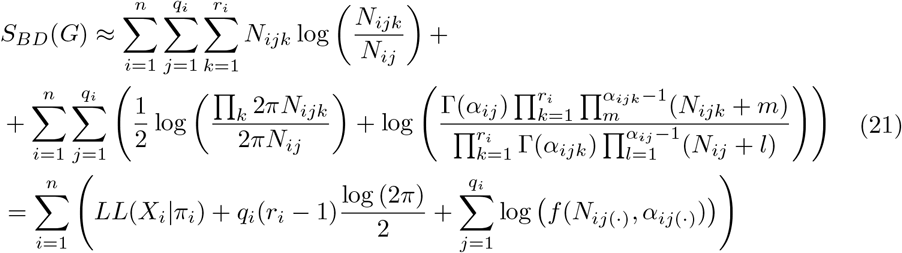

where *LL* is the log-likelihood and *f* is a function that depends on the prior and the joint counts. Isolating *LL* and collecting the rest of the summands into *C*_*i*_ yields the expression in terms of entropy:

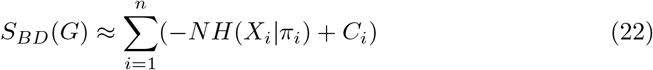

Hence, the condition for accepting a test structure *G*^*′*^ is, once again, that there is at least one *i* such that

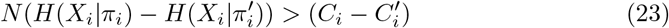

Clearly, the difference between the condition imposed by BD and the one imposed by MDL lies primarily in the structure of the penalty term Δ*C*_*i*_. Otherwise, just like with MDL, model selection based on BD can be interpreted as a decision-theoretic matter of recognizing conditional independence when Δ*H*_*i*_ is below the threshold given by Δ*C*_*i*_.

### 2.2 MDL Penalty Miscalibration

Let *X, Y, Z* be a triple of 4-variate random variables related by *Y* → *X* ← *Z*. Here we will demonstrate the scenario where MDL fails to recognize the *X* ← *Z* edge, i.e. the dependence of *X* on *Z* given *Y* . Consider a dataset where the joint events {*X* = *x*_*i*_}∩ {*Y* = *y*_*j*_} ∩ {*Z* = *z*_*j*_} correspond to the empirical counts *N*_*ijk*_ with the respective indices *i, j, k* = 1, …, 4 of the instantiations of the variables. Suppose the sample size is *N* = 64 and let these empirical counts be given by

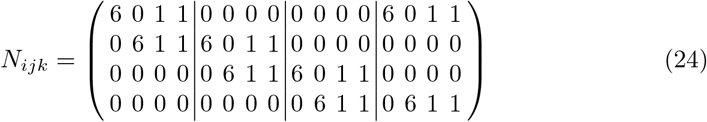

where the vertical lines represent the transition in the *k* index and clearly show the dependence on *Z*. Marginalization over *i* (the X index) gives the counts for *P* (*Y, Z*):

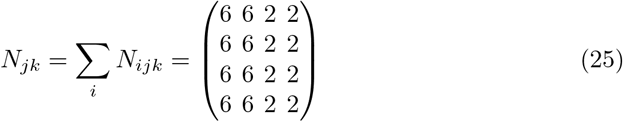

Marginalization over *k* (the Z index) gives the counts for *P* (*X, Y*):

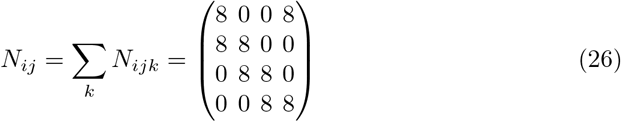

Then the respective counts for *P* (*X*), *P* (*Y*), *P* (*Z*) are given by

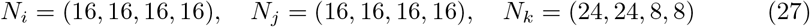

To demonstrate that MDL rejects *X* ← *Z* edge it is sufficient to evaluate MDL introduced in (5) for the structure with and without the edge, i.e. for *π* = *Y* and *π*^*′*^ = *Y* ∩ *Z*.

Evaluating the entropy term for *π* yields

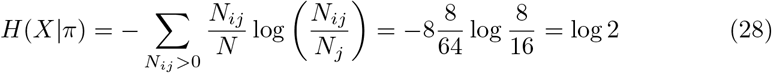

Hence, the sought score of the structure without *X* ← *Z* is

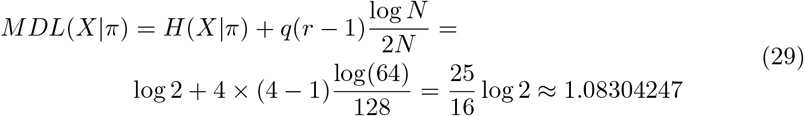

For *π*^*′*^ the entropy terms is

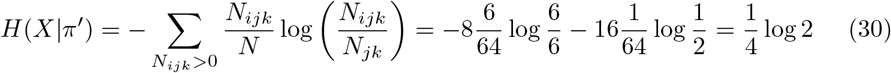

Therefore, the score of the structure with *X* ← *Z* edge

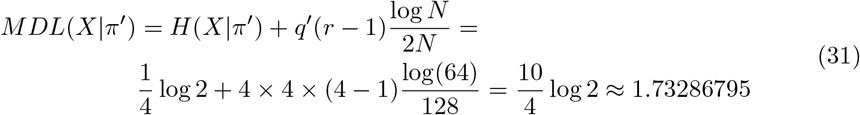

where *q*^*′*^ = 4 *×* 4 is the cardinality of *π*^*′*^.

Because *MDL*(*X*|*π*) *< MDL*(*X*|*π*^*′*^), the parent set *π*^*′*^ is rejected. In other words, the strong dependency *X* ← *Z* that clearly appears in *N*_*ijk*_ is completely disregarded by MDL with the absolute certainty of a massive margin between in the score values.

For reference, interrogating the same configuration with the K2 and BDeu with *alpha* = 20 yields

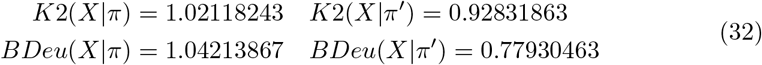

showing no signs of struggle in recognizing the edge.

Doubling the sample size to *N* = 2 *×* 64 does not resolve the issue, rejecting *π*^*′*^, albeit with a smaller margin.

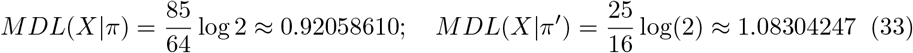

### 2.3 Minimum Uncertainty (MU) Principle

A proper BN model is, as a rule, significantly sparser than the fully entangled network due to conditional independence (CI) constraints. One of the core objectives of BNSL is to identify CI relationships that allow efficient factorization of the underlying distribution and constrain the inherently ill-posed problem. Functionally, BNSL is the process of discriminatively disentangling a minimal sufficient factorization structure of the underlying distribution from the background of redundancy. CI stands out as the sole BN model specification that determines the quality of this disentanglement.

The primary challenge in recognizing CI lies in the numerical satisfiability of the independence criterion. Whatever form the idealized independence criterion takes, its numerical realization always has limited precision. Inferring CI from observations, compiled in the finite, often insufficient, sample size dataset, is always subject to sampling error. This is further compounded by the inherent uncertainty in the correspondence between the observations and the source distribution, not only due to non-uniqueness of the distribution able to produce the same observations, but also due to the variation in the observations arising from the same distribution.

It follows that, if the true distribution is unknown, the empirical realization of the CI criterion can never be numerically satisfied with certainty. There is always a possibility that Δ*H*_*i*_ = 0 does not actually mean independence, and, conversely, that Δ*H*_*i*_ *>* 0 actually corresponds to independence because of the error or uncertainty in probability estimates. This means that to avoid performance irregularities, incongruence, or inconsistent behavior, a scoring function needs to integrate a mechanism that can adequately differentiate between dependence and independence scenarios of the CI decision problem. It also explains the irregularities observed with MDL and BD scoring whose penalizing mechanisms are not specifically designed to properly recognize CI despite the apparent algorithmic demand (Section 2.1).

Motivated by this reasoning, we formulate BNSL in terms of minimization of uncertainty in the empirical realization of the fundamental model specification (i.e. empirically consistent CI topology), establishing what we refer to as the *Minimum Uncertainty* model selection principle. From this principle we derive the preliminary criterion that integrates the CI decision task into its penalizing mechanism by design.

In order for the penalty to be congruent with the independence criterion it must be derived from the same principles, otherwise competing goals lead to an overall incoherent behavior. The most straightforward way to address this is to relate the threshold Δ*C*_*i*_ to the variation in the criterion Δ*H*_*i*_ due to the variation of its argument.

This can be achieved by approximating the conditional entropy *H*(*X*|*Y*) in terms of the unconditional entropy *H*(*X*) and a residual term, i.e. *H*(*X*|*Y*) ≈ *H*(*X*) + *R*, allowing the derivation of an upper bound *µ* in *H*(*X*) − *H*(*X*|*Y*) *< µ* for the near-conditional-independence scenario where small variation in parameters can be attributed to sampling error or uncertainty.

Let ***h*** be a perturbation of a member ***p*** of a (*r* −1)-simplex *S*, such that 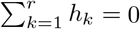 0, so that ***p*** + ***h*** ∈ *S*. Define entropy as 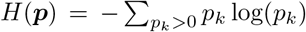 to extend its domain to the simplex boundary. Since *H* is analytic over the interior of *S* it is equal to its Taylor expansion

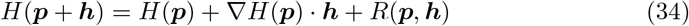

where *R*(***p, h***) represents higher order terms. Hence, for two *r*-variate random variables *X* and *Y* the CI criterion becomes

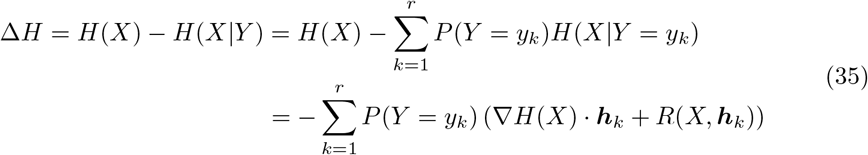

where ***h***_*k*_ is a perturbation to the distribution of *X* due to conditioning on events {*Y* = *y*_*k*_}.

Let ***p***_*X*_ ∈ *S* be the density of *X* and *z*_*k*_ be a probability estimate for the smallest observable event in the data conditioned on {*Y* = *y*_*k*_}, then

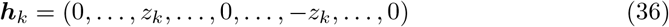

with nonzero components *h*_*ki*_ = *z*_*k*_ and *h*_*kj*_ = −*z*_*k*_ is the smallest observable perturbation that satisfies 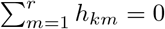 and ***p***_*X*_ + ***h***_*k*_ ∈ *S*.

Approximation of the conditional entropy *H*(*X*|*Y* = *y*_*k*_) in (35) with an expansion in terms of the components of ***p***_*X*_ and ***h***_*k*_ yields

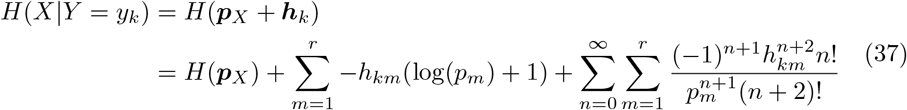

Determining upper and lower bounds for each term of the expansion of *H*(*X*|*Y* = *y*_*k*_) leads to the following inequalities (Gogoshin and Rodin 2025):

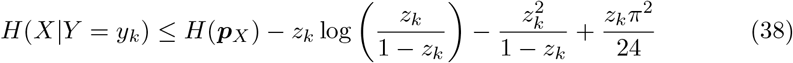

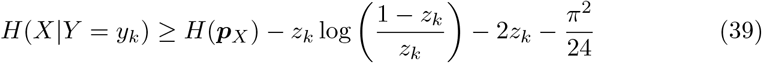

Hence, the upper bound for Δ*H* is

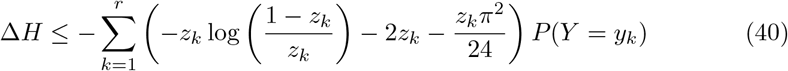

Using maximum likelihood probability estimates *z*_*k*_ = 1*/N*_*k*_ and *P* (*Y* = *y*_*k*_) = *N*_*k*_*/N* where *N*_*k*_ is the sample count for {*Y* = *y*_*k*_} yields the computable bound *µ*(*Y*) that depends only on the structure of conditioning events

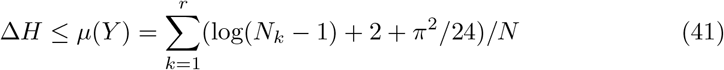

with *N*_*k*_ *>* 1. Since in the event {*Y* = *y*_*k*_} with a very small sample count such as *N*_*k*_ = 1 the sampling error dominates everything else, the term log(*N*_*k*_ − 1) can be safely changed to log(*N*_*k*_). Generalizing *µ* to several *r*_*i*_-variate conditioning variables *Y*_*i*_ gives

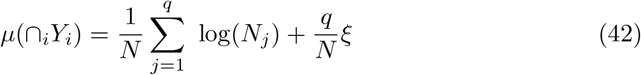

where *q* = П_*i*_ *r*_*i*_ is the number and *N*_*j*_ is the sample count of the joint events of ∩_*i*_*Y*_*i*_ and *ξ* = 2 + *π*^2^*/*24.

The model selection criterion for a test structure *G*^*′*^ that modifies the parent set *π*_*i*_ of *X*_*i*_ to 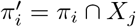 can then be stated as

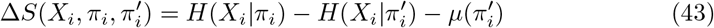

where the condition for accepting *G*^*′*^ is that there is an edge *X*_*i*_ ← *X*_*j*_ such that *S*(*X*_*i*_, *X*_*j*_) *>* 0, and *µ* serves the purpose of penalizing statistical uncertainty in near-conditional-independence scenario. Note that the penalty for conditioning *X*_*i*_ on the set 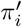 is the same as that for modifying *π*_*i*_ to 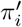 above. This is because parent set is just a joint distribution of the parent variables. Hence, the score can be defined for an arbitrary parent set of *X*_*i*_ as

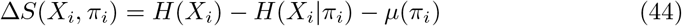

The optimal parent set of *X*_*i*_ must satisfy

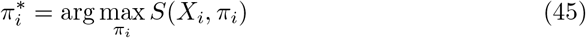

which is equivalent to

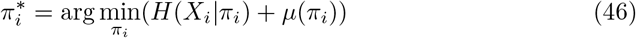

Therefore, *S*(*X*_*i*_, *π*_*i*_) can be redefined as

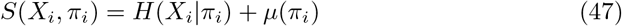

Where 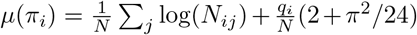, with *N*_*ij*_ being the parent set *π*_*i*_ counts and *q*_*i*_ being its cardinality. This is the form of the scoring criterion derived in Gogoshin and Rodin (2025).

### 2.4 Toward a Resolution-Aware, Calibrated MU Criterion

The scoring criterion detailed in the previous section was obtained in the course of ascertaining whether it is, in principle, possible to use such an approach for model selection. The penalty used for this purpose in Gogoshin and Rodin (2025) was derived as a rough upper bound and, as such, has limitations.

First, the proximity of the penalty to the actual range of Δ*H* was not established, so it is unclear whether this penalty could be more discriminating or, perhaps, more relaxed.

Second, this penalty is a function of parent configuration only and does not account for the cardinality of the child variable. This implies substantive applicability limitations since the resolution of empirical conditional probability estimates should be a function of not only the size of the conditioning events but also of the size or number of the joint events embedded within the conditioning event, depending on the parametrization. The latter point is fairly evident given that for any fixed sample count of a conditioning (parent) event {*Y* = *y*_*k*_} of *X*, increasing cardinality *r* of *X* leads to higher fragmentation of the conditional probability *P* (*X*|*Y* = *y*_*k*_) with a necessarily decreasing sample count of the conditional events {*X* = *x*_*j*_|*Y* = *y*_*k*_} and a subsequent loss of resolution.

Third, interrogating the test problem defined in Section 2.2 with the criterion *S* defined in (47), using the previously obtained quantities *H*(*X*|*π*) = log 2, 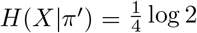 and 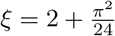, yields

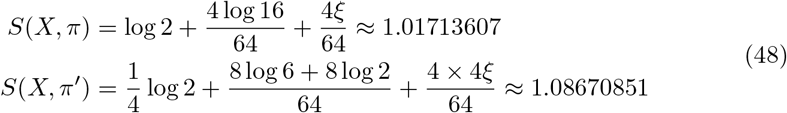

which, in line with MDL, incorrectly rejects *X* ← *Z* since *S*(*X, π*) *< S*(*X, π*^*′*^). However, unlike for MDL, doubling the sample size *N* = 2 *×* 64 is sufficient to improve the sensitivity profile and accept *π*^*′*^, correctly identifying the dependence.

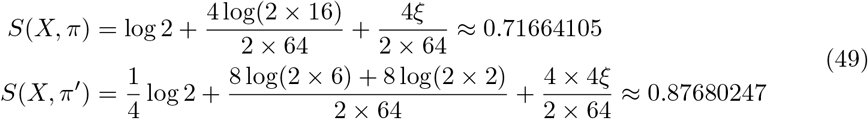

The two examples demonstrate that although we substantially improved upon MDL, significantly narrowing the excessive rejection gap, the penalty still needs adjustment in order to better align its sensitivity profile with the reality.

The unaccounted loss of resolution and the sensitivity adjustment will be addressed in the next section by an appropriately parametrized and a more direct bound derivation. Then the results will be assessed (and compared to other scores) via binary decision metrics.

### 2.5 The New Model Uncertainty Bounds

Again, suppose ***p***_*X*_ is a member of (*r* − 1)-simplex *S* where *z*_*k*_ = 1*/N*_*k*_ is the empirical probability estimate for the smallest observable event in the data conditioned on the *k*-th state {*Y* = *y*_*k*_} of a *q*-state variable *Y* . Due to convexity and symmetry of entropy over the interior of the simplex the vertex neighborhoods of the simplex must correspond to the greatest variation in *H* for a given perturbation. Hence, a generic upper bound can be derived by considering a test region with ***p***_*X*_ located in the immediate neighborhood of one of the vertices

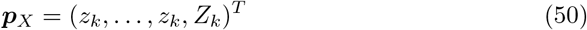

with *Z*_*k*_ = 1 − (*r* − 1)*z*_*k*_ and a perturbation ***h***_*k*_ ∈ *S* with and an even number of nonzero components equally split between *w*_*k*_ positive and *w*_*k*_ negative instantiations,

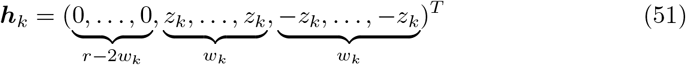

so that **1**^*T*^ · ***h***_*k*_ = 0. Equation (35) then takes the form:

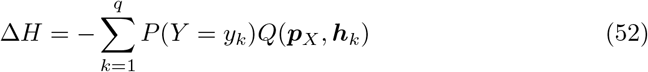

where the remaining term of the expansion *Q*(***p***_*X*_, ***h***_*k*_) are given by

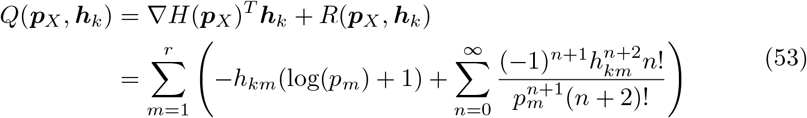

with *h*_*km*_ and *p*_*m*_ being the respective components of ***h***_*k*_ and ***p***_*X*_ . Evaluating Equation (53) with the current configuration of ***h***_*k*_ yields

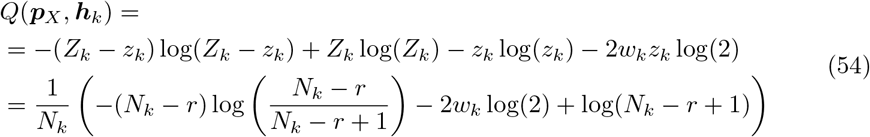

provided that |*Z*_*k*_| ≥ |*z*_*k*_| which can also be expressed as |*N*_*k*_ − *r* + 1| ≥ 1.

Constraining the admissible parameters by *N*_*k*_ ≥ *r* restricts the expansion to the interior of the simplex. When *N*_*k*_ = *r* the estimation region lies at the center of the simplex, while for *N*_*k*_ *< r* the test location always appears on its boundary. In the latter case, at least the *Z*_*k*_ component of ***p***_*X*_ is zero and the expansion must be restricted to the interior of a simplex of a lower dimension to remain coherent. This situation is addressed with the last bound at the end of this section.

Let *T*_*k*_ = *N*_*k*_ − *r* + 1 in Equation (54) to simplify the notation. Substituting *Q* into Equation (52) and evaluating the result with ***h***_*k*_ in (51) leads to

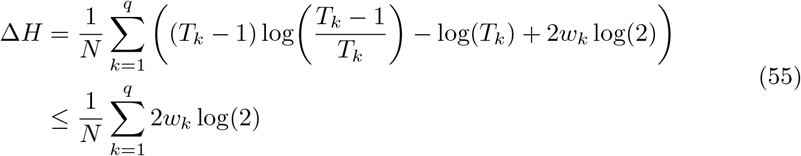

The expression under the sum has negative derivative over the admissible domain and, therefore, attains its maximum at *T*_*k*_ = 1, yielding the bound.

Permuting the positive and the negative components of ***h***_*k*_ so that

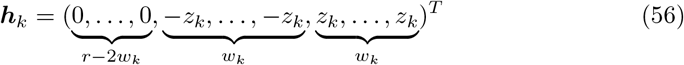

and re-evaluating Equation (53) with subsequent substitution of the result into Equation (52) yields the following:

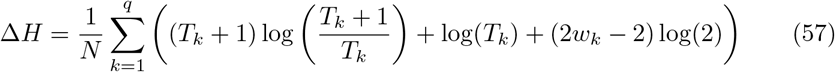

provided |*T*_*k*_| ≥ 1. Since the expression under the sum has a positive derivative over the admissible range of *T*_*k*_, its minimum is attained at *T*_*k*_ = 1, which implies strict positivity for any choice of parameters.

Further, recognizing that (*T*_*k*_ + 1) log((*T*_*k*_ + 1)*/T*_*k*_) ≤ 2 log(2) and that log(*T*_*k*_) ≤ log(*N*_*k*_) yields a much simpler and more general bound:

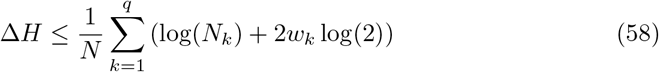

Finally, permuting ***h***_*k*_ so that zero appears in the last component

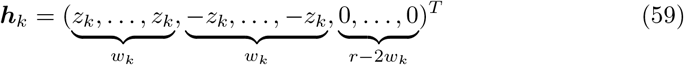

yields an expression containing only *w*_*k*_ terms identical to what was already obtained in (55):

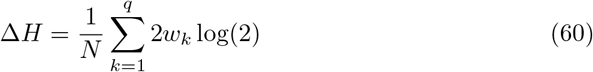

This bound also addresses the degenerate regime of the expansion (*N*_*k*_ *< r*) where at least the *Z*_*k*_ component of ***p***_*X*_ is zero.

The bounds obtained thus far are dominated by (58) for any choice of parameters. In fact, the restriction of (58) to the case *N*_*k*_ = 1 reproduces the other bounds. It follows that this result can be used as a generic upper bound for the considered parametrization of the perturbation neighborhood.

### 2.6 The New Calibrated MU Criterion and Its Interpretation

In this section, the result obtained in Equation (58) which still depends on the unknown perturbation parameter *w*_*k*_ is converted into a penalty fit for practical applications. One of the ways to eliminate *w*_*k*_ is to assume that it is the outcome of a uniformly distributed random variable *W* with 0 ≤ *w*_*k*_ ≤ *r/*2, and whose expectation value therefore satisfies *E*(*W*) = 1*/q* ∑_*k*_ *w*_*k*_ = *r/*4. This leads to further simplification and generalization of the bound

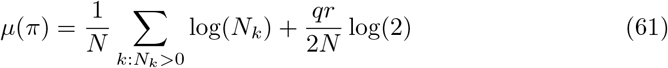

where *q* is now the number of non-zero counts *N*_*k*_ *>* 0 of *π*. A remarkable fact is that the structure of (61), although much simpler, is still surprisingly similar to the BD penalty *C*_*i*_ obtained in (21), despite their vastly different derivation methods. On the other hand, MDL penalty is completely missing the term that depends on the actual counts, which explains its failure to scale properly as will be demonstrated in Section 3.1.

Investigating the asymptotic form of the derived cost function yields 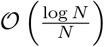 as can be seen from

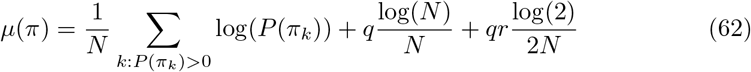

where *π*_*k*_ are individual instantiations of *π* and *q* = card {*π*_*k*_ ∈ *π* : *P* (*π*_*k*_) *>* 0}.

Since for any variable the cost *µ* is a function of its parent configuration, we can define empirical *certainty* of *X*_*i*_ due its parent set *π*_*i*_ as

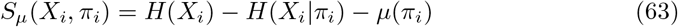

so that the information gain due to *π*_*i*_ must overcome the sampling resolution uncertainty in order for the structure to be considered well supported by evidence. If the optimal *π*_*i*_ is understood as the parent set with the best numerical evidence (i.e. maximum certainty)

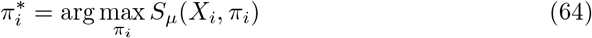

the underlying optimality criterion actually seeks the parent set with the least *uncertainty*

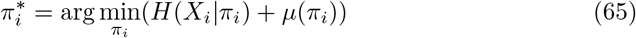

Hence, in this framework, the minimized objective function represents the measure of *model uncertainty* and the associated MU criterion for a graph *G* with *n*-nodes *X*_*i*_ is given by

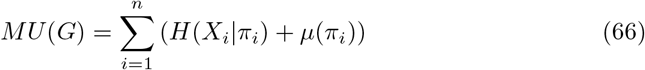

with *µ* defined in (61) and its asymptotic variant defined in (62).

Interrogating the problem defined in Section 2.2 with MU, using the previously obtained quantities *H*(*X*|*π*) = log 2 and 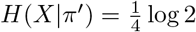, yields

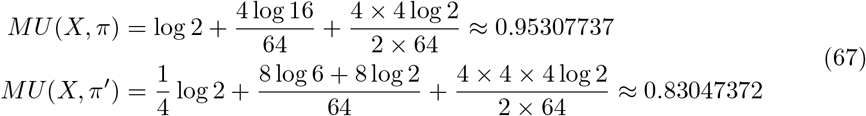

which gives *MU* (*X, π*) *> MU* (*X, π*^*′*^) and implies that the edge *X* → *Z*, discarded by MDL, is successfully accepted by the calibrated MU criterion with a sensible margin, comparable to what was seen with K2 and BDeu. The *minimum uncertainty* principle underlying the calibrated MU criterion achieves this by prioritizing multiscale statistical resolution (evidence quality), increasing rejection rate when it drops. In plain language, it amounts to the idea that, all other things being equal, the best explanations (structural elements) are to be inferred from the most internally coherent and unambiguous observations, refraining from speculative conclusions where evidence is lacking.

### 2.7 Discriminative Power Evaluation Methodology

A way of assessing the ability of a scoring criterion to correctly identify conditional independence (CI) relationships is to estimate its sensitivity and specificity profiles or, equivalently, its type I and type II error rates. For this purpose, the classical decision-theoretic framework requires two distributions that correspond to CI and its alternative, conditional *dependence*.

The statistical model needed for this inquiry must be specified by the joint probability distribution *P* (*X, Y, π*) whose internal structure either admits CI factorization or, as its disjoint falsifiable alternative, is not factorizable in this way. Framing this more precisely in terms of entropy, either *H*(*X, Y, π*) = *H*(*X*|*π, Y*) + *H*(*Y, π*), or *H*(*X, Y, π*) = *H*(*X*|*π*) + *H*(*Y, π*) where *X* and *Y* are individual variables and *π* is a possibly empty set of conditioning variables.

Note that the two disjoint alternatives can be readily reduced to the already familiar equivalent requirement that ether Δ*H* = 0 or Δ*H >* 0 where Δ*H* = *H*(*X*|*π*) − *H*(*X*|*π, Y*).

Assuming *π* is not empty, the described conditional relationship between *X* and *Y* can be modeled with networks in Figure 1 that correspond to three possible factorizations of the *H*(*X, π*) term. Each model configuration, denoted as mode A, mode B, and mode C is a pair of structures *M*_0_ and *M*_1_ where *Y* → *X* is either present or absent and correspond to the CI hypothesis and its alternative respectively.

**Fig. 1.**
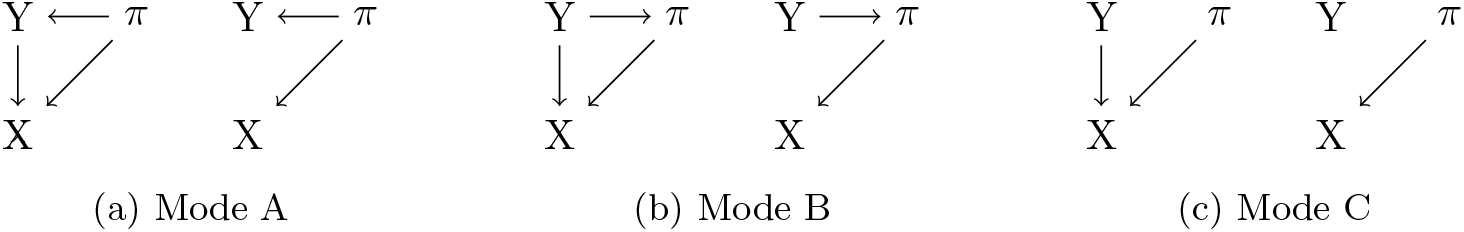
The test alternatives *M*_0_ (on the left) and *M*_1_ (on the right) for modes A, B, C.

In Figure 1a, in the absence of *Y* → *X* edge, *X* is independent of *Y* given *π*. The corresponding model specification is given by the following factorizations of the joint probability

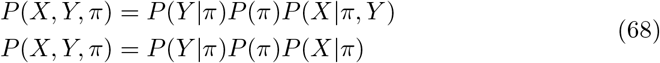

The dependency configuration of mode B (Fig. 1b) is equivalent to mode A as a whole. Here, again, in the absence of *Y* → *X* edge, *X* is independent of *Y* given *π*. Locally, however, whether *Y* or *π* is defined as a root node can impact the distribution of *H*(*X*|*π*) − *H*(*X*|*π, Y*).

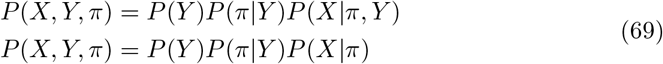

The third scenario (Fig. 1c) captures unconditional independence that arises when *P* (*Y, π*) = *P* (*Y*)*P* (*π*) and corresponds to the following factorizations:

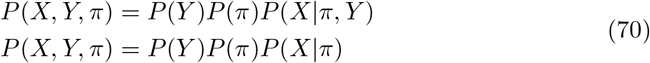

The testing procedure must assess the ability of a given scoring criterion to discriminate between dependence and independence configurations. To gather information needed for discriminative power evaluation, the procedure must sample the space of distributions for a set of fixed parameters corresponding to the particular similarity class of relationships.

For the scenario A (Fig. 1a) let *P* (*π*) be a (*r*_*π*_)-dimensional vector, *P* (*Y* |*π* = *π*_*k*_) a (*r*_*Y*_)-dimensional vector and *P* (*X*|*π* = *π*_*k*_, *Y* = *y*_*j*_) a (*r*_*X*_)-dimensional vector, each sampled from an uninformative Dirichlet distribution with appropriate parameters, so that dim(*P* (*X*|*Y*)) = *r*_*π*_ *× r*_*Y*_ and dim(*P* (*X*|*Y, π*)) = *r*_*Y*_ *× r*_*π*_ *× r*_*X*_ . To make sure that the generated structure replicates dependency relationships it is sufficient that at least a pair of vectors of each conditional probability are linearly independent (Gogoshin et al. 2021).

The product of these factors forms the joint distribution 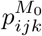 replicating the desired factorization property of the first equation in (68) to model dependency

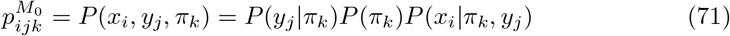

Marginalizing the joint distribution over *y*_*j*_ generates *P* (*x*_*i*_|*π*_*k*_) = ∑_*j*_ *P* (*x*_*i*_, *y*_*j*_, *π*_*k*_)*/P* (*π*_*k*_) required to form the distribution 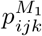, the second equation of (68) that models CI

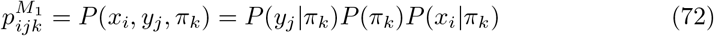

For the scenario B in Figure 1b the procedure is very similar with *Y* and *π* reversing the roles in that now *P* (*Y*) is a (*r*_*Y*_)-dimensional and *P* (*π*|*Y* = *y*_*j*_) is sampled as a (*r*_*π*_)-dimensional vector so that dim(*P* (*π*|*Y*)) = *r*_*Y*_ *× r*_*π*_. *P* (*X*|*Y, π*), obtained in the way identical to model A, is then used to generate the joint distributions that appear in (69) for *M*_0_ and *M*_1_ respectively.

Finally, the scenario C (Fig. 1c) differs from B only in that *P* (*π*|*Y* = *y*_*j*_) = *P* (*π*) for all *j* and so only demands one (*r*_*π*_)-dimensional vector to be sampled. The other quantities needed for the joint distributions of (70) are obtained in the same way as in B.

For each mode the joint distribution pair *M*_0_ and *M*_1_ can now be used to simulate the effect of sampling error for a given sample size *N* . Generating categorical counts *C* = {*N*_*ijk*_} for the joint states *N*_*ijk*_ from a multinomial distribution **Mlt** to obtain 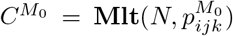 and 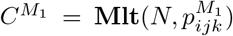 with *M*_0_ and *M*_1_ as its respective conjugate priors yields the desired distributions needed for the binary test of the CI property.

To combine all of the above into one concise assessment, the procedure for evaluating operating characteristics needs an uninformative prior across all parameters of the model, including the mode configurations. Hence, the parameter sampling should span the modes {*A, B, C*} of the model with equal probability. The resulting binary test alternatives *M*_0_ and *M*_1_ for an 8-state node model computed for three different sample sizes *N* can be seen in Figures 2-4. As expected, with increasing sample size the overlap between *M*_0_ and *M*_1_ decreases, which reflects an improvement in resolution of the data.

**Fig. 2.**
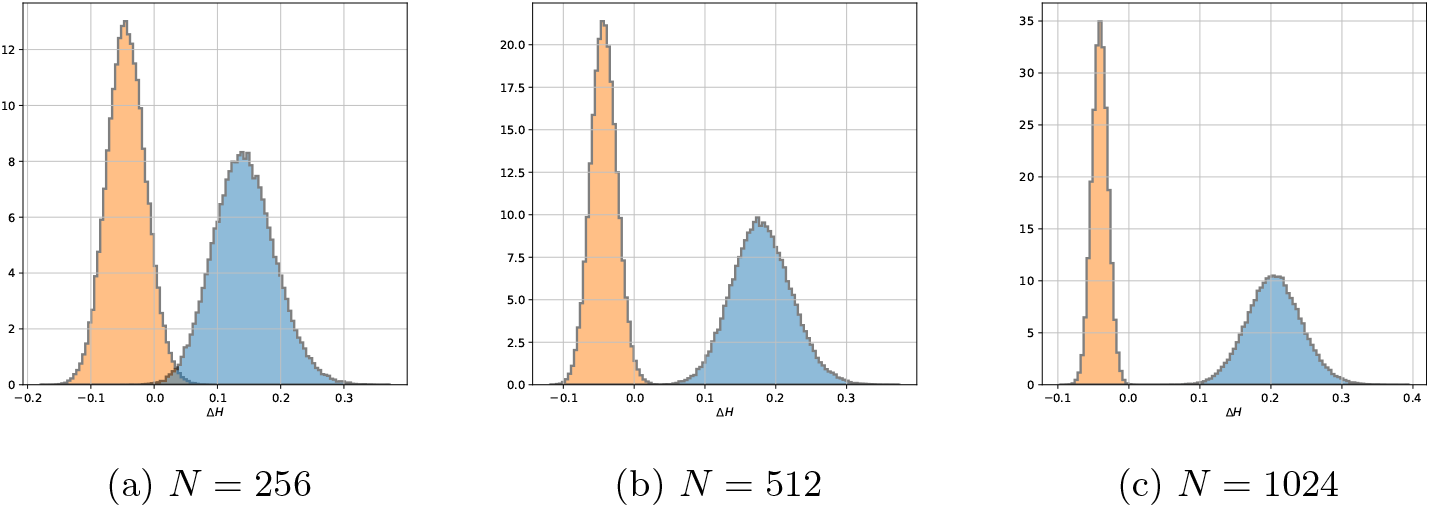
The alternatives *M*_0_ and *M*_1_ with an uninformative mode prior evaluated with the MU score.

**Fig. 3.**
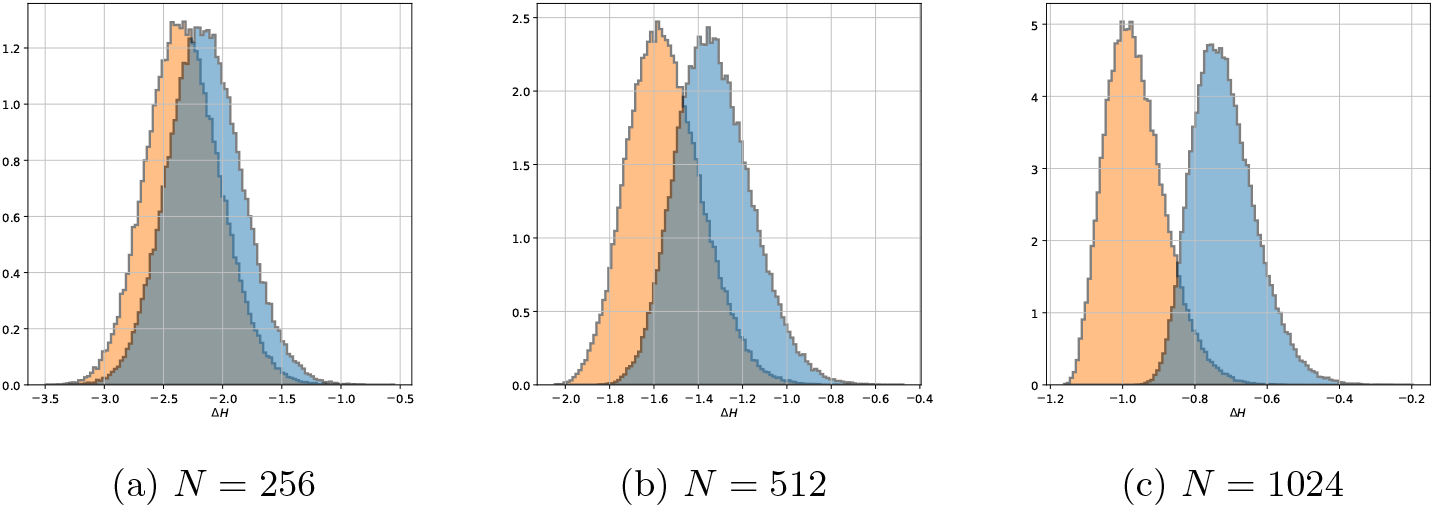
The alternatives *M*_0_ and *M*_1_ with an uninformative mode prior evaluated with the MDL score.

The complete discriminative power profile can now be calculated for every score *S*, considered thus far, as the empirical probability of correctly identifying dependent and independent scenarios. Using the appropriate components of *M*_0_ and *M*_1_ test alternatives, one can obtain distributions of the cost of adding an edge *Y* → *X* via Δ*S* = *S*(*X*|*π*) − *S*(*X*|*π, Y*) (Figures 2-5).

Then the True Negative Rate (TNR) can be obtained as an empirical probability 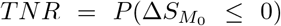 and the True Positive Rate (TPR) is given by 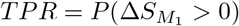. In this work, TNR and TPR will be used interchangeably with specificity and sensitivity, the two terms frequently used in diagnostic binary decision contexts. Note that TPR is also frequently called the *power* of a binary statistical hypothesis test.

Finally, the results obtained as TNR and TPR can be compiled into a single value denominated as accuracy (ACC) via the expression *ACC* = (*TNR* + *TPR*)*/*2. This measure of accuracy is well-ballanced and does not need additional weights because the deision rate evaluation procedure above produces equal number of samples used for TNR and TPR estimation, generating a pair *M*_0_ and *M*_1_ for every sample.

## 3 Results

### 3.1 Discriminative Power Evaluation Results

This section illustrates the application of the discriminative power evaluation framework, developed in Section 2.7, to four scoring functions presented in tables in the order of increasing performance - MDL, BDeu with *α* = 20, K2, and MU which was obtained in Section 2.6.

All sensitivity (TPR) and specificity (TNR) values were calculated via sampling from the Dirichlet prior to obtain *M* 0 and *M* 1 hypotheses, and sampling the joint counts for each hypothesis from the corresponding Multinomial distribution as described in Section 2.7. This decision test simulation was repeated 100000 times for each sample size N. The maximum (worst-case *p* = 0.5) standard error (SE) and the maximum 95% margin of error (MOE) across all calculations are negligible and do not affect the values, presented in the following tables, rounded to two decimal places in any meaningful way.

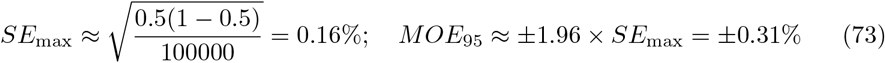

Table 1 compiles the results for models where the conditional distribution is initialized with Jeffrey’s prior, i.e., with *α* = 0.5. The estimation spans cardinality triples (card(*V*) : *V* ∈ (*Y, π, X*)) which appear in the first column. The procedure is carried out for a range of increasing sample sizes (the second column of the table) that captures the variability in all four scores simultaneously. Each scoring function is allocated three rows of the table which correspond to Accuracy (ACC), TNR, and TPR values from left to right.

**Table 1.** ACC (in bold) followed by TNR and TPR for a dependency model with Jeffrey’s prior over a variety of cardinality triples (card(*V*) : *V* ∈ (*Y, π, X*)) appearing in the first column.

| card | N | MDL |  |  | BDeu |  |  | K2 |  |  | MU |  |  |
| --- | --- | --- | --- | --- | --- | --- | --- | --- | --- | --- | --- | --- | --- |
| (3, 3, 3) | 100 | **0.76** | 1.00 | 0.52 | **0.92** | 0.87 | 0.96 | **0.93** | 0.93 | 0.93 | **0.94** | 0.98 | 0.91 |
|  | 200 | **0.87** | 1.00 | 0.75 | **0.96** | 0.93 | 0.99 | **0.97** | 0.97 | 0.97 | **0.98** | 0.99 | 0.96 |
|  | 300 | **0.92** | 1.00 | 0.85 | **0.97** | 0.95 | 0.99 | **0.99** | 0.99 | 0.99 | **0.99** | 1.00 | 0.98 |
|  | 400 | **0.95** | 1.00 | 0.90 | **0.98** | 0.96 | 1.00 | **0.99** | 0.99 | 0.99 | **0.99** | 1.00 | 0.99 |
|  | 500 | **0.96** | 1.00 | 0.93 | **0.98** | 0.97 | 1.00 | **0.99** | 0.99 | 0.99 | **0.99** | 1.00 | 0.99 |
|  | 600 | **0.97** | 1.00 | 0.94 | **0.99** | 0.98 | 1.00 | **1.00** | 1.00 | 1.00 | **1.00** | 1.00 | 0.99 |
| (5, 5, 5) | 100 | **0.50** | 1.00 | 0.00 | **0.93** | 1.00 | 0.86 | **0.95** | 0.91 | 0.98 | **0.97** | 0.96 | 0.99 |
|  | 200 | **0.50** | 1.00 | 0.01 | **0.99** | 1.00 | 0.97 | **0.99** | 0.98 | 1.00 | **1.00** | 0.99 | 1.00 |
|  | 300 | **0.52** | 1.00 | 0.05 | **1.00** | 1.00 | 0.99 | **1.00** | 1.00 | 1.00 | **1.00** | 1.00 | 1.00 |
|  | 400 | **0.58** | 1.00 | 0.15 | **1.00** | 1.00 | 1.00 | **1.00** | 1.00 | 1.00 | **1.00** | 1.00 | 1.00 |
|  | 500 | **0.66** | 1.00 | 0.33 | **1.00** | 1.00 | 1.00 | **1.00** | 1.00 | 1.00 | **1.00** | 1.00 | 1.00 |
|  | 600 | **0.76** | 1.00 | 0.52 | **1.00** | 1.00 | 1.00 | **1.00** | 1.00 | 1.00 | **1.00** | 1.00 | 1.00 |
| (7, 7, 7) | 100 | **0.50** | 1.00 | 0.00 | **0.70** | 1.00 | 0.41 | **0.91** | 0.82 | 0.99 | **0.96** | 0.99 | 0.94 |
|  | 200 | **0.50** | 1.00 | 0.00 | **0.80** | 1.00 | 0.61 | **0.98** | 0.96 | 1.00 | **1.00** | 0.99 | 1.00 |
|  | 300 | **0.50** | 1.00 | 0.00 | **0.89** | 1.00 | 0.79 | **1.00** | 0.99 | 1.00 | **1.00** | 1.00 | 1.00 |
|  | 400 | **0.50** | 1.00 | 0.00 | **0.95** | 1.00 | 0.90 | **1.00** | 1.00 | 1.00 | **1.00** | 1.00 | 1.00 |
|  | 500 | **0.50** | 1.00 | 0.00 | **0.98** | 1.00 | 0.96 | **1.00** | 1.00 | 1.00 | **1.00** | 1.00 | 1.00 |
|  | 600 | **0.50** | 1.00 | 0.00 | **0.99** | 1.00 | 0.98 | **1.00** | 1.00 | 1.00 | **1.00** | 1.00 | 1.00 |
| (9, 9, 9) | 100 | **0.50** | 1.00 | 0.00 | **0.56** | 1.00 | 0.13 | **0.84** | 0.70 | 0.99 | **0.64** | 1.00 | 0.28 |
|  | 200 | **0.50** | 1.00 | 0.00 | **0.55** | 1.00 | 0.10 | **0.94** | 0.88 | 1.00 | **0.98** | 1.00 | 0.96 |
|  | 300 | **0.50** | 1.00 | 0.00 | **0.57** | 1.00 | 0.14 | **0.98** | 0.96 | 1.00 | **1.00** | 1.00 | 1.00 |
|  | 400 | **0.50** | 1.00 | 0.00 | **0.60** | 1.00 | 0.20 | **1.00** | 0.99 | 1.00 | **1.00** | 1.00 | 1.00 |
|  | 500 | **0.50** | 1.00 | 0.00 | **0.65** | 1.00 | 0.30 | **1.00** | 1.00 | 1.00 | **1.00** | 1.00 | 1.00 |
|  | 600 | **0.50** | 1.00 | 0.00 | **0.70** | 1.00 | 0.41 | **1.00** | 1.00 | 1.00 | **1.00** | 1.00 | 1.00 |

A more complete set of results is presented in Supplement A, Tables S1 and S2 which also extend the investigation to more complex model configurations. Specifically, for Table S2, the fixed parent set *π* is assumed to be comprised of 3 nodes whose joint cardinality is card(*π*) = 3 *×* 2 *×* 3, while card(*X*) and card(*Y*) incrementally increase.

Table 2 repeats the procedure of Table 1 for dependency models initialized with a flat prior. An extended set of results for this setup can be found in Tables S3 and S4 in Supplement A.

**Table 2.** ACC (in bold) followed by TNR and TPR for a dependency model with a flat prior with a variety of cardinality triples (card(*V*) : *V* ∈ (*Y, π, X*)) appearing in the first column

| card | N | MDL |  |  | BDeu |  |  | K2 |  |  | MU |  |  |
| --- | --- | --- | --- | --- | --- | --- | --- | --- | --- | --- | --- | --- | --- |
| (3, 3, 3) | 100 | **0.63** | 1.00 | 0.26 | **0.89** | 0.93 | 0.85 | **0.89** | 0.88 | 0.89 | **0.87** | 0.97 | 0.78 |
|  | 200 | **0.75** | 1.00 | 0.49 | **0.95** | 0.97 | 0.93 | **0.95** | 0.95 | 0.95 | **0.93** | 0.99 | 0.88 |
|  | 300 | **0.82** | 1.00 | 0.64 | **0.97** | 0.98 | 0.96 | **0.97** | 0.98 | 0.97 | **0.96** | 1.00 | 0.92 |
|  | 400 | **0.87** | 1.00 | 0.73 | **0.98** | 0.99 | 0.97 | **0.98** | 0.98 | 0.98 | **0.97** | 1.00 | 0.95 |
|  | 500 | **0.90** | 1.00 | 0.80 | **0.99** | 0.99 | 0.98 | **0.99** | 0.99 | 0.99 | **0.98** | 1.00 | 0.96 |
|  | 600 | **0.92** | 1.00 | 0.84 | **0.99** | 0.99 | 0.99 | **0.99** | 0.99 | 0.99 | **0.99** | 1.00 | 0.97 |
| (5, 5, 5) | 100 | **0.50** | 1.00 | 0.00 | **0.69** | 1.00 | 0.39 | **0.88** | 0.80 | 0.96 | **0.93** | 0.93 | 0.93 |
|  | 200 | **0.50** | 1.00 | 0.00 | **0.79** | 1.00 | 0.58 | **0.97** | 0.95 | 0.99 | **0.99** | 0.99 | 0.99 |
|  | 300 | **0.50** | 1.00 | 0.00 | **0.88** | 1.00 | 0.75 | **0.99** | 0.99 | 1.00 | **1.00** | 1.00 | 1.00 |
|  | 400 | **0.50** | 1.00 | 0.01 | **0.93** | 1.00 | 0.86 | **1.00** | 1.00 | 1.00 | **1.00** | 1.00 | 1.00 |
|  | 500 | **0.51** | 1.00 | 0.02 | **0.96** | 1.00 | 0.92 | **1.00** | 1.00 | 1.00 | **1.00** | 1.00 | 1.00 |
|  | 600 | **0.53** | 1.00 | 0.05 | **0.98** | 1.00 | 0.96 | **1.00** | 1.00 | 1.00 | **1.00** | 1.00 | 1.00 |
| (7, 7, 7) | 100 | **0.50** | 1.00 | 0.00 | **0.52** | 1.00 | 0.04 | **0.79** | 0.60 | 0.98 | **0.87** | 0.96 | 0.78 |
|  | 200 | **0.50** | 1.00 | 0.00 | **0.52** | 1.00 | 0.03 | **0.92** | 0.84 | 1.00 | **0.98** | 0.98 | 0.98 |
|  | 300 | **0.50** | 1.00 | 0.00 | **0.52** | 1.00 | 0.04 | **0.98** | 0.95 | 1.00 | **1.00** | 0.99 | 1.00 |
|  | 400 | **0.50** | 1.00 | 0.00 | **0.53** | 1.00 | 0.07 | **0.99** | 0.99 | 1.00 | **1.00** | 1.00 | 1.00 |
|  | 500 | **0.50** | 1.00 | 0.00 | **0.56** | 1.00 | 0.11 | **1.00** | 1.00 | 1.00 | **1.00** | 1.00 | 1.00 |
|  | 600 | **0.50** | 1.00 | 0.00 | **0.58** | 1.00 | 0.17 | **1.00** | 1.00 | 1.00 | **1.00** | 1.00 | 1.00 |
| (9, 9, 9) | 100 | **0.50** | 1.00 | 0.00 | **0.50** | 1.00 | 0.01 | **0.70** | 0.43 | 0.97 | **0.55** | 1.00 | 0.11 |
|  | 200 | **0.50** | 1.00 | 0.00 | **0.50** | 1.00 | 0.00 | **0.81** | 0.62 | 1.00 | **0.87** | 1.00 | 0.74 |
|  | 300 | **0.50** | 1.00 | 0.00 | **0.50** | 1.00 | 0.00 | **0.90** | 0.80 | 1.00 | **0.99** | 1.00 | 0.98 |
|  | 400 | **0.50** | 1.00 | 0.00 | **0.50** | 1.00 | 0.00 | **0.96** | 0.92 | 1.00 | **1.00** | 1.00 | 1.00 |
|  | 500 | **0.50** | 1.00 | 0.00 | **0.50** | 1.00 | 0.00 | **0.99** | 0.97 | 1.00 | **1.00** | 1.00 | 1.00 |
|  | 600 | **0.50** | 1.00 | 0.00 | **0.50** | 1.00 | 0.00 | **1.00** | 0.99 | 1.00 | **1.00** | 1.00 | 1.00 |

As can be observed from the tables, MDL score undergoes a catastrophic loss of sensitivity with increasing cardinality, and, although somewhat mitigating, increase in the sample size cannot fully compensate. BDeu with *α* = 20 generally performs better, but struggles with specificity at lower cardinalities and tends to lose sensitivity at higher cardinalities, repeating the same behavior on a smaller scale. A lower *α* fares somewhat better at lower cardinalities but is even worse at higher cardinalities (Tables S5 and S6 in Supplement A). K2 score has a robust performance profile across all tests, but tends to be overly optimistic, trading specificity for sensitivity even in more challenging settings where the evidence quality might be lacking, e.g., higher cardinality experiments in all of the tables. Finally, MU performs the same or better than K2 in almost all considered scenarios, either reaching 99% first, or maintaining superior specificity in the few cases where its performance is less decisive (e.g., the lowest considered cardinalities in Table S3).

It is essential that a score treats dependencies of increasing statistical uncertainty with greater scrutiny, favoring specificity in situations where firm grounds for a given relationship cannot be easily established. The sensitivity bias exhibited by K2 is thus an undesirable characteristic which MU overcomes. Furthermore, these results show that MU decisively addressed the problems exhibited by MDL, which was one of the key motivations of this study.

### 3.2 Analysis of the Multiscale Apolipoprotein E Dataset

Here, we evaluate how the proposed scoring function affects BN structure reconstruction quality on real data, using a case study representative of many biomedical applications — with the dimensionality of *p/n* ≈ 0.02, mixed variable types, and varying arities and subsampling regimes.

The dataset originates from a study of variation in the apolipoprotein E (APOE) gene and plasma lipid/lipoprotein levels in 854 European American (EA) donors from Rochester, MN, and 702 African American (AA) donors from Jackson, MS (Gogoshin et al. (2017) and references therein). Of the 30 tracked variables, 10 are plasma lipid/lipoprotein measurements and the remaining 20 are APOE gene single-nucleotide polymorphisms (SNPs). In the figures for this section, SNPs are shown as numeric node labels, whereas the other variables are displayed using self-explanatory abbreviations for lipid/lipoprotein measurements (CHOL, HDL, TRIG, APO E, APO B).

In this analysis, a universal BN is learned from the pooled dataset spanning both EA and AA samples. Continuous variables are max-entropy discretized to prescribed coarseness, yielding discrete variables with a given domain cardinality.

The resulting model is then reevaluated on a 50% random subset, EA subset, and AA subset of the pooled data. The effects of this subsampling are visualized by structural changes in the extracted APOE Markov Blanket (MB) — unsupported by the modified evidence edges are rendered as dotted lines in the corresponding figures.

Finally, the discretization partitioning is refined, and the process is repeated with this new data representation. The refinement process illustrates what the sequences of models may be converging to before degradation due to finite-sample variance noise.

The figures also show three binary decision metrics, appearing next to each edge, which are, from top to bottom: (i) edge specificity, (ii) edge sensitivity, and (iii) the percentage of TPR below the actual edge score. The last number does not play a role in the binary hypothesis test; instead, it can be treated as a normalized relative dependency strength measure in that it estimates where a given edge score appears in the TPR range. The measure is fixed-scale (0-100%) and can be used to directly compare edge strengths within and across the networks under mild statistical assumptions. This discriminative power analysis follows the statistical model developed in Section 2.7 with a fixed *π* ∩ *X* density obtained as an empirical estimate from the data, and model mode distributions generated from *Y* variation alone.

Because in the previous section the results suggest that K2 tends to outperform BDeu in most circumstances, in the rest of this section we omit BDeu and only consider learning characteristics of MDL, K2, and MU scores.

#### 3.2.1 MDL Learning Characteristics

Figure 6 shows MB structures of APOE extracted from the universal BN models learned via MDL criterion (see Supplement B for full BNs). In each case, the continuous variables in the dataset were discretized to prescribed coarseness - (a) 3-state, (b) 5-state, (c) 7-state. For MDL, any further discretization refinement results in the loss of a well established APOE – TRIG dependency, signalling the unacceptable degradation of sensitivity not observed with the other two scores, as will be shown below. The MBs obtained with MDL share the variables with the exception of the first configuration (Figure 6a) which excludes 4075 from the envelope likely due to the variable discretization coarseness.

**Fig. 4.**
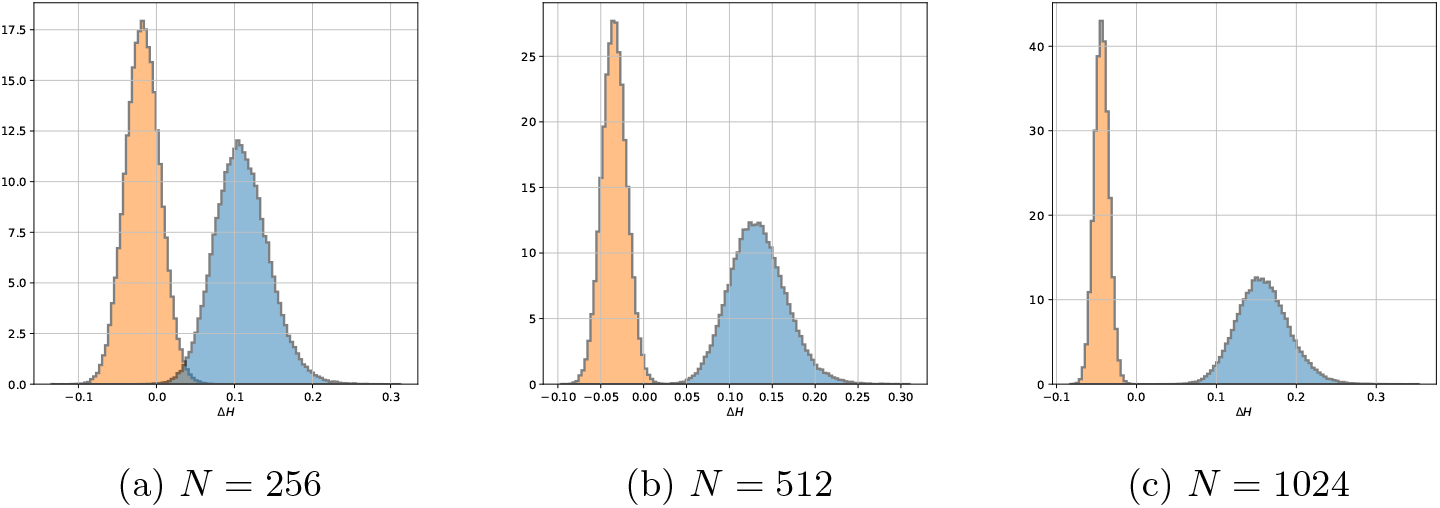
The alternatives *M*_0_ and *M*_1_ with an uninformative mode prior evaluated with the K2 score.

**Fig. 5.**
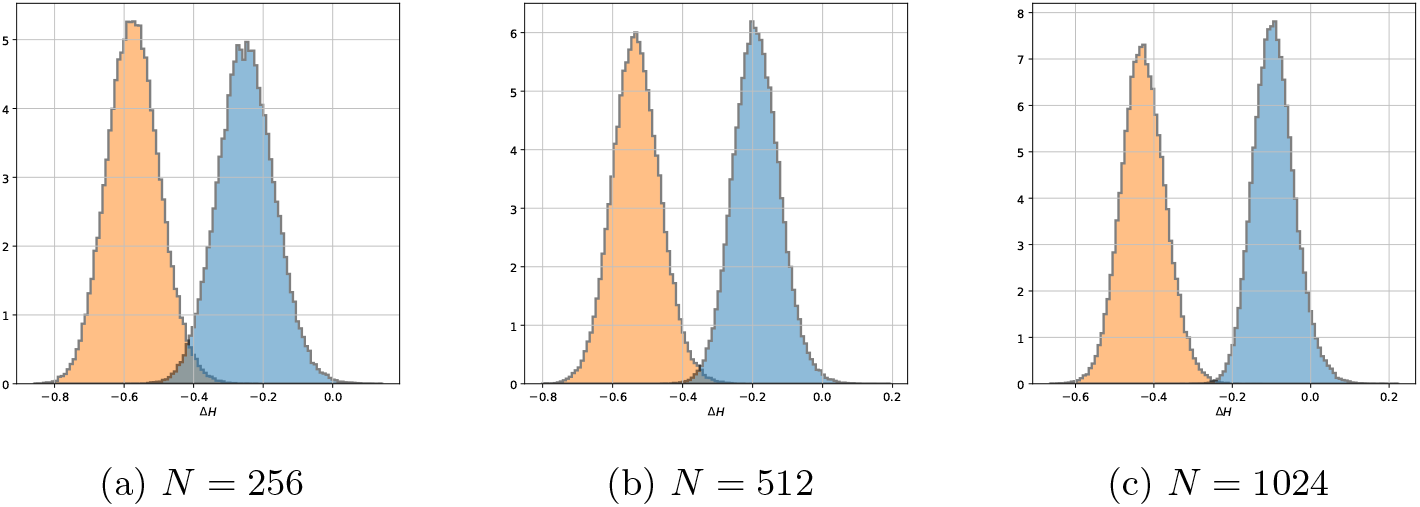
The alternatives *M*_0_ and *M*_1_ with an uninformative mode prior evaluated with the BDeu score.

**Fig. 6.**
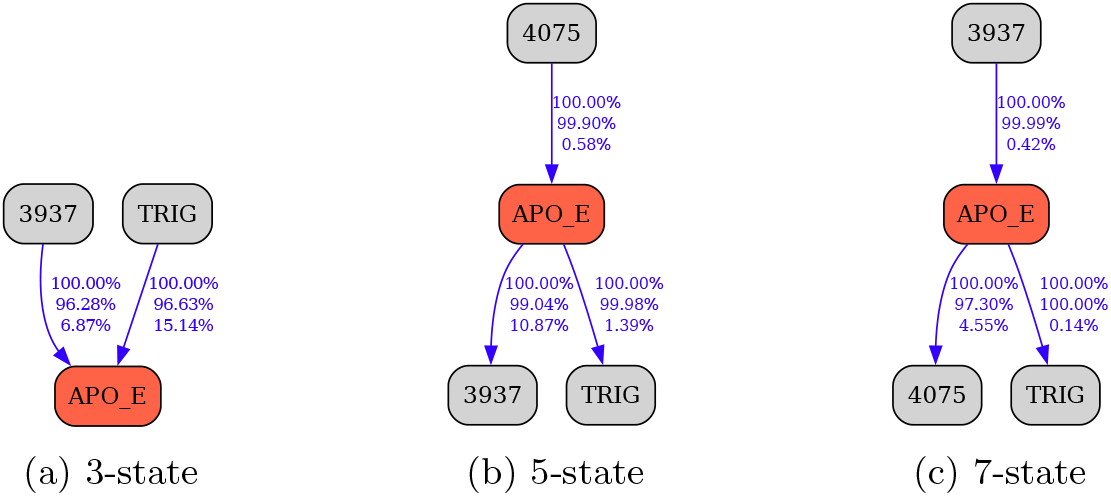
Learning APOE Markov blanket via MDL criterion with data of varying discretization coarseness. The three numbers shown next to each edge are, from top to bottom: edge specificity, edge sensitivity, and the percentage of TPR below the actual edge score which can be interpreted as a normalized edge strength. See text for further details.

The networks shown in Figure 7 correspond to the Figure 6a reevaluated on the 50% random subsample of the complete data, EA subset, and AA subset. The loss of 3937 – APOE edge as a result of AA conditioning (Figure 7c) is unlikely to be attributable to the reduction of the sample size alone, since the loss of this edge is not observed with the random 50% subsampling of the data.

**Fig. 7.**
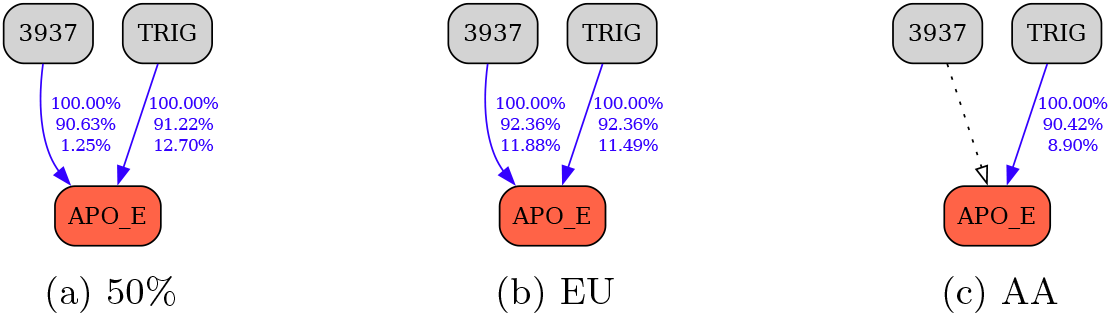
Learning via MDL criterion, 3-state discretization. (a) Random 1/2 subsample. (b) European American subsample. (c) African American subsample.

Figure 8 realizes the same MDL-based analysis for the 5-state discretized data (corresponding to Figure 6b). The dependencies appear to be robust to loss of sampling resolution; AA stratification eliminates APOE – 3937 edge, repeating the previously observed behavior.

**Fig. 8.**
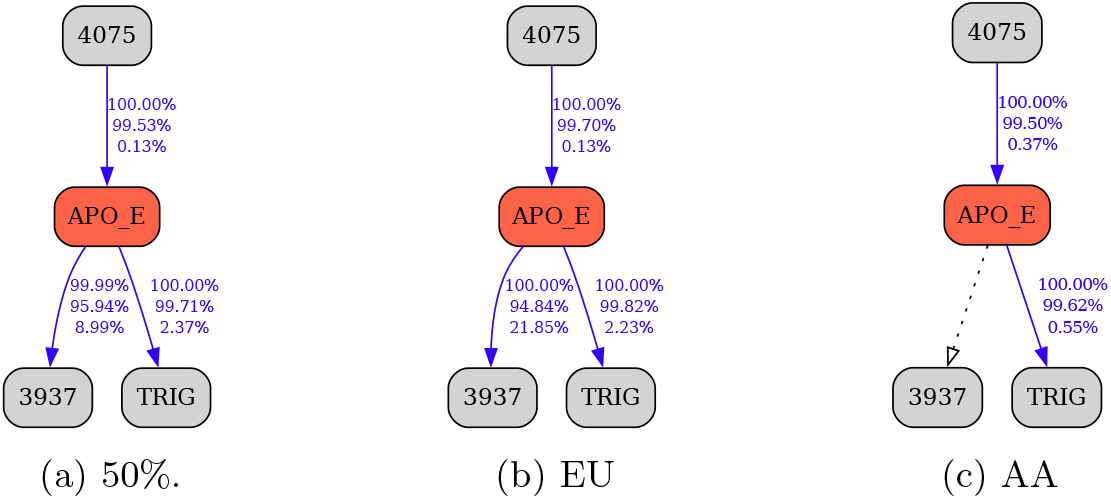
Learning via MDL criterion, 5-state discretization.

Figure 9 carries out this procedure for the 7-state discretized data (corresponding to Figure 6c). Two dependencies, APOE – 4075 and APOE – TRIG, are no longer robust to subsampling. As before, AA stratification eliminates APOE – 3937 dependency in AA conditioned BN (Figure 9c), but now this result has substantially more weight, given the apparent reduction of sensitivity as inferred from the loss of the other two edges, including the well-established TRIG relationship. The loss of sensitivity for MDL is expected in this range of parameters and is coherent with the results of the discriminative power evaluation in the previous subsection.

**Fig. 9.**
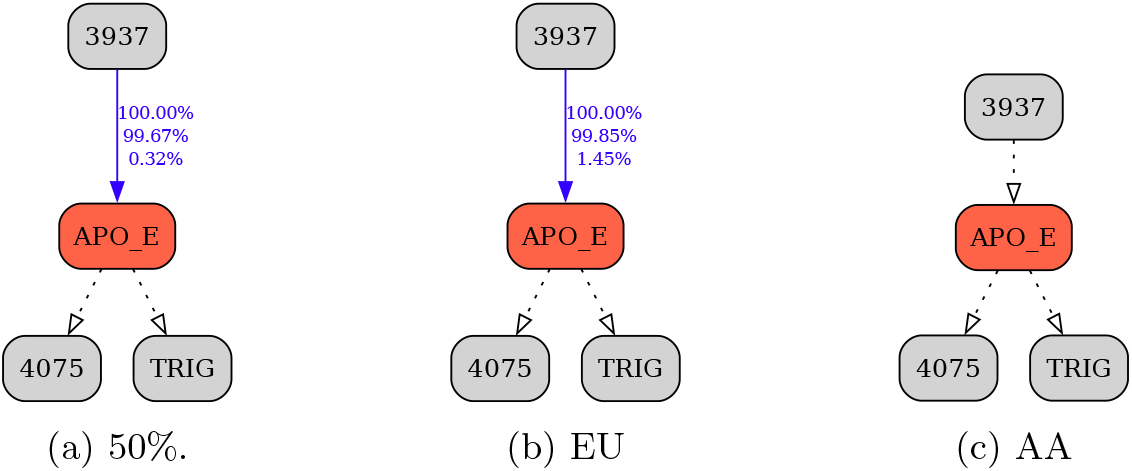
Learning via MDL criterion, 7-state discretization.

#### 3.2.2 K2 Learning Characteristics

Figures 10-12 repeat the above analysis using the K2 score learning, extracting Markov blankets from full BNs for each discretization coarseness (see Supplement B for full BNs). All of the obtained MBs in Figure 10 share the nodes {3937, 4036, 4075, TRIG} while {CHOL, HDL, APOB, 545} appear to be less certain (shared only between two out of three BNs) and may be artifacts due to discretization. This can be explained by K2-score having a bias towards sensitivity as was observed in Subsection 3.1.

**Fig. 10.**
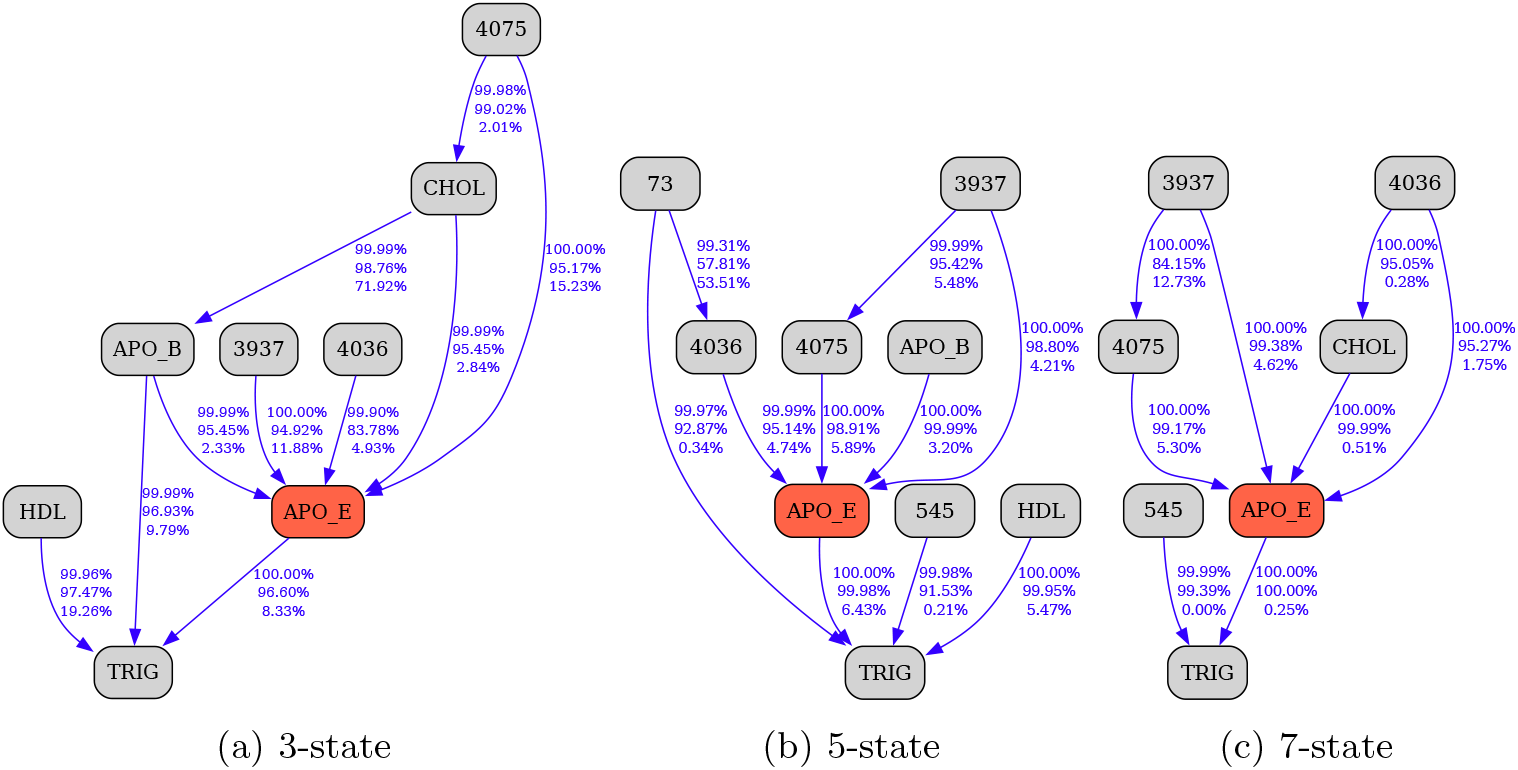
Learning APOE Markov blanket via K2 criterion with data of varying discretization coarsenes.

Using the stratified data to reevaluate the model in Figure 10a reveals the same loss pattern for the 3937 – APOE edge as was observed with the MDL-derived results (Figure 11). Another detail is the elimination of 4036 – APOE due to the EA conditioning. This occurs because 4036 is a rare variant present only in the AA group (25/702 samples) in this dataset, which can serve as the sensitivity check for the procedure instead of TRIG. Both the CHOL – APOE and the APOB – APOE dependencies appear to be unstable with respect to 50% random subsampling, likely due to coarse-grained discretization.

**Fig. 11.**
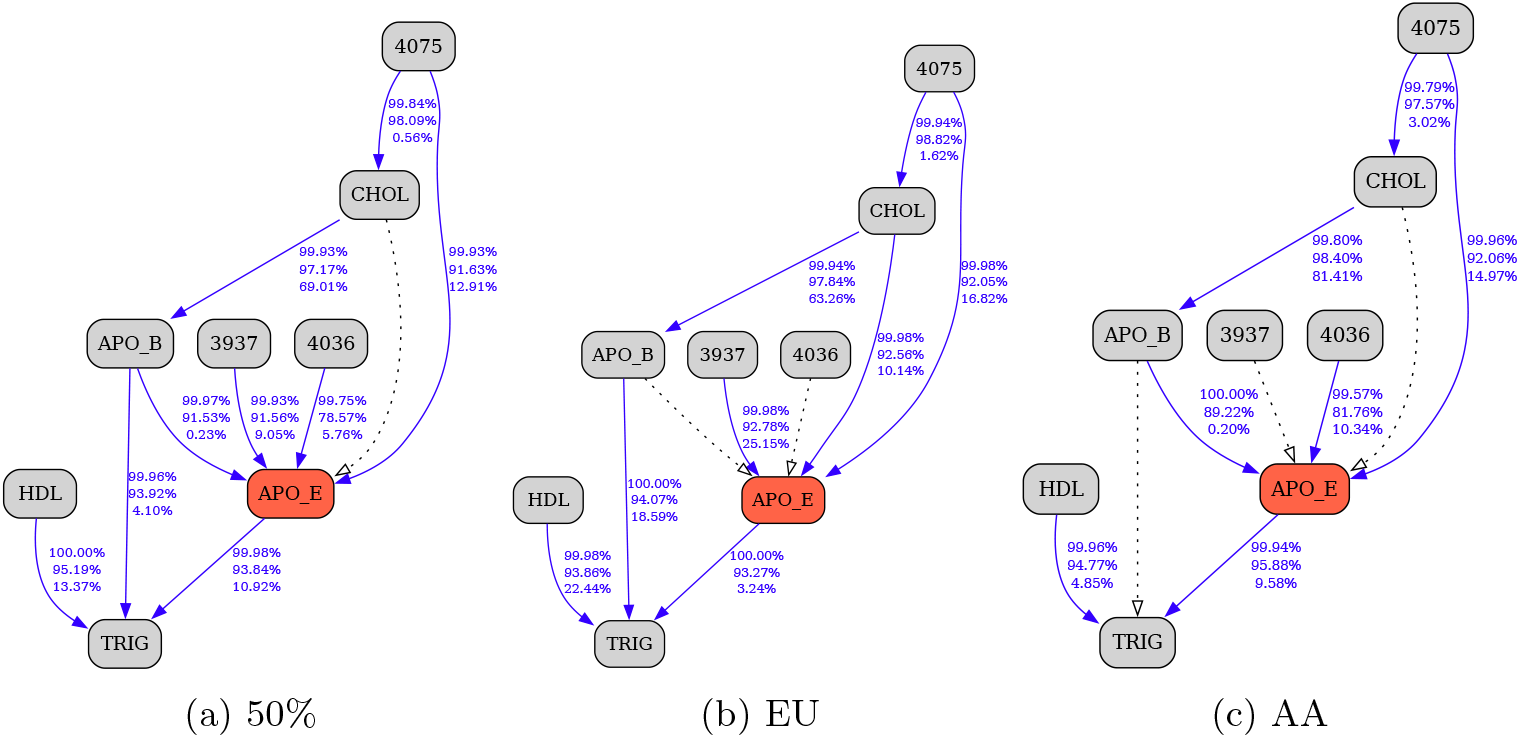
Learning via K2 criterion, 3-state discretization.

Figure 12 shows the MB obtained with K2 score from the 5-state discretized data (corresponding to Figure 10b). The 3937 – APOE dependency behaves as before, disappearing in the AA conditioned BN (Figure 12c). The 4036 – APOE edge behaves as expected, vanishing in Figure 12b. Other vanishing edges appear to be unstable due to subsampling. Note that here APOB – APOE edge is robust, while CHOL is pushed out of the MB envelope. Given that APOB is involved with cholesterol transport, it is likely “competing” with CHOL for the predictor role.

**Fig. 12.**
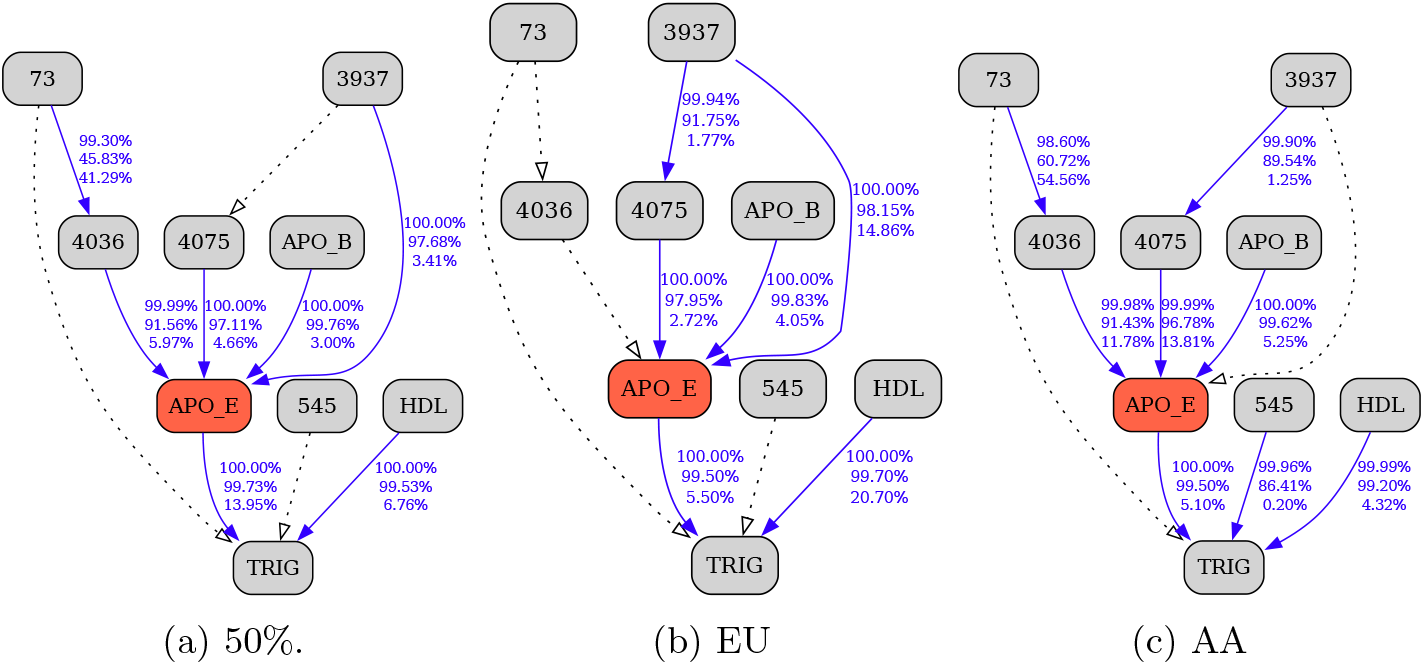
Learning via K2 criterion, 5-state discretization.

Figure 13 follows the procedure for the structure in Figure 10c, obtained with 7-state discretization, with similar results. Both the 3937 – APOE and the 4036 – APOE edges behave consistently with the previous observations. This time the model pulls CHOL into the MB, substituting the robust APOB – APOE dependency and validating the hypothesis that CHOL node is competing with APOB for the explanatory role. Here, the envelope is also missing HDL, another expected codependent of APOE that was present in the other two structures, illustrating the conflicted nature of K2 score.

**Fig. 13.**
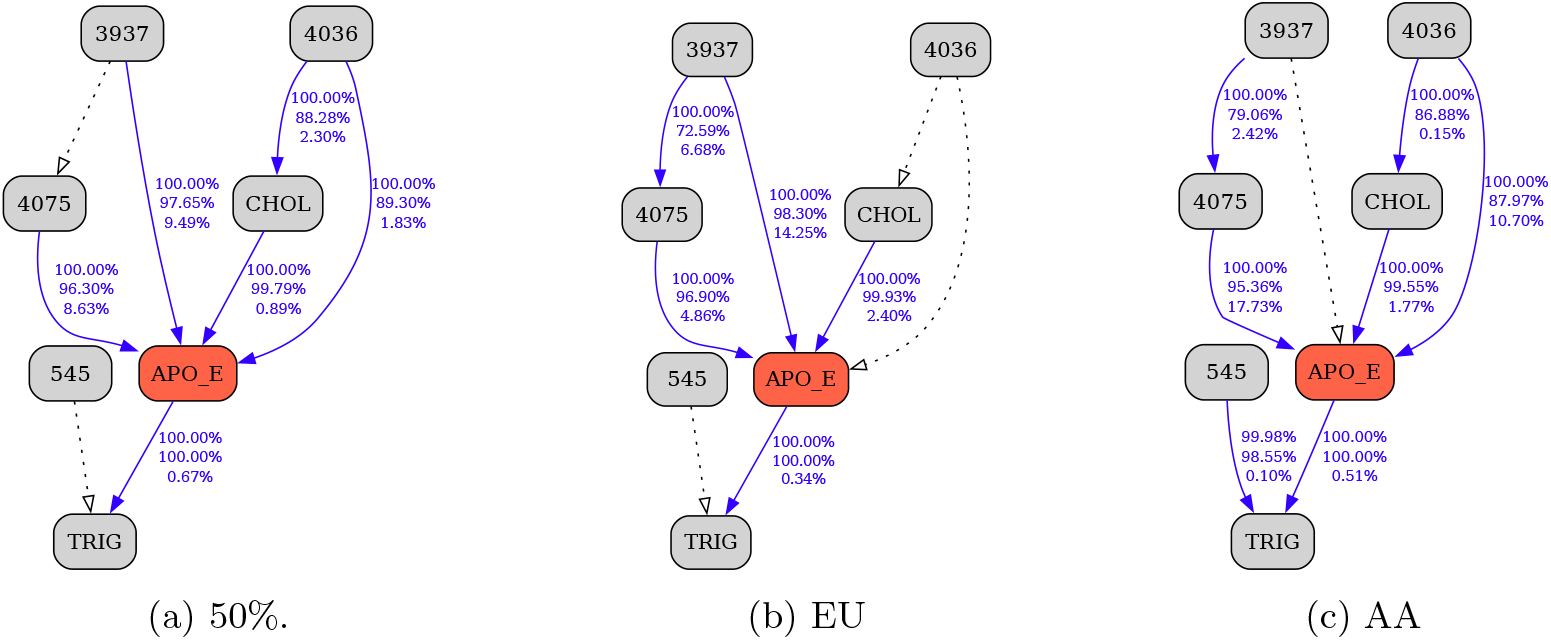
Learning via K2 criterion, 7-state discretization.

#### 3.2.3 MU Learning Characteristics

Figures 14-17 illustrate the results of the same analysis based on MU criterion (see Supplement B for full BNs). Unlike K2, MU yields MBs that share almost all nodes except APOB, which shows up only in the 3-state configuration (Figure 14a). Since 4075 is also present, this dependency can be used as a coherence indicator again. Note that MU correctly identifies all of the expected APOE dependencies (i.e., TRIG, CHOL, and HDL), including them in every envelope.

**Fig. 14.**
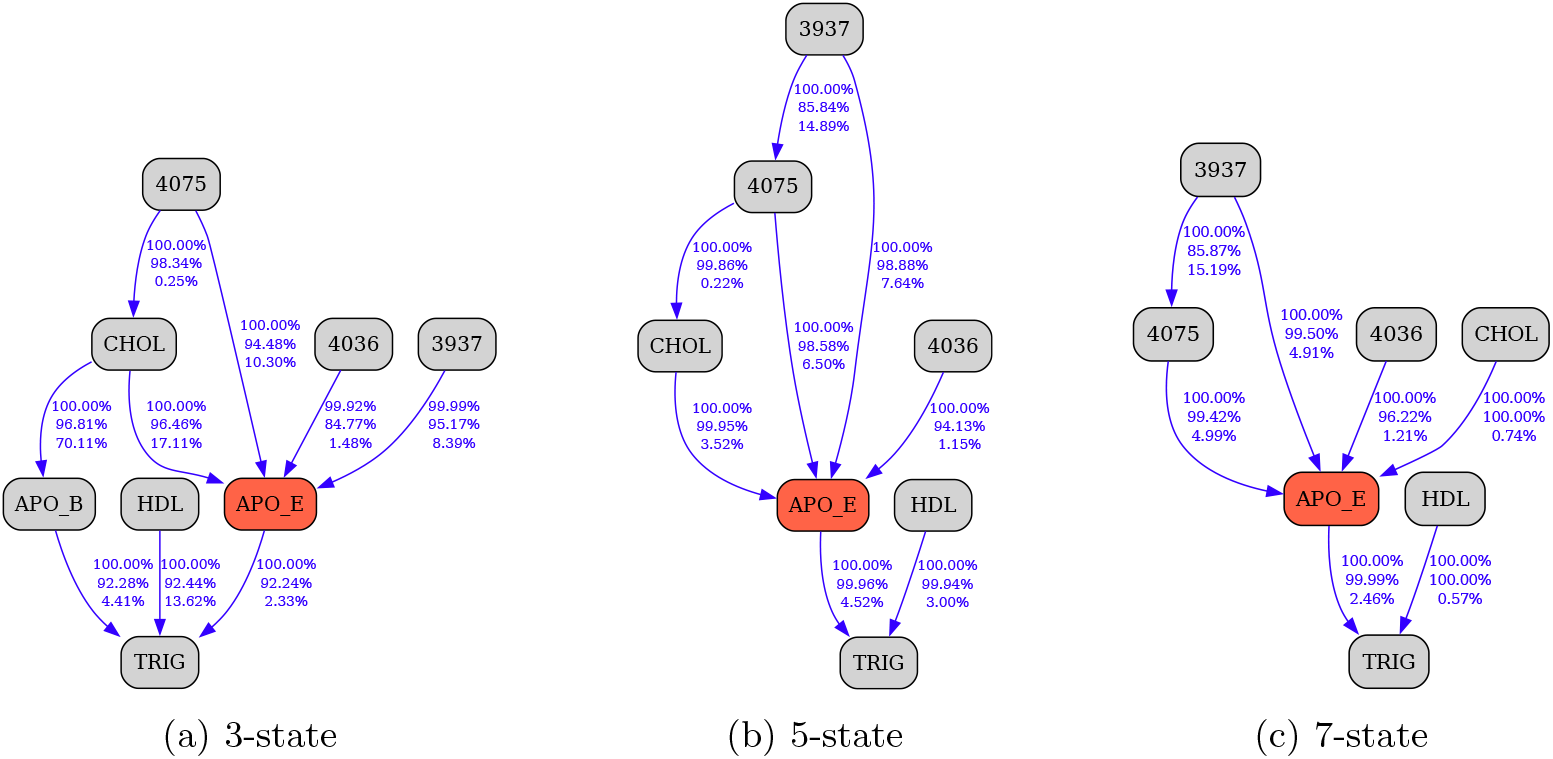
Learning APOE Markov blanket via MU criterion with data of varying discretization coarsenes.

Figure 15 (3-state) identifies the same pattern for 3937 – APOE edge and coherent behavior of 4075 – APOE, validating prior observations. There is edge instability around TRIG due to subsampling, which is likely a result of coarse discretization, given the inevitable information loss within the TRIG dependency complex composed of the exclusively continuous parents – APOE, APOB, and HDL.

**Fig. 15.**
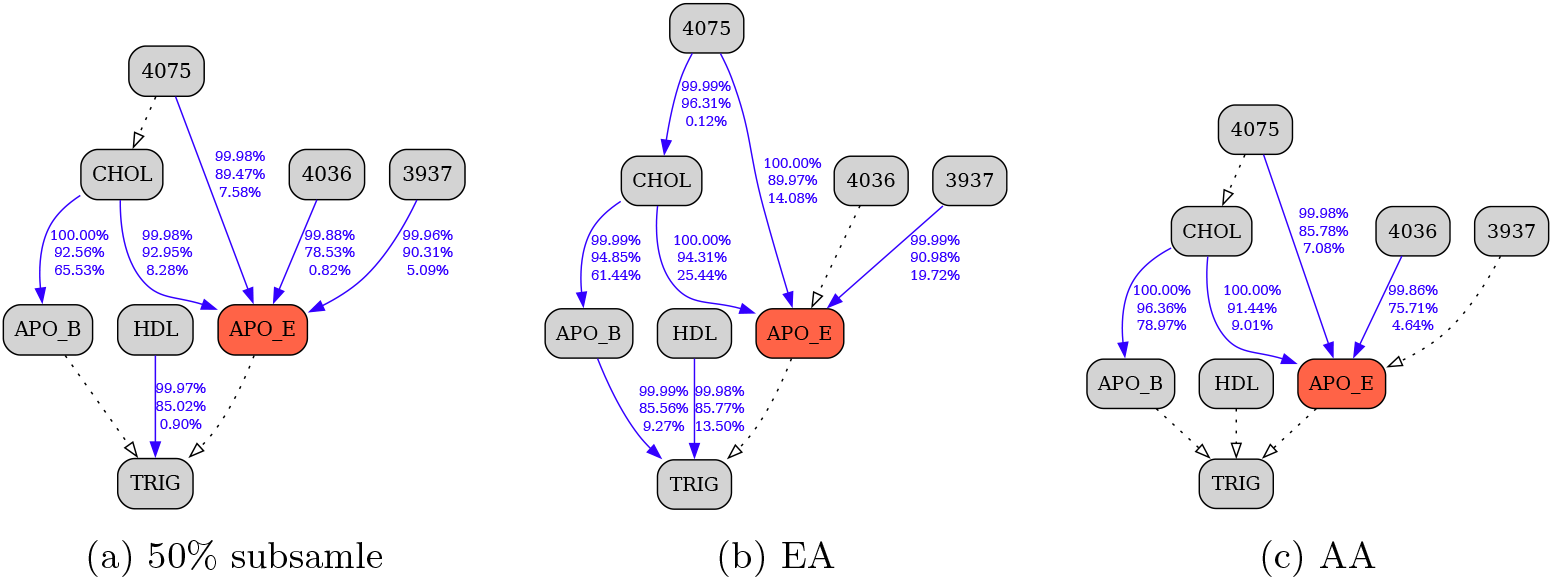
Learning via MU criterion, 3-state discretization.

Figure 16 (5-state) verifies the prior 3937 and 4036 findings. APOB is removed in favor of CHOL variable alone. The 4075 – CHOL edge is unstable under subsampling, as it was in the previous coarse-grained scenario.

**Fig. 16.**
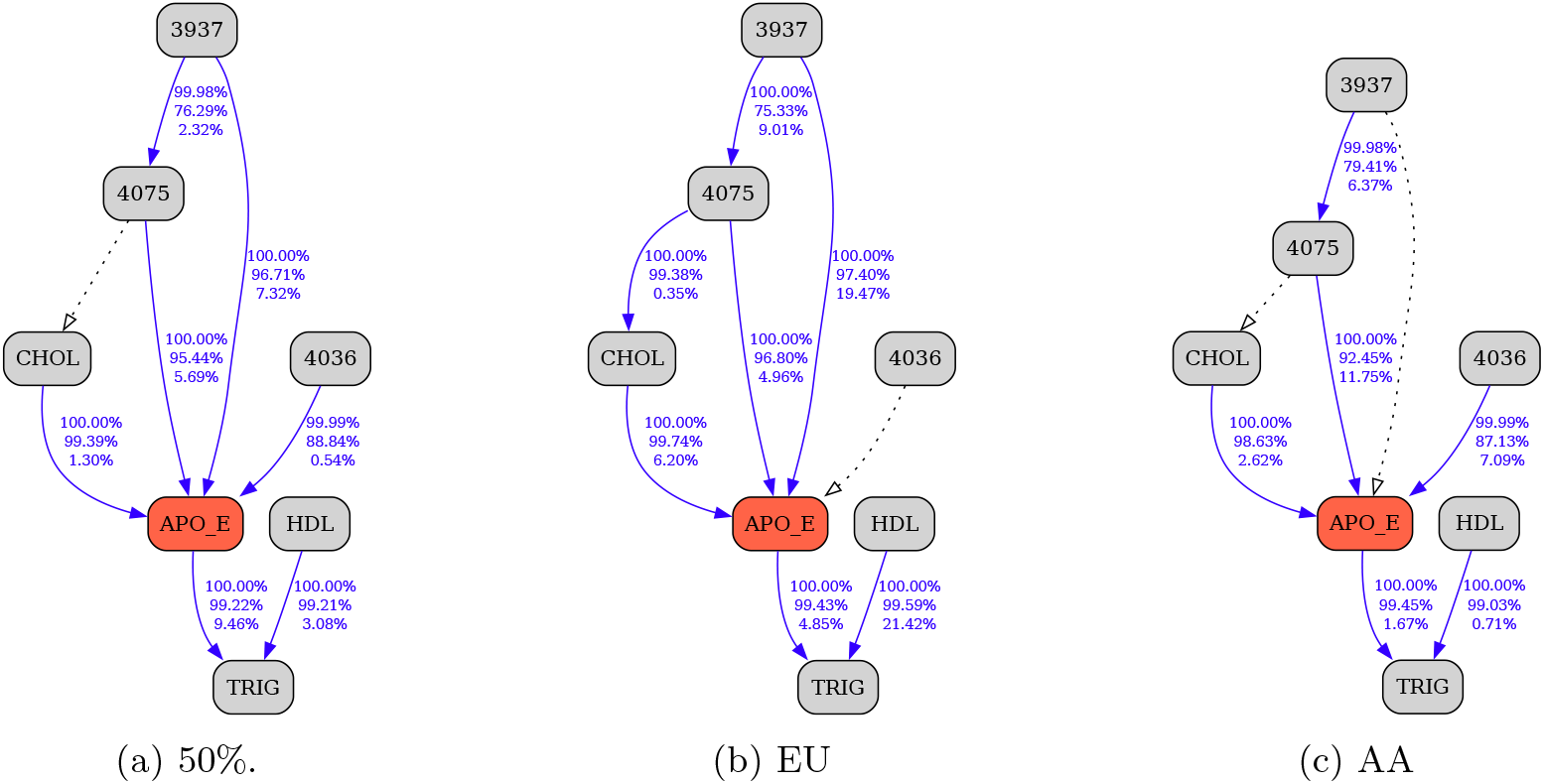
Learning via MU criterion, 5-state discretization.

Finally, Figure 17 repeats the analysis for the 7-state discretized data.

**Fig. 17.**
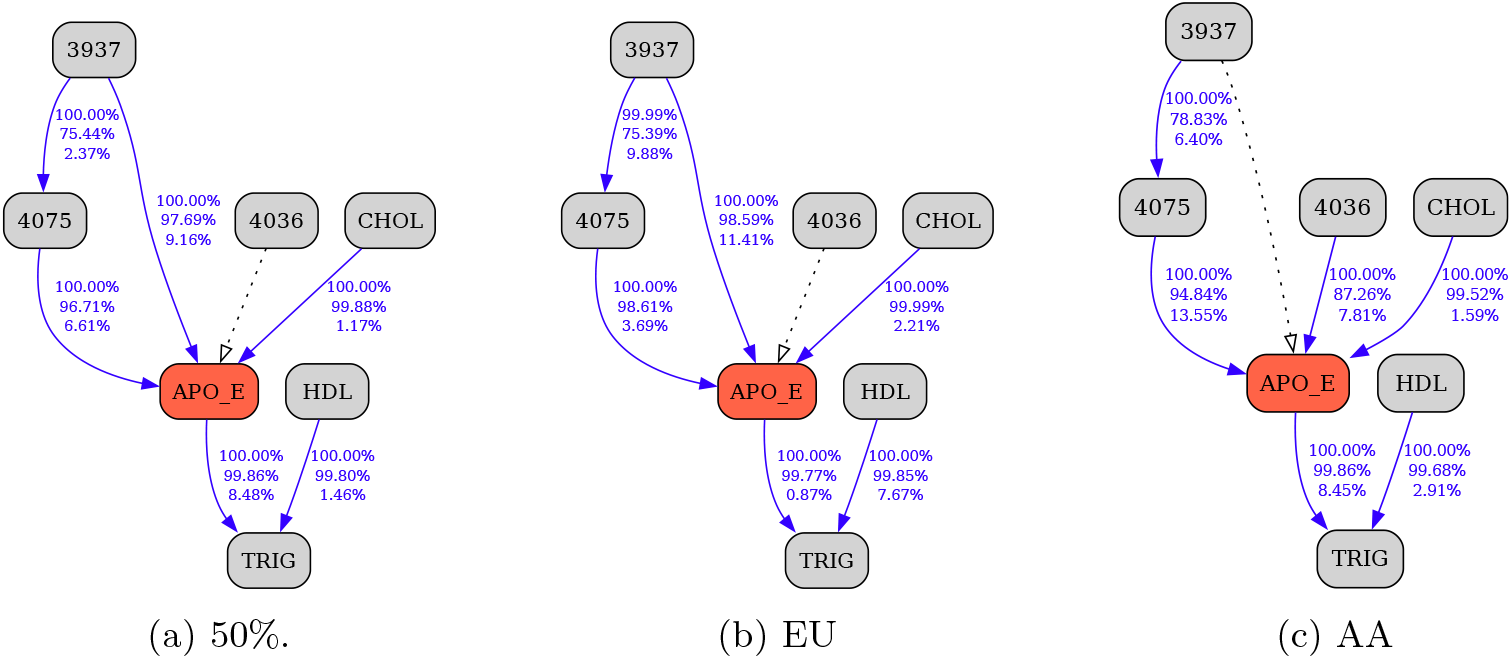
Learning via MU criterion, 7-state discretization.

Remarkably, unlike with the other two scoring functions, the Markov blanket of APOE remains almost entirely unchanged, retaining all the nodes and most of the structure found in the coarser-grained analysis. This observably more consistent behavior of the MU score replicates the observations compiled in Section 3.1, where its emphasis on specificity, combined with a competitive sensitivity profile, gives it the advantage of being more stable and authoritative. Subsampling and stratifying reveal the same pattern for the 3937 and the 4036 dependencies. As with K2, CHOL – 4075 edge ambiguity appearing in the coarser granularity experiments resolves with complete decoupling.

#### 3.2.4 Failure of MDL and Consistency of MU

Figure 18 shows MBs obtained with every score for the 9-state discretized data (see Supplement B for full BNs). Results repeating the stratified analyses are excluded because MDL lost the TRIG dependency, which served as its only sensitivity indicator. For all scores, 3937 exhibits the same pattern, appearing only in the EA strata, and 4036 behaves as expected where it is present. At this level of data granularity, K2 loses CHOL while MU remains almost unchanged, swapping CHOL for its codependent APOB (the substitution that was observed with K2 score before) and still correctly identifying the HDL association.

**Fig. 18.**
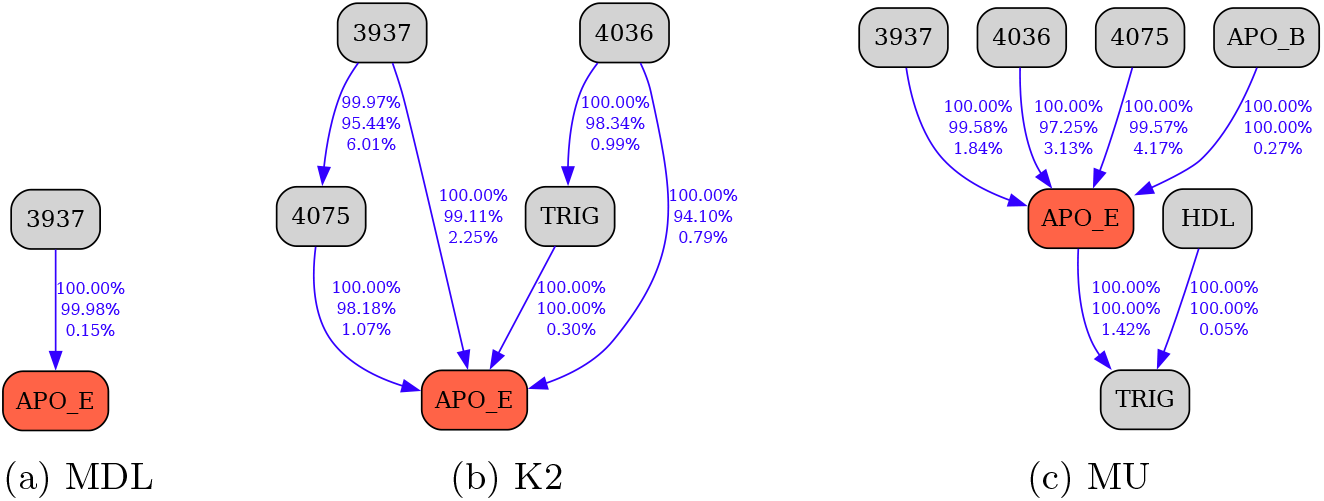
Learning APOE Markov blanket via MDL, K2, and MU for 9-state discretized data.

Overall, this “real world” application recapitulates the results obtained in 3.1 — not only does MU clearly outperform the standard information-theoretic MDL score, overcoming all of its limitations, but it also behaves more consistently than the best of the considered Bayesian scores (K2) by ramping up its specificity when the uncertainty is high due to data scarcity or the growing search space complexity.

## 4 Discussion

The new scoring criterion and the discriminative power evaluation framework developed here advance the MU model selection principle from a conceptual prototype into a fully functional, robust, calibrated instrument with quantified operating characteristics.

The original motivations for the uncertainty-based model selection principle included addressing scoring irregularities observed with MDL and the BD family, reconciling incommensurable scores across data sources, improving interpretability and generality of the learned network structures, and grounding model selection in observational evidence rather than abstract constructs (e.g., parsimony). The calibrated MU criterion satisfies all of the above and goes further, outperforming the other considered scoring functions across a variety of challenging configurations and exhibiting robust, consistent behavior over a wide range of parameters. The numerical simulation studies and multiscale genetic epidemiology application in Section 3 demonstrate performance and interpretability gains achieved through the principled operationalization of uncertainty-based model selection reasoning, with the computational complexity of the new MU criterion comparable to that of MDL, incurring negligible overhead to achieve its superior characteristics.

The discriminative power evaluation method presented in Section 2.7 has an independent merit: not only does it reveal performance gaps in other model selection principles (Section 3.1) but, more importantly, provides a context-independent measure of confidence (i.e., evidence quality) for any given dependency, which can be assessed and compared under varying parameters and sample sizes. This, in turn, enables a more nuanced, “customized” application and interpretation of BN models in practical research across diverse domains, with sensitivity, specificity, and normalized dependency strength assigned to each edge.

The computational burden of this quality assessment, while nontrivial, is modest relative to the search for an optimal model, as it is a post-processing step whose cost grows linearly with the number of reevaluated edges. The most demanding component is model sampling and reevaluation to accumulate sufficient statistics for the binary decision test. That said, algorithmic — or even analytical — shortcuts could substantially accelerate the procedure; even a statically-typed language reimplementation may yield one–to–two orders of magnitude speedups. Further algorithmic and analytical refinements may reduce the computational cost of the discriminative power evaluation procedure and broaden its use in future score design.

## 5 Conclusions

By grounding model selection in evidence quality, the new calibrated minimum uncertainty criterion delivers consistent gains over conventional scoring functions and enables reliable BN recovery across diverse application domains. The new discriminative power evaluation framework significantly aids the understanding of score-based BNSL, revealing score-performance irregularities that would otherwise be difficult to assess and mitigate. Independently, it enables context-free, interpretable, per-edge evaluation of recovered BNs. Overall, the methodology developed in this work facilitates straightforward integration of BN modeling, together with its precisely quantified interpretation, into research across disciplines seeking deeper and demonstrably credible insights.

## Declarations

### Ethics approval and consent to participate

Not applicable.

### Consent for publication

Not applicable.

### Availability of data and materials

The principal results of the study were obtained via numerical simulations. The APOE example data is described in Gogoshin et al. (2017) and references therein, and is available directly from the authors, or as part of the BNOmics package, at https://bitbucket.org/77D/bnomics.

Relevant code and software are available directly from the authors, or as part of the BNOmics package, at https://bitbucket.org/77D/bnomics.

### Competing interests

None declared.

### Funding

This work was supported by the NIH NLM R01LM013138 grant (to A.S.R.), NIH NLM R01LM013876 grant (to A.S.R.), Dr. Susumu Ohno Chair in Theoretical Biology (held by A.S.R.), and Susumu Ohno Distinguished Investigator Fellowship (to G.G.).

### Authors’ contributions

Grigoriy Gogoshin: Conceptualization; Data curation; Formal analysis; Investigation; Methodology; Software; Validation; Visualization; Writing - original draft. Andrei S. Rodin: Conceptualization; Funding acquisition; Investigation; Project administration; Supervision; Writing - original draft.

## Acknowledgements

The authors are grateful to Sergio Branciamore, Arthur D. Riggs, Nagarajan Vaidehi, Peter P. Lee, Amanda J. Myers, Russell C. Rockne, Nadia Carlesso, Konstancja Urbaniak, Babgen Manookian, Elizaveta Mukhaleva, and Grady Kestler for stimulating discussions on the accuracy, robustness, and interpretability of network modeling across diverse biomedical domains.

## Supplement

### Supplement A

**Table S1.** ACC (in bold) followed by TNR and TPR for a dependency model with Jeffrey’s prior over a variety of triples (card(*V*) : *V* ∈ (*Y, π, X*)) appearing in the first column.

| card | N | MDL |  |  | BDeu |  |  | K2 |  |  | MU |  |  |
| --- | --- | --- | --- | --- | --- | --- | --- | --- | --- | --- | --- | --- | --- |
| (2, 2, 2) | 100 | **0.80** | 0.98 | 0.61 | **0.73** | 0.59 | 0.87 | **0.81** | 0.89 | 0.73 | **0.80** | 0.97 | 0.62 |
|  | 200 | **0.85** | 0.99 | 0.70 | **0.77** | 0.64 | 0.91 | **0.86** | 0.93 | 0.80 | **0.85** | 0.99 | 0.71 |
|  | 300 | **0.87** | 0.99 | 0.75 | **0.80** | 0.68 | 0.92 | **0.89** | 0.94 | 0.84 | **0.87** | 0.99 | 0.75 |
|  | 400 | **0.89** | 1.00 | 0.78 | **0.82** | 0.70 | 0.94 | **0.91** | 0.95 | 0.86 | **0.89** | 0.99 | 0.78 |
|  | 500 | **0.90** | 1.00 | 0.81 | **0.83** | 0.72 | 0.94 | **0.92** | 0.96 | 0.88 | **0.90** | 0.99 | 0.80 |
|  | 600 | **0.91** | 1.00 | 0.83 | **0.84** | 0.74 | 0.95 | **0.93** | 0.96 | 0.89 | **0.91** | 1.00 | 0.82 |
| (3, 3, 3) | 100 | **0.76** | 1.00 | 0.52 | **0.92** | 0.87 | 0.96 | **0.93** | 0.93 | 0.93 | **0.94** | 0.98 | 0.91 |
|  | 200 | **0.87** | 1.00 | 0.75 | **0.96** | 0.93 | 0.99 | **0.97** | 0.97 | 0.97 | **0.98** | 0.99 | 0.96 |
|  | 300 | **0.92** | 1.00 | 0.85 | **0.97** | 0.95 | 0.99 | **0.99** | 0.99 | 0.99 | **0.99** | 1.00 | 0.98 |
|  | 400 | **0.95** | 1.00 | 0.90 | **0.98** | 0.96 | 1.00 | **0.99** | 0.99 | 0.99 | **0.99** | 1.00 | 0.99 |
|  | 500 | **0.96** | 1.00 | 0.93 | **0.98** | 0.97 | 1.00 | **0.99** | 0.99 | 0.99 | **0.99** | 1.00 | 0.99 |
|  | 600 | **0.97** | 1.00 | 0.94 | **0.99** | 0.98 | 1.00 | **1.00** | 1.00 | 1.00 | **1.00** | 1.00 | 0.99 |
| (4, 4, 4) | 100 | **0.53** | 1.00 | 0.07 | **0.97** | 0.99 | 0.95 | **0.95** | 0.93 | 0.97 | **0.97** | 0.97 | 0.97 |
|  | 200 | **0.65** | 1.00 | 0.30 | **0.99** | 1.00 | 0.99 | **0.99** | 0.98 | 1.00 | **1.00** | 1.00 | 1.00 |
|  | 300 | **0.79** | 1.00 | 0.58 | **1.00** | 1.00 | 1.00 | **1.00** | 1.00 | 1.00 | **1.00** | 1.00 | 1.00 |
|  | 400 | **0.87** | 1.00 | 0.75 | **1.00** | 1.00 | 1.00 | **1.00** | 1.00 | 1.00 | **1.00** | 1.00 | 1.00 |
|  | 500 | **0.92** | 1.00 | 0.85 | **1.00** | 1.00 | 1.00 | **1.00** | 1.00 | 1.00 | **1.00** | 1.00 | 1.00 |
|  | 600 | **0.95** | 1.00 | 0.90 | **1.00** | 1.00 | 1.00 | **1.00** | 1.00 | 1.00 | **1.00** | 1.00 | 1.00 |
| (5, 5, 5) | 100 | **0.50** | 1.00 | 0.00 | **0.93** | 1.00 | 0.86 | **0.95** | 0.91 | 0.98 | **0.97** | 0.96 | 0.99 |
|  | 200 | **0.50** | 1.00 | 0.01 | **0.99** | 1.00 | 0.97 | **0.99** | 0.98 | 1.00 | **1.00** | 0.99 | 1.00 |
|  | 300 | **0.52** | 1.00 | 0.05 | **1.00** | 1.00 | 0.99 | **1.00** | 1.00 | 1.00 | **1.00** | 1.00 | 1.00 |
|  | 400 | **0.58** | 1.00 | 0.15 | **1.00** | 1.00 | 1.00 | **1.00** | 1.00 | 1.00 | **1.00** | 1.00 | 1.00 |
|  | 500 | **0.66** | 1.00 | 0.33 | **1.00** | 1.00 | 1.00 | **1.00** | 1.00 | 1.00 | **1.00** | 1.00 | 1.00 |
|  | 600 | **0.76** | 1.00 | 0.52 | **1.00** | 1.00 | 1.00 | **1.00** | 1.00 | 1.00 | **1.00** | 1.00 | 1.00 |
| (6, 6, 6) | 100 | **0.50** | 1.00 | 0.00 | **0.83** | 1.00 | 0.65 | **0.93** | 0.87 | 0.99 | **0.97** | 0.97 | 0.98 |
|  | 200 | **0.50** | 1.00 | 0.00 | **0.94** | 1.00 | 0.87 | **0.99** | 0.98 | 1.00 | **1.00** | 0.99 | 1.00 |
|  | 300 | **0.50** | 1.00 | 0.00 | **0.98** | 1.00 | 0.96 | **1.00** | 1.00 | 1.00 | **1.00** | 1.00 | 1.00 |
|  | 400 | **0.50** | 1.00 | 0.00 | **0.99** | 1.00 | 0.99 | **1.00** | 1.00 | 1.00 | **1.00** | 1.00 | 1.00 |
|  | 500 | **0.50** | 1.00 | 0.01 | **1.00** | 1.00 | 1.00 | **1.00** | 1.00 | 1.00 | **1.00** | 1.00 | 1.00 |
|  | 600 | **0.51** | 1.00 | 0.02 | **1.00** | 1.00 | 1.00 | **1.00** | 1.00 | 1.00 | **1.00** | 1.00 | 1.00 |
| (7, 7, 7) | 100 | **0.50** | 1.00 | 0.00 | **0.70** | 1.00 | 0.41 | **0.91** | 0.82 | 0.99 | **0.96** | 0.99 | 0.94 |
|  | 200 | **0.50** | 1.00 | 0.00 | **0.80** | 1.00 | 0.61 | **0.98** | 0.96 | 1.00 | **1.00** | 0.99 | 1.00 |
|  | 300 | **0.50** | 1.00 | 0.00 | **0.89** | 1.00 | 0.79 | **1.00** | 0.99 | 1.00 | **1.00** | 1.00 | 1.00 |
|  | 400 | **0.50** | 1.00 | 0.00 | **0.95** | 1.00 | 0.90 | **1.00** | 1.00 | 1.00 | **1.00** | 1.00 | 1.00 |
|  | 500 | **0.50** | 1.00 | 0.00 | **0.98** | 1.00 | 0.96 | **1.00** | 1.00 | 1.00 | **1.00** | 1.00 | 1.00 |
|  | 600 | **0.50** | 1.00 | 0.00 | **0.99** | 1.00 | 0.98 | **1.00** | 1.00 | 1.00 | **1.00** | 1.00 | 1.00 |
| (8, 8, 8) | 100 | **0.50** | 1.00 | 0.00 | **0.61** | 1.00 | 0.23 | **0.87** | 0.76 | 0.99 | **0.85** | 1.00 | 0.71 |
|  | 200 | **0.50** | 1.00 | 0.00 | **0.65** | 1.00 | 0.29 | **0.96** | 0.93 | 1.00 | **1.00** | 1.00 | 1.00 |
|  | 300 | **0.50** | 1.00 | 0.00 | **0.71** | 1.00 | 0.42 | **0.99** | 0.98 | 1.00 | **1.00** | 1.00 | 1.00 |
|  | 400 | **0.50** | 1.00 | 0.00 | **0.79** | 1.00 | 0.58 | **1.00** | 1.00 | 1.00 | **1.00** | 1.00 | 1.00 |
|  | 500 | **0.50** | 1.00 | 0.00 | **0.86** | 1.00 | 0.72 | **1.00** | 1.00 | 1.00 | **1.00** | 1.00 | 1.00 |
|  | 600 | **0.50** | 1.00 | 0.00 | **0.92** | 1.00 | 0.83 | **1.00** | 1.00 | 1.00 | **1.00** | 1.00 | 1.00 |
| (9, 9, 9) | 100 | **0.50** | 1.00 | 0.00 | **0.56** | 1.00 | 0.13 | **0.84** | 0.70 | 0.99 | **0.64** | 1.00 | 0.28 |
|  | 200 | **0.50** | 1.00 | 0.00 | **0.55** | 1.00 | 0.10 | **0.94** | 0.88 | 1.00 | **0.98** | 1.00 | 0.96 |
|  | 300 | **0.50** | 1.00 | 0.00 | **0.57** | 1.00 | 0.14 | **0.98** | 0.96 | 1.00 | **1.00** | 1.00 | 1.00 |
|  | 400 | **0.50** | 1.00 | 0.00 | **0.60** | 1.00 | 0.20 | **1.00** | 0.99 | 1.00 | **1.00** | 1.00 | 1.00 |
|  | 500 | **0.50** | 1.00 | 0.00 | **0.65** | 1.00 | 0.30 | **1.00** | 1.00 | 1.00 | **1.00** | 1.00 | 1.00 |
|  | 600 | **0.50** | 1.00 | 0.00 | **0.70** | 1.00 | 0.41 | **1.00** | 1.00 | 1.00 | **1.00** | 1.00 | 1.00 |

**Table S2.** ACC (in bold) followed by TNR and TPR for a dependency model with Jeffrey’s prior over a variety of cardinality triples (card(*V*) : *V* ∈ (*Y, π, X*)) appearing in the first column.

| card | N | MDL |  |  | BDeu |  |  | K2 |  |  | MU |  |  |
| --- | --- | --- | --- | --- | --- | --- | --- | --- | --- | --- | --- | --- | --- |
| $(2, 3 \times 2 \times 3, 2)$ | 100 | **0.57** | 1.00 | 0.13 | **0.84** | 0.93 | 0.76 | **0.83** | 0.84 | 0.81 | **0.83** | 0.88 | 0.79 |
|  | 200 | **0.63** | 1.00 | 0.26 | **0.92** | 0.97 | 0.86 | **0.91** | 0.92 | 0.90 | **0.90** | 0.97 | 0.83 |
|  | 300 | **0.70** | 1.00 | 0.39 | **0.95** | 0.99 | 0.91 | **0.95** | 0.95 | 0.94 | **0.93** | 0.99 | 0.86 |
|  | 400 | **0.75** | 1.00 | 0.51 | **0.96** | 0.99 | 0.94 | **0.96** | 0.97 | 0.96 | **0.94** | 0.99 | 0.89 |
|  | 500 | **0.80** | 1.00 | 0.60 | **0.97** | 0.99 | 0.95 | **0.97** | 0.98 | 0.97 | **0.95** | 0.99 | 0.91 |
|  | 600 | **0.83** | 1.00 | 0.66 | **0.98** | 1.00 | 0.96 | **0.98** | 0.98 | 0.98 | **0.96** | 1.00 | 0.93 |
| $(3, 3 \times 2 \times 3, 3)$ | 100 | **0.50** | 1.00 | 0.00 | **0.85** | 1.00 | 0.70 | **0.89** | 0.87 | 0.92 | **0.89** | 0.83 | 0.95 |
|  | 200 | **0.50** | 1.00 | 0.00 | **0.92** | 1.00 | 0.84 | **0.97** | 0.96 | 0.98 | **0.98** | 0.98 | 0.98 |
|  | 300 | **0.51** | 1.00 | 0.01 | **0.96** | 1.00 | 0.92 | **0.99** | 0.99 | 0.99 | **0.99** | 1.00 | 0.99 |
|  | 400 | **0.52** | 1.00 | 0.04 | **0.98** | 1.00 | 0.96 | **1.00** | 0.99 | 1.00 | **1.00** | 1.00 | 1.00 |
|  | 500 | **0.54** | 1.00 | 0.09 | **0.99** | 1.00 | 0.97 | **1.00** | 1.00 | 1.00 | **1.00** | 1.00 | 1.00 |
|  | 600 | **0.59** | 1.00 | 0.17 | **0.99** | 1.00 | 0.98 | **1.00** | 1.00 | 1.00 | **1.00** | 1.00 | 1.00 |
| $(4, 3 \times 2 \times 3, 4)$ | 100 | **0.50** | 1.00 | 0.00 | **0.77** | 1.00 | 0.55 | **0.90** | 0.85 | 0.95 | **0.90** | 0.84 | 0.97 |
|  | 200 | **0.50** | 1.00 | 0.00 | **0.85** | 1.00 | 0.70 | **0.98** | 0.96 | 0.99 | **0.99** | 0.98 | 1.00 |
|  | 300 | **0.50** | 1.00 | 0.00 | **0.91** | 1.00 | 0.82 | **0.99** | 0.99 | 1.00 | **1.00** | 1.00 | 1.00 |
|  | 400 | **0.50** | 1.00 | 0.00 | **0.95** | 1.00 | 0.90 | **1.00** | 1.00 | 1.00 | **1.00** | 1.00 | 1.00 |
|  | 500 | **0.50** | 1.00 | 0.00 | **0.97** | 1.00 | 0.94 | **1.00** | 1.00 | 1.00 | **1.00** | 1.00 | 1.00 |
|  | 600 | **0.50** | 1.00 | 0.00 | **0.98** | 1.00 | 0.97 | **1.00** | 1.00 | 1.00 | **1.00** | 1.00 | 1.00 |
| $(5, 3 \times 2 \times 3, 5)$ | 100 | **0.50** | 1.00 | 0.00 | **0.70** | 1.00 | 0.40 | **0.88** | 0.81 | 0.96 | **0.93** | 0.91 | 0.94 |
|  | 200 | **0.50** | 1.00 | 0.00 | **0.73** | 1.00 | 0.47 | **0.97** | 0.94 | 1.00 | **0.99** | 0.99 | 1.00 |
|  | 300 | **0.50** | 1.00 | 0.00 | **0.79** | 1.00 | 0.59 | **0.99** | 0.99 | 1.00 | **1.00** | 1.00 | 1.00 |
|  | 400 | **0.50** | 1.00 | 0.00 | **0.85** | 1.00 | 0.71 | **1.00** | 1.00 | 1.00 | **1.00** | 1.00 | 1.00 |
|  | 500 | **0.50** | 1.00 | 0.00 | **0.90** | 1.00 | 0.80 | **1.00** | 1.00 | 1.00 | **1.00** | 1.00 | 1.00 |
|  | 600 | **0.50** | 1.00 | 0.00 | **0.94** | 1.00 | 0.88 | **1.00** | 1.00 | 1.00 | **1.00** | 1.00 | 1.00 |
| $(6, 3 \times 2 \times 3, 6)$ | 100 | **0.50** | 1.00 | 0.00 | **0.64** | 1.00 | 0.28 | **0.86** | 0.76 | 0.96 | **0.89** | 0.98 | 0.79 |
|  | 200 | **0.50** | 1.00 | 0.00 | **0.63** | 1.00 | 0.26 | **0.96** | 0.91 | 1.00 | **0.99** | 1.00 | 0.99 |
|  | 300 | **0.50** | 1.00 | 0.00 | **0.65** | 1.00 | 0.30 | **0.99** | 0.98 | 1.00 | **1.00** | 1.00 | 1.00 |
|  | 400 | **0.50** | 1.00 | 0.00 | **0.69** | 1.00 | 0.39 | **1.00** | 0.99 | 1.00 | **1.00** | 1.00 | 1.00 |
|  | 500 | **0.50** | 1.00 | 0.00 | **0.74** | 1.00 | 0.49 | **1.00** | 1.00 | 1.00 | **1.00** | 1.00 | 1.00 |
|  | 600 | **0.50** | 1.00 | 0.00 | **0.80** | 1.00 | 0.59 | **1.00** | 1.00 | 1.00 | **1.00** | 1.00 | 1.00 |
| $(7, 3 \times 2 \times 3, 7)$ | 100 | **0.50** | 1.00 | 0.00 | **0.61** | 1.00 | 0.22 | **0.84** | 0.71 | 0.96 | **0.69** | 1.00 | 0.39 |
|  | 200 | **0.50** | 1.00 | 0.00 | **0.57** | 1.00 | 0.13 | **0.94** | 0.88 | 1.00 | **0.97** | 1.00 | 0.94 |
|  | 300 | **0.50** | 1.00 | 0.00 | **0.56** | 1.00 | 0.12 | **0.98** | 0.95 | 1.00 | **1.00** | 1.00 | 1.00 |
|  | 400 | **0.50** | 1.00 | 0.00 | **0.57** | 1.00 | 0.14 | **0.99** | 0.99 | 1.00 | **1.00** | 1.00 | 1.00 |
|  | 500 | **0.50** | 1.00 | 0.00 | **0.59** | 1.00 | 0.18 | **1.00** | 1.00 | 1.00 | **1.00** | 1.00 | 1.00 |
|  | 600 | **0.50** | 1.00 | 0.00 | **0.62** | 1.00 | 0.24 | **1.00** | 1.00 | 1.00 | **1.00** | 1.00 | 1.00 |
| $(8, 3 \times 2 \times 3, 8)$ | 100 | **0.50** | 1.00 | 0.00 | **0.59** | 1.00 | 0.18 | **0.81** | 0.66 | 0.97 | **0.54** | 1.00 | 0.07 |
|  | 200 | **0.50** | 1.00 | 0.00 | **0.53** | 1.00 | 0.07 | **0.91** | 0.83 | 1.00 | **0.84** | 1.00 | 0.68 |
|  | 300 | **0.50** | 1.00 | 0.00 | **0.52** | 1.00 | 0.04 | **0.96** | 0.92 | 1.00 | **0.99** | 1.00 | 0.98 |
|  | 400 | **0.50** | 1.00 | 0.00 | **0.52** | 1.00 | 0.04 | **0.99** | 0.97 | 1.00 | **1.00** | 1.00 | 1.00 |
|  | 500 | **0.50** | 1.00 | 0.00 | **0.52** | 1.00 | 0.04 | **0.99** | 0.99 | 1.00 | **1.00** | 1.00 | 1.00 |
|  | 600 | **0.50** | 1.00 | 0.00 | **0.53** | 1.00 | 0.05 | **1.00** | 1.00 | 1.00 | **1.00** | 1.00 | 1.00 |
| $(9, 3 \times 2 \times 3, 9)$ | 100 | **0.50** | 1.00 | 0.00 | **0.58** | 1.00 | 0.16 | **0.79** | 0.62 | 0.96 | **0.50** | 1.00 | 0.01 |
|  | 200 | **0.50** | 1.00 | 0.00 | **0.52** | 1.00 | 0.04 | **0.88** | 0.77 | 1.00 | **0.63** | 1.00 | 0.25 |
|  | 300 | **0.50** | 1.00 | 0.00 | **0.51** | 1.00 | 0.01 | **0.94** | 0.88 | 1.00 | **0.92** | 1.00 | 0.84 |
|  | 400 | **0.50** | 1.00 | 0.00 | **0.50** | 1.00 | 0.01 | **0.97** | 0.95 | 1.00 | **1.00** | 1.00 | 0.99 |
|  | 500 | **0.50** | 1.00 | 0.00 | **0.50** | 1.00 | 0.01 | **0.99** | 0.98 | 1.00 | **1.00** | 1.00 | 1.00 |
|  | 600 | **0.50** | 1.00 | 0.00 | **0.50** | 1.00 | 0.01 | **1.00** | 0.99 | 1.00 | **1.00** | 1.00 | 1.00 |

**Table S3.** ACC (in bold) followed by TNR and TPR for a dependency model with a flat prior with a variety of cardinality triples (card(*V*) : *V* ∈ (*Y, π, X*)) appearing in the first column

| card | N | MDL |  |  | BDeu |  |  | K2 |  |  | MU |  |  |
| --- | --- | --- | --- | --- | --- | --- | --- | --- | --- | --- | --- | --- | --- |
| (2, 2, 2) | 100 | **0.74** | 0.98 | 0.49 | **0.74** | 0.69 | 0.78 | **0.78** | 0.87 | 0.69 | **0.74** | 0.97 | 0.51 |
|  | 200 | **0.80** | 0.99 | 0.60 | **0.79** | 0.74 | 0.84 | **0.84** | 0.91 | 0.76 | **0.79** | 0.98 | 0.60 |
|  | 300 | **0.83** | 0.99 | 0.66 | **0.82** | 0.78 | 0.86 | **0.87** | 0.93 | 0.80 | **0.82** | 0.99 | 0.65 |
|  | 400 | **0.85** | 1.00 | 0.70 | **0.84** | 0.80 | 0.88 | **0.88** | 0.94 | 0.83 | **0.84** | 0.99 | 0.69 |
|  | 500 | **0.86** | 1.00 | 0.73 | **0.85** | 0.82 | 0.89 | **0.90** | 0.95 | 0.85 | **0.86** | 0.99 | 0.72 |
|  | 600 | **0.88** | 1.00 | 0.75 | **0.86** | 0.83 | 0.90 | **0.91** | 0.95 | 0.86 | **0.87** | 0.99 | 0.74 |
| (3, 3, 3) | 100 | **0.63** | 1.00 | 0.26 | **0.89** | 0.93 | 0.85 | **0.89** | 0.88 | 0.89 | **0.87** | 0.97 | 0.78 |
|  | 200 | **0.75** | 1.00 | 0.49 | **0.95** | 0.97 | 0.93 | **0.95** | 0.95 | 0.95 | **0.93** | 0.99 | 0.88 |
|  | 300 | **0.82** | 1.00 | 0.64 | **0.97** | 0.98 | 0.96 | **0.97** | 0.98 | 0.97 | **0.96** | 1.00 | 0.92 |
|  | 400 | **0.87** | 1.00 | 0.73 | **0.98** | 0.99 | 0.97 | **0.98** | 0.98 | 0.98 | **0.97** | 1.00 | 0.95 |
|  | 500 | **0.90** | 1.00 | 0.80 | **0.99** | 0.99 | 0.98 | **0.99** | 0.99 | 0.99 | **0.98** | 1.00 | 0.96 |
|  | 600 | **0.92** | 1.00 | 0.84 | **0.99** | 0.99 | 0.99 | **0.99** | 0.99 | 0.99 | **0.99** | 1.00 | 0.97 |
| (4, 4, 4) | 100 | **0.51** | 1.00 | 0.01 | **0.85** | 1.00 | 0.70 | **0.90** | 0.86 | 0.95 | **0.92** | 0.95 | 0.89 |
|  | 200 | **0.53** | 1.00 | 0.06 | **0.93** | 1.00 | 0.87 | **0.97** | 0.96 | 0.99 | **0.98** | 0.99 | 0.96 |
|  | 300 | **0.58** | 1.00 | 0.16 | **0.97** | 1.00 | 0.94 | **0.99** | 0.99 | 0.99 | **0.99** | 1.00 | 0.99 |
|  | 400 | **0.66** | 1.00 | 0.31 | **0.98** | 1.00 | 0.97 | **1.00** | 0.99 | 1.00 | **1.00** | 1.00 | 0.99 |
|  | 500 | **0.73** | 1.00 | 0.46 | **0.99** | 1.00 | 0.98 | **1.00** | 1.00 | 1.00 | **1.00** | 1.00 | 1.00 |
|  | 600 | **0.79** | 1.00 | 0.59 | **1.00** | 1.00 | 0.99 | **1.00** | 1.00 | 1.00 | **1.00** | 1.00 | 1.00 |
| (5, 5, 5) | 100 | **0.50** | 1.00 | 0.00 | **0.69** | 1.00 | 0.39 | **0.88** | 0.80 | 0.96 | **0.93** | 0.93 | 0.93 |
|  | 200 | **0.50** | 1.00 | 0.00 | **0.79** | 1.00 | 0.58 | **0.97** | 0.95 | 0.99 | **0.99** | 0.99 | 0.99 |
|  | 300 | **0.50** | 1.00 | 0.00 | **0.88** | 1.00 | 0.75 | **0.99** | 0.99 | 1.00 | **1.00** | 1.00 | 1.00 |
|  | 400 | **0.50** | 1.00 | 0.01 | **0.93** | 1.00 | 0.86 | **1.00** | 1.00 | 1.00 | **1.00** | 1.00 | 1.00 |
|  | 500 | **0.51** | 1.00 | 0.02 | **0.96** | 1.00 | 0.92 | **1.00** | 1.00 | 1.00 | **1.00** | 1.00 | 1.00 |
|  | 600 | **0.53** | 1.00 | 0.05 | **0.98** | 1.00 | 0.96 | **1.00** | 1.00 | 1.00 | **1.00** | 1.00 | 1.00 |
| (6, 6, 6) | 100 | **0.50** | 1.00 | 0.00 | **0.57** | 1.00 | 0.14 | **0.84** | 0.70 | 0.97 | **0.92** | 0.93 | 0.90 |
|  | 200 | **0.50** | 1.00 | 0.00 | **0.60** | 1.00 | 0.19 | **0.95** | 0.91 | 1.00 | **0.99** | 0.98 | 0.99 |
|  | 300 | **0.50** | 1.00 | 0.00 | **0.65** | 1.00 | 0.30 | **0.99** | 0.98 | 1.00 | **1.00** | 1.00 | 1.00 |
|  | 400 | **0.50** | 1.00 | 0.00 | **0.72** | 1.00 | 0.44 | **1.00** | 0.99 | 1.00 | **1.00** | 1.00 | 1.00 |
|  | 500 | **0.50** | 1.00 | 0.00 | **0.79** | 1.00 | 0.57 | **1.00** | 1.00 | 1.00 | **1.00** | 1.00 | 1.00 |
|  | 600 | **0.50** | 1.00 | 0.00 | **0.84** | 1.00 | 0.69 | **1.00** | 1.00 | 1.00 | **1.00** | 1.00 | 1.00 |
| (7, 7, 7) | 100 | **0.50** | 1.00 | 0.00 | **0.52** | 1.00 | 0.04 | **0.79** | 0.60 | 0.98 | **0.87** | 0.96 | 0.78 |
|  | 200 | **0.50** | 1.00 | 0.00 | **0.52** | 1.00 | 0.03 | **0.92** | 0.84 | 1.00 | **0.98** | 0.98 | 0.98 |
|  | 300 | **0.50** | 1.00 | 0.00 | **0.52** | 1.00 | 0.04 | **0.98** | 0.95 | 1.00 | **1.00** | 0.99 | 1.00 |
|  | 400 | **0.50** | 1.00 | 0.00 | **0.53** | 1.00 | 0.07 | **0.99** | 0.99 | 1.00 | **1.00** | 1.00 | 1.00 |
|  | 500 | **0.50** | 1.00 | 0.00 | **0.56** | 1.00 | 0.11 | **1.00** | 1.00 | 1.00 | **1.00** | 1.00 | 1.00 |
|  | 600 | **0.50** | 1.00 | 0.00 | **0.58** | 1.00 | 0.17 | **1.00** | 1.00 | 1.00 | **1.00** | 1.00 | 1.00 |
| (8, 8, 8) | 100 | **0.50** | 1.00 | 0.00 | **0.51** | 1.00 | 0.01 | **0.74** | 0.51 | 0.97 | **0.71** | 0.99 | 0.44 |
|  | 200 | **0.50** | 1.00 | 0.00 | **0.50** | 1.00 | 0.00 | **0.87** | 0.73 | 1.00 | **0.97** | 0.99 | 0.94 |
|  | 300 | **0.50** | 1.00 | 0.00 | **0.50** | 1.00 | 0.00 | **0.95** | 0.90 | 1.00 | **1.00** | 1.00 | 1.00 |
|  | 400 | **0.50** | 1.00 | 0.00 | **0.50** | 1.00 | 0.00 | **0.98** | 0.97 | 1.00 | **1.00** | 1.00 | 1.00 |
|  | 500 | **0.50** | 1.00 | 0.00 | **0.50** | 1.00 | 0.01 | **1.00** | 0.99 | 1.00 | **1.00** | 1.00 | 1.00 |
|  | 600 | **0.50** | 1.00 | 0.00 | **0.51** | 1.00 | 0.01 | **1.00** | 1.00 | 1.00 | **1.00** | 1.00 | 1.00 |
| (9, 9, 9) | 100 | **0.50** | 1.00 | 0.00 | **0.50** | 1.00 | 0.01 | **0.70** | 0.43 | 0.97 | **0.55** | 1.00 | 0.11 |
|  | 200 | **0.50** | 1.00 | 0.00 | **0.50** | 1.00 | 0.00 | **0.81** | 0.62 | 1.00 | **0.87** | 1.00 | 0.74 |
|  | 300 | **0.50** | 1.00 | 0.00 | **0.50** | 1.00 | 0.00 | **0.90** | 0.80 | 1.00 | **0.99** | 1.00 | 0.98 |
|  | 400 | **0.50** | 1.00 | 0.00 | **0.50** | 1.00 | 0.00 | **0.96** | 0.92 | 1.00 | **1.00** | 1.00 | 1.00 |
|  | 500 | **0.50** | 1.00 | 0.00 | **0.50** | 1.00 | 0.00 | **0.99** | 0.97 | 1.00 | **1.00** | 1.00 | 1.00 |
|  | 600 | **0.50** | 1.00 | 0.00 | **0.50** | 1.00 | 0.00 | **1.00** | 0.99 | 1.00 | **1.00** | 1.00 | 1.00 |

**Table S4.** ACC (in bold) followed by TNR and TPR for a dependency model with a flat prior over a variety of cardinality triples (card(*V*) : *V* ∈ (*Y, π, X*)) appearing in the first column.

| card | N | MDL |  |  | BDeu |  |  | K2 |  |  | MU |  |  |
| --- | --- | --- | --- | --- | --- | --- | --- | --- | --- | --- | --- | --- | --- |
| $(2, 3 \times 2 \times 3, 2)$ | 100 | **0.53** | 1.00 | 0.05 | **0.73** | 0.96 | 0.50 | **0.76** | 0.75 | 0.77 | **0.76** | 0.83 | 0.68 |
|  | 200 | **0.54** | 1.00 | 0.09 | **0.80** | 0.99 | 0.62 | **0.86** | 0.85 | 0.87 | **0.81** | 0.96 | 0.66 |
|  | 300 | **0.58** | 1.00 | 0.15 | **0.85** | 1.00 | 0.70 | **0.91** | 0.91 | 0.91 | **0.83** | 0.98 | 0.68 |
|  | 400 | **0.61** | 1.00 | 0.23 | **0.88** | 1.00 | 0.76 | **0.94** | 0.94 | 0.94 | **0.85** | 0.99 | 0.71 |
|  | 500 | **0.65** | 1.00 | 0.30 | **0.90** | 1.00 | 0.81 | **0.95** | 0.95 | 0.95 | **0.87** | 0.99 | 0.75 |
|  | 600 | **0.69** | 1.00 | 0.37 | **0.92** | 1.00 | 0.84 | **0.96** | 0.96 | 0.96 | **0.88** | 0.99 | 0.77 |
| $(3, 3 \times 2 \times 3, 3)$ | 100 | **0.50** | 1.00 | 0.00 | **0.65** | 1.00 | 0.31 | **0.81** | 0.73 | 0.89 | **0.81** | 0.73 | 0.89 |
|  | 200 | **0.50** | 1.00 | 0.00 | **0.69** | 1.00 | 0.37 | **0.92** | 0.87 | 0.96 | **0.94** | 0.96 | 0.92 |
|  | 300 | **0.50** | 1.00 | 0.00 | **0.74** | 1.00 | 0.47 | **0.96** | 0.94 | 0.99 | **0.97** | 0.99 | 0.94 |
|  | 400 | **0.50** | 1.00 | 0.00 | **0.78** | 1.00 | 0.57 | **0.98** | 0.97 | 0.99 | **0.98** | 1.00 | 0.96 |
|  | 500 | **0.50** | 1.00 | 0.00 | **0.83** | 1.00 | 0.65 | **0.99** | 0.98 | 1.00 | **0.99** | 1.00 | 0.97 |
|  | 600 | **0.50** | 1.00 | 0.01 | **0.87** | 1.00 | 0.73 | **1.00** | 0.99 | 1.00 | **0.99** | 1.00 | 0.98 |
| $(4, 3 \times 2 \times 3, 4)$ | 100 | **0.50** | 1.00 | 0.00 | **0.57** | 1.00 | 0.14 | **0.80** | 0.67 | 0.92 | **0.82** | 0.71 | 0.92 |
|  | 200 | **0.50** | 1.00 | 0.00 | **0.56** | 1.00 | 0.11 | **0.91** | 0.84 | 0.99 | **0.96** | 0.95 | 0.97 |
|  | 300 | **0.50** | 1.00 | 0.00 | **0.57** | 1.00 | 0.14 | **0.96** | 0.93 | 1.00 | **0.99** | 0.99 | 0.99 |
|  | 400 | **0.50** | 1.00 | 0.00 | **0.59** | 1.00 | 0.17 | **0.99** | 0.97 | 1.00 | **1.00** | 1.00 | 0.99 |
|  | 500 | **0.50** | 1.00 | 0.00 | **0.62** | 1.00 | 0.23 | **0.99** | 0.99 | 1.00 | **1.00** | 1.00 | 1.00 |
|  | 600 | **0.50** | 1.00 | 0.00 | **0.65** | 1.00 | 0.30 | **1.00** | 1.00 | 1.00 | **1.00** | 1.00 | 1.00 |
| $(5, 3 \times 2 \times 3, 5)$ | 100 | **0.50** | 1.00 | 0.00 | **0.54** | 1.00 | 0.07 | **0.77** | 0.59 | 0.94 | **0.84** | 0.81 | 0.87 |
|  | 200 | **0.50** | 1.00 | 0.00 | **0.51** | 1.00 | 0.02 | **0.88** | 0.77 | 0.99 | **0.96** | 0.95 | 0.97 |
|  | 300 | **0.50** | 1.00 | 0.00 | **0.51** | 1.00 | 0.02 | **0.95** | 0.90 | 1.00 | **0.99** | 0.99 | 0.99 |
|  | 400 | **0.50** | 1.00 | 0.00 | **0.51** | 1.00 | 0.02 | **0.98** | 0.96 | 1.00 | **1.00** | 1.00 | 1.00 |
|  | 500 | **0.50** | 1.00 | 0.00 | **0.51** | 1.00 | 0.02 | **0.99** | 0.99 | 1.00 | **1.00** | 1.00 | 1.00 |
|  | 600 | **0.50** | 1.00 | 0.00 | **0.52** | 1.00 | 0.03 | **1.00** | 0.99 | 1.00 | **1.00** | 1.00 | 1.00 |
| $(6, 3 \times 2 \times 3, 6)$ | 100 | **0.50** | 1.00 | 0.00 | **0.52** | 1.00 | 0.04 | **0.73** | 0.52 | 0.95 | **0.78** | 0.94 | 0.62 |
|  | 200 | **0.50** | 1.00 | 0.00 | **0.50** | 1.00 | 0.01 | **0.84** | 0.69 | 0.99 | **0.95** | 0.98 | 0.92 |
|  | 300 | **0.50** | 1.00 | 0.00 | **0.50** | 1.00 | 0.00 | **0.92** | 0.83 | 1.00 | **0.99** | 0.99 | 0.99 |
|  | 400 | **0.50** | 1.00 | 0.00 | **0.50** | 1.00 | 0.00 | **0.96** | 0.92 | 1.00 | **1.00** | 1.00 | 1.00 |
|  | 500 | **0.50** | 1.00 | 0.00 | **0.50** | 1.00 | 0.00 | **0.98** | 0.97 | 1.00 | **1.00** | 1.00 | 1.00 |
|  | 600 | **0.50** | 1.00 | 0.00 | **0.50** | 1.00 | 0.00 | **0.99** | 0.99 | 1.00 | **1.00** | 1.00 | 1.00 |
| $(7, 3 \times 2 \times 3, 7)$ | 100 | **0.50** | 1.00 | 0.00 | **0.52** | 1.00 | 0.03 | **0.71** | 0.46 | 0.95 | **0.60** | 1.00 | 0.21 |
|  | 200 | **0.50** | 1.00 | 0.00 | **0.50** | 1.00 | 0.00 | **0.79** | 0.59 | 1.00 | **0.86** | 1.00 | 0.72 |
|  | 300 | **0.50** | 1.00 | 0.00 | **0.50** | 1.00 | 0.00 | **0.87** | 0.75 | 1.00 | **0.98** | 1.00 | 0.95 |
|  | 400 | **0.50** | 1.00 | 0.00 | **0.50** | 1.00 | 0.00 | **0.93** | 0.87 | 1.00 | **1.00** | 1.00 | 0.99 |
|  | 500 | **0.50** | 1.00 | 0.00 | **0.50** | 1.00 | 0.00 | **0.97** | 0.94 | 1.00 | **1.00** | 1.00 | 1.00 |
|  | 600 | **0.50** | 1.00 | 0.00 | **0.50** | 1.00 | 0.00 | **0.99** | 0.97 | 1.00 | **1.00** | 1.00 | 1.00 |
| $(8, 3 \times 2 \times 3, 8)$ | 100 | **0.50** | 1.00 | 0.00 | **0.51** | 1.00 | 0.03 | **0.68** | 0.41 | 0.95 | **0.51** | 1.00 | 0.03 |
|  | 200 | **0.50** | 1.00 | 0.00 | **0.50** | 1.00 | 0.00 | **0.75** | 0.51 | 1.00 | **0.66** | 1.00 | 0.32 |
|  | 300 | **0.50** | 1.00 | 0.00 | **0.50** | 1.00 | 0.00 | **0.82** | 0.65 | 1.00 | **0.89** | 1.00 | 0.79 |
|  | 400 | **0.50** | 1.00 | 0.00 | **0.50** | 1.00 | 0.00 | **0.89** | 0.78 | 1.00 | **0.98** | 1.00 | 0.97 |
|  | 500 | **0.50** | 1.00 | 0.00 | **0.50** | 1.00 | 0.00 | **0.94** | 0.88 | 1.00 | **1.00** | 1.00 | 1.00 |
|  | 600 | **0.50** | 1.00 | 0.00 | **0.50** | 1.00 | 0.00 | **0.97** | 0.94 | 1.00 | **1.00** | 1.00 | 1.00 |
| $(9, 3 \times 2 \times 3, 9)$ | 100 | **0.50** | 1.00 | 0.00 | **0.51** | 1.00 | 0.03 | **0.66** | 0.37 | 0.95 | **0.50** | 1.00 | 0.00 |
|  | 200 | **0.50** | 1.00 | 0.00 | **0.50** | 1.00 | 0.00 | **0.71** | 0.43 | 1.00 | **0.53** | 1.00 | 0.06 |
|  | 300 | **0.50** | 1.00 | 0.00 | **0.50** | 1.00 | 0.00 | **0.77** | 0.55 | 1.00 | **0.70** | 1.00 | 0.40 |
|  | 400 | **0.50** | 1.00 | 0.00 | **0.50** | 1.00 | 0.00 | **0.84** | 0.68 | 1.00 | **0.91** | 1.00 | 0.82 |
|  | 500 | **0.50** | 1.00 | 0.00 | **0.50** | 1.00 | 0.00 | **0.90** | 0.80 | 1.00 | **0.99** | 1.00 | 0.97 |
|  | 600 | **0.50** | 1.00 | 0.00 | **0.50** | 1.00 | 0.00 | **0.94** | 0.89 | 1.00 | **1.00** | 1.00 | 1.00 |

**Table S5.** ACC (in bold) followed by TNR and TPR for a dependency model with Jeffrey’s prior over a variety of cardinality triples (card(*V*) : *V* ∈ (*Y, π, X*)) appearing in the first column, evaluated for BDeu with *α* ∈ {1, 10, 20, 100}.

| card | N | BDeu 1 |  |  | BDeu 10 |  |  | BDeu 20 |  |  | BDeu 100 |  |  |
| --- | --- | --- | --- | --- | --- | --- | --- | --- | --- | --- | --- | --- | --- |
| (2, 2, 2) | 100 | **0.81** | 0.96 | 0.66 | **0.78** | 0.73 | 0.83 | **0.73** | 0.59 | 0.87 | **0.62** | 0.32 | 0.93 |
|  | 200 | **0.86** | 0.98 | 0.75 | **0.83** | 0.79 | 0.88 | **0.77** | 0.64 | 0.91 | **0.64** | 0.33 | 0.95 |
|  | 300 | **0.89** | 0.98 | 0.79 | **0.86** | 0.82 | 0.90 | **0.80** | 0.68 | 0.92 | **0.65** | 0.34 | 0.96 |
|  | 400 | **0.90** | 0.99 | 0.82 | **0.88** | 0.84 | 0.91 | **0.82** | 0.70 | 0.94 | **0.66** | 0.36 | 0.97 |
|  | 500 | **0.91** | 0.99 | 0.84 | **0.89** | 0.85 | 0.92 | **0.83** | 0.72 | 0.94 | **0.67** | 0.37 | 0.97 |
|  | 600 | **0.92** | 0.99 | 0.85 | **0.90** | 0.86 | 0.93 | **0.84** | 0.74 | 0.95 | **0.68** | 0.38 | 0.97 |
| (3, 3, 3) | 100 | **0.86** | 1.00 | 0.72 | **0.94** | 0.97 | 0.92 | **0.92** | 0.87 | 0.96 | **0.68** | 0.36 | 1.00 |
|  | 200 | **0.92** | 1.00 | 0.85 | **0.98** | 0.99 | 0.97 | **0.96** | 0.93 | 0.99 | **0.70** | 0.39 | 1.00 |
|  | 300 | **0.95** | 1.00 | 0.91 | **0.99** | 0.99 | 0.98 | **0.97** | 0.95 | 0.99 | **0.72** | 0.43 | 1.00 |
|  | 400 | **0.97** | 1.00 | 0.94 | **0.99** | 1.00 | 0.99 | **0.98** | 0.96 | 1.00 | **0.73** | 0.46 | 1.00 |
|  | 500 | **0.98** | 1.00 | 0.96 | **1.00** | 1.00 | 0.99 | **0.98** | 0.97 | 1.00 | **0.74** | 0.49 | 1.00 |
|  | 600 | **0.98** | 1.00 | 0.97 | **1.00** | 1.00 | 1.00 | **0.99** | 0.98 | 1.00 | **0.76** | 0.51 | 1.00 |
| (4, 4, 4) | 100 | **0.76** | 1.00 | 0.53 | **0.94** | 1.00 | 0.87 | **0.97** | 0.99 | 0.95 | **0.80** | 0.60 | 1.00 |
|  | 200 | **0.86** | 1.00 | 0.72 | **0.98** | 1.00 | 0.97 | **0.99** | 1.00 | 0.99 | **0.85** | 0.69 | 1.00 |
|  | 300 | **0.92** | 1.00 | 0.84 | **0.99** | 1.00 | 0.99 | **1.00** | 1.00 | 1.00 | **0.87** | 0.75 | 1.00 |
|  | 400 | **0.96** | 1.00 | 0.91 | **1.00** | 1.00 | 0.99 | **1.00** | 1.00 | 1.00 | **0.89** | 0.79 | 1.00 |
|  | 500 | **0.97** | 1.00 | 0.95 | **1.00** | 1.00 | 1.00 | **1.00** | 1.00 | 1.00 | **0.91** | 0.82 | 1.00 |
|  | 600 | **0.98** | 1.00 | 0.97 | **1.00** | 1.00 | 1.00 | **1.00** | 1.00 | 1.00 | **0.92** | 0.85 | 1.00 |
| (5, 5, 5) | 100 | **0.66** | 1.00 | 0.32 | **0.85** | 1.00 | 0.70 | **0.93** | 1.00 | 0.86 | **0.92** | 0.85 | 1.00 |
|  | 200 | **0.70** | 1.00 | 0.41 | **0.95** | 1.00 | 0.90 | **0.99** | 1.00 | 0.97 | **0.96** | 0.93 | 1.00 |
|  | 300 | **0.78** | 1.00 | 0.56 | **0.98** | 1.00 | 0.96 | **1.00** | 1.00 | 0.99 | **0.98** | 0.96 | 1.00 |
|  | 400 | **0.85** | 1.00 | 0.70 | **0.99** | 1.00 | 0.99 | **1.00** | 1.00 | 1.00 | **0.99** | 0.97 | 1.00 |
|  | 500 | **0.91** | 1.00 | 0.81 | **1.00** | 1.00 | 0.99 | **1.00** | 1.00 | 1.00 | **0.99** | 0.98 | 1.00 |
|  | 600 | **0.94** | 1.00 | 0.88 | **1.00** | 1.00 | 1.00 | **1.00** | 1.00 | 1.00 | **0.99** | 0.99 | 1.00 |
| (6, 6, 6) | 100 | **0.59** | 1.00 | 0.19 | **0.72** | 1.00 | 0.45 | **0.83** | 1.00 | 0.65 | **0.97** | 0.96 | 0.99 |
|  | 200 | **0.58** | 1.00 | 0.15 | **0.83** | 1.00 | 0.66 | **0.94** | 1.00 | 0.87 | **1.00** | 0.99 | 1.00 |
|  | 300 | **0.60** | 1.00 | 0.20 | **0.91** | 1.00 | 0.83 | **0.98** | 1.00 | 0.96 | **1.00** | 1.00 | 1.00 |
|  | 400 | **0.65** | 1.00 | 0.29 | **0.96** | 1.00 | 0.92 | **0.99** | 1.00 | 0.99 | **1.00** | 1.00 | 1.00 |
|  | 500 | **0.70** | 1.00 | 0.40 | **0.98** | 1.00 | 0.97 | **1.00** | 1.00 | 1.00 | **1.00** | 1.00 | 1.00 |
|  | 600 | **0.76** | 1.00 | 0.52 | **0.99** | 1.00 | 0.99 | **1.00** | 1.00 | 1.00 | **1.00** | 1.00 | 1.00 |
| (7, 7, 7) | 100 | **0.57** | 1.00 | 0.14 | **0.62** | 1.00 | 0.25 | **0.70** | 1.00 | 0.41 | **0.97** | 0.99 | 0.94 |
|  | 200 | **0.52** | 1.00 | 0.05 | **0.67** | 1.00 | 0.33 | **0.80** | 1.00 | 0.61 | **1.00** | 1.00 | 1.00 |
|  | 300 | **0.52** | 1.00 | 0.05 | **0.74** | 1.00 | 0.48 | **0.89** | 1.00 | 0.79 | **1.00** | 1.00 | 1.00 |
|  | 400 | **0.53** | 1.00 | 0.06 | **0.82** | 1.00 | 0.64 | **0.95** | 1.00 | 0.90 | **1.00** | 1.00 | 1.00 |
|  | 500 | **0.54** | 1.00 | 0.09 | **0.89** | 1.00 | 0.77 | **0.98** | 1.00 | 0.96 | **1.00** | 1.00 | 1.00 |
|  | 600 | **0.56** | 1.00 | 0.13 | **0.94** | 1.00 | 0.87 | **0.99** | 1.00 | 0.98 | **1.00** | 1.00 | 1.00 |
| (8, 8, 8) | 100 | **0.57** | 1.00 | 0.13 | **0.57** | 1.00 | 0.14 | **0.61** | 1.00 | 0.23 | **0.90** | 1.00 | 0.81 |
|  | 200 | **0.51** | 1.00 | 0.02 | **0.56** | 1.00 | 0.12 | **0.65** | 1.00 | 0.29 | **0.98** | 1.00 | 0.96 |
|  | 300 | **0.50** | 1.00 | 0.01 | **0.58** | 1.00 | 0.16 | **0.71** | 1.00 | 0.42 | **1.00** | 1.00 | 0.99 |
|  | 400 | **0.50** | 1.00 | 0.01 | **0.62** | 1.00 | 0.24 | **0.79** | 1.00 | 0.58 | **1.00** | 1.00 | 1.00 |
|  | 500 | **0.51** | 1.00 | 0.01 | **0.67** | 1.00 | 0.34 | **0.86** | 1.00 | 0.72 | **1.00** | 1.00 | 1.00 |
|  | 600 | **0.51** | 1.00 | 0.01 | **0.73** | 1.00 | 0.47 | **0.92** | 1.00 | 0.83 | **1.00** | 1.00 | 1.00 |
| (9, 9, 9) | 100 | **0.58** | 1.00 | 0.17 | **0.55** | 1.00 | 0.09 | **0.56** | 1.00 | 0.13 | **0.81** | 1.00 | 0.61 |
|  | 200 | **0.50** | 1.00 | 0.01 | **0.52** | 1.00 | 0.04 | **0.55** | 1.00 | 0.10 | **0.91** | 1.00 | 0.83 |
|  | 300 | **0.50** | 1.00 | 0.00 | **0.52** | 1.00 | 0.04 | **0.57** | 1.00 | 0.14 | **0.97** | 1.00 | 0.94 |
|  | 400 | **0.50** | 1.00 | 0.00 | **0.52** | 1.00 | 0.05 | **0.60** | 1.00 | 0.20 | **0.99** | 1.00 | 0.98 |
|  | 500 | **0.50** | 1.00 | 0.00 | **0.54** | 1.00 | 0.07 | **0.65** | 1.00 | 0.30 | **1.00** | 1.00 | 1.00 |
|  | 600 | **0.50** | 1.00 | 0.00 | **0.56** | 1.00 | 0.11 | **0.70** | 1.00 | 0.41 | **1.00** | 1.00 | 1.00 |

**Table S6.**
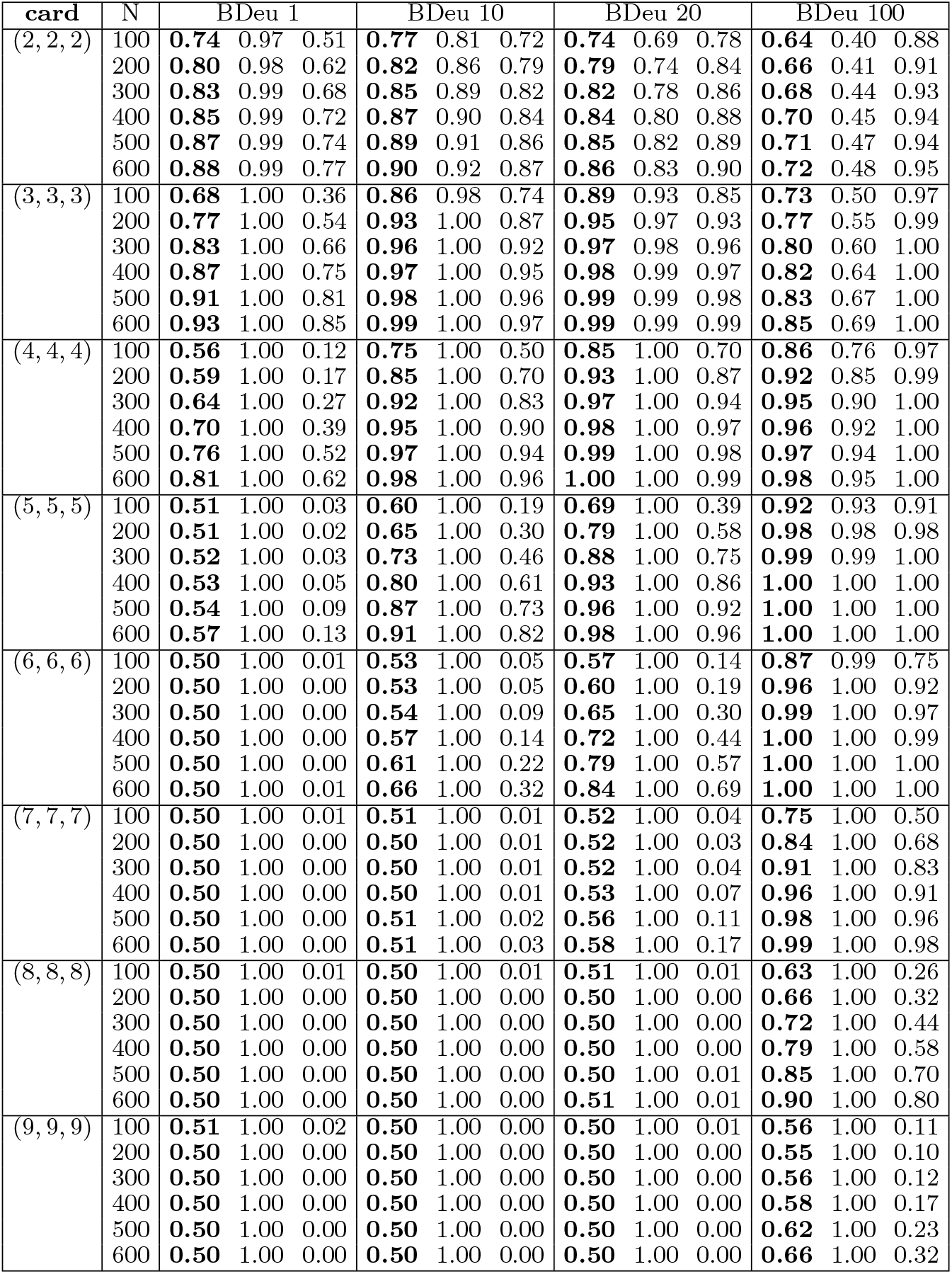
ACC (in bold) followed by TNR and TPR for a dependency model with flat prior over a variety of cardinality triples (card(*V*) : *V* ∈ (*Y, π, X*)) appearing in the first column. for BDeu with *α* ∈ {1, 10, 20, 100}.

### Supplement B

**Fig. S1.**
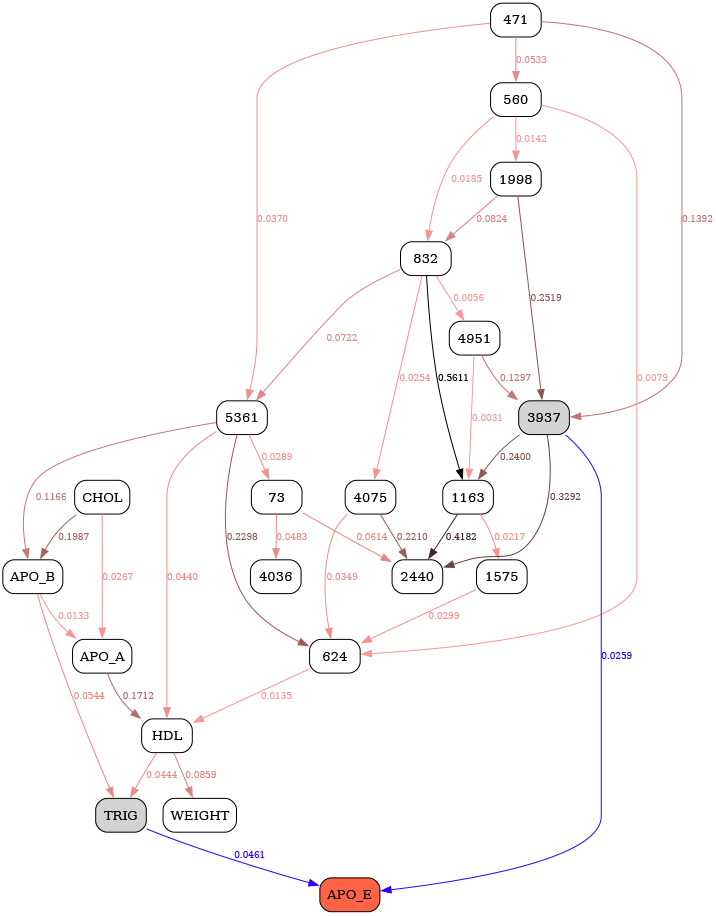
MDL 3-state.

**Fig. S2.**
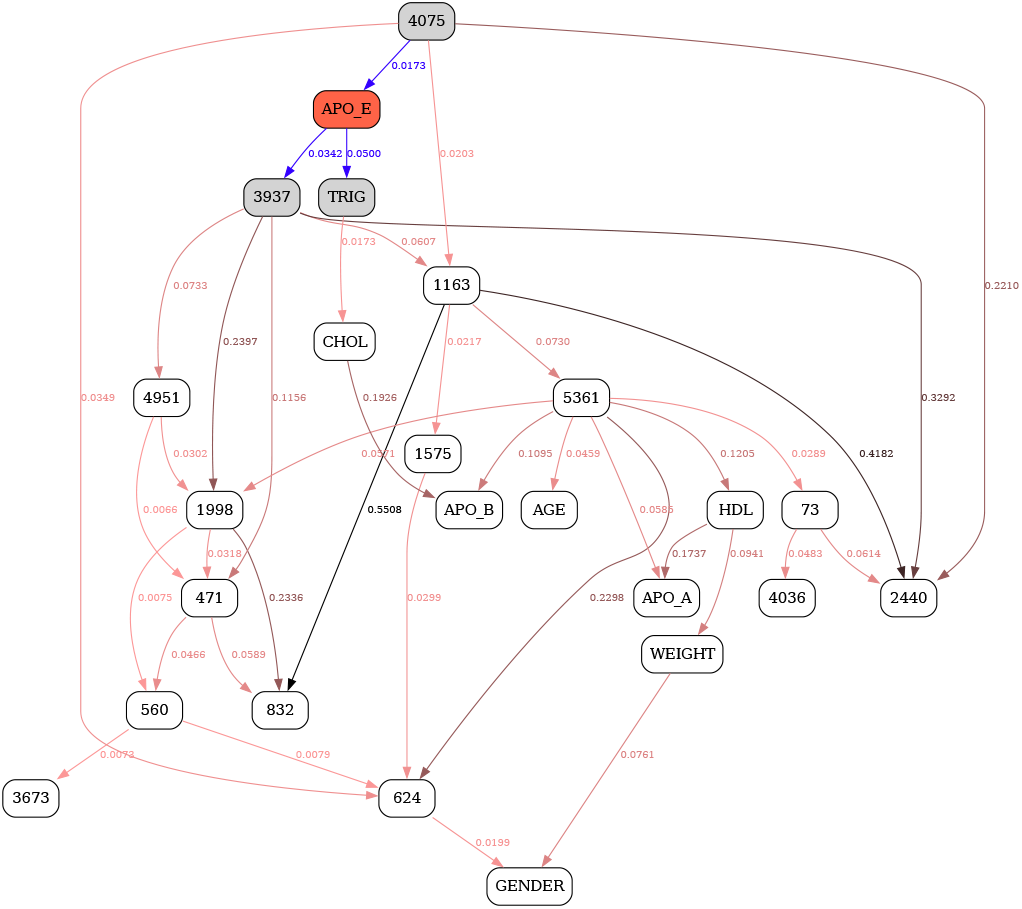
MDL 5-state.

**Fig. S3.**
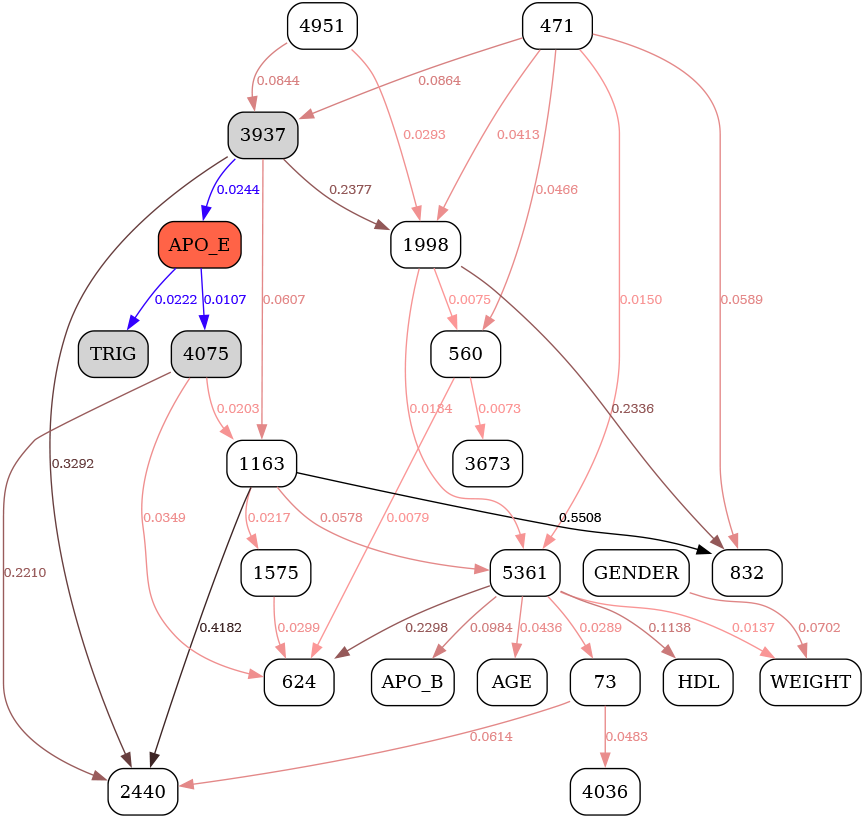
MDL 7-state.

**Fig. S4.**
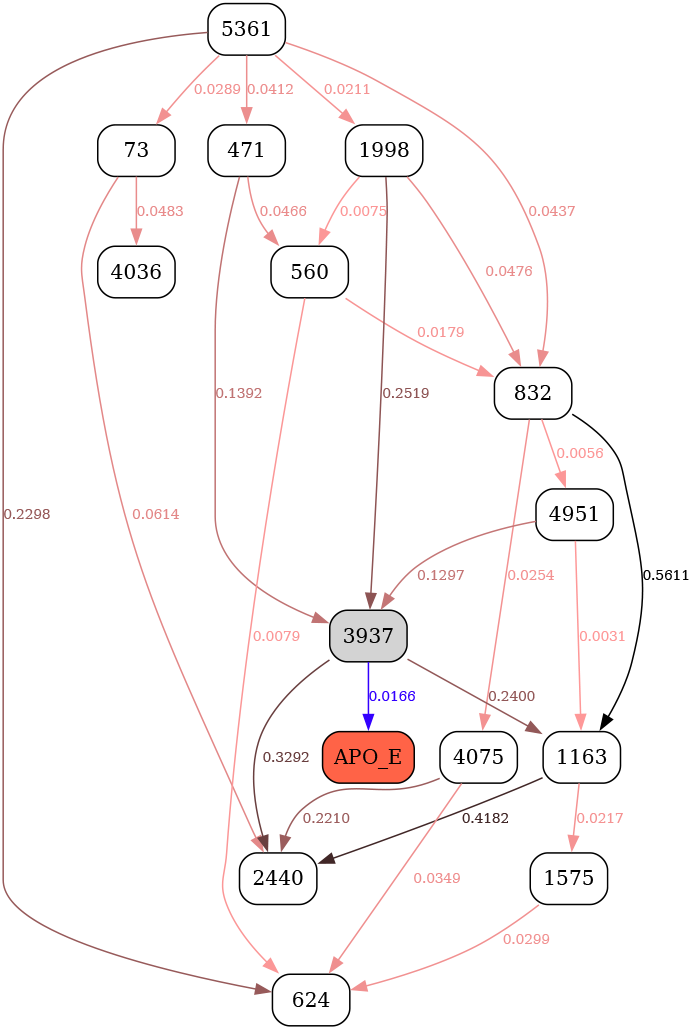
MDL 9-state.

**Fig. S5.**
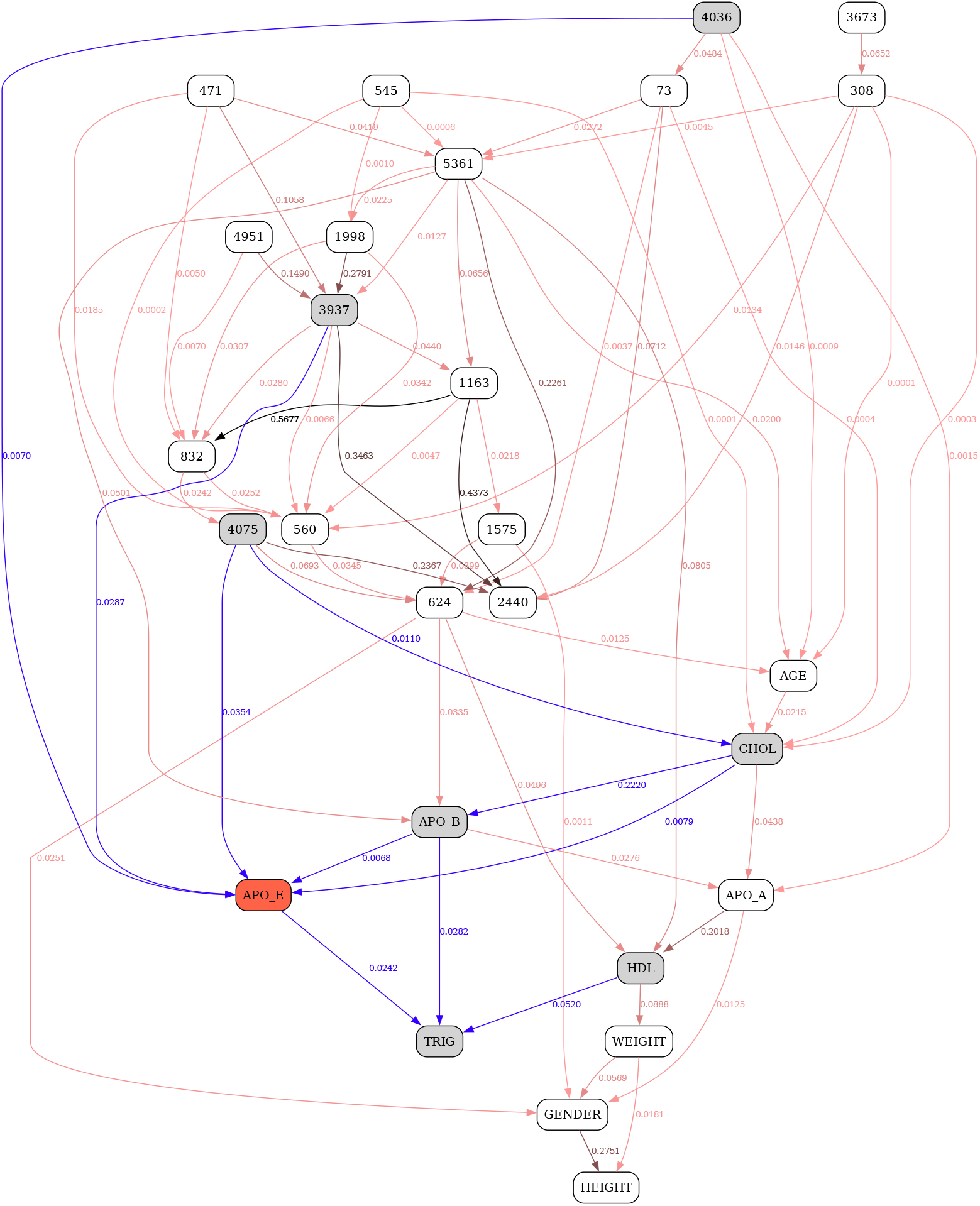
K2 3-state.

**Fig. S6.**
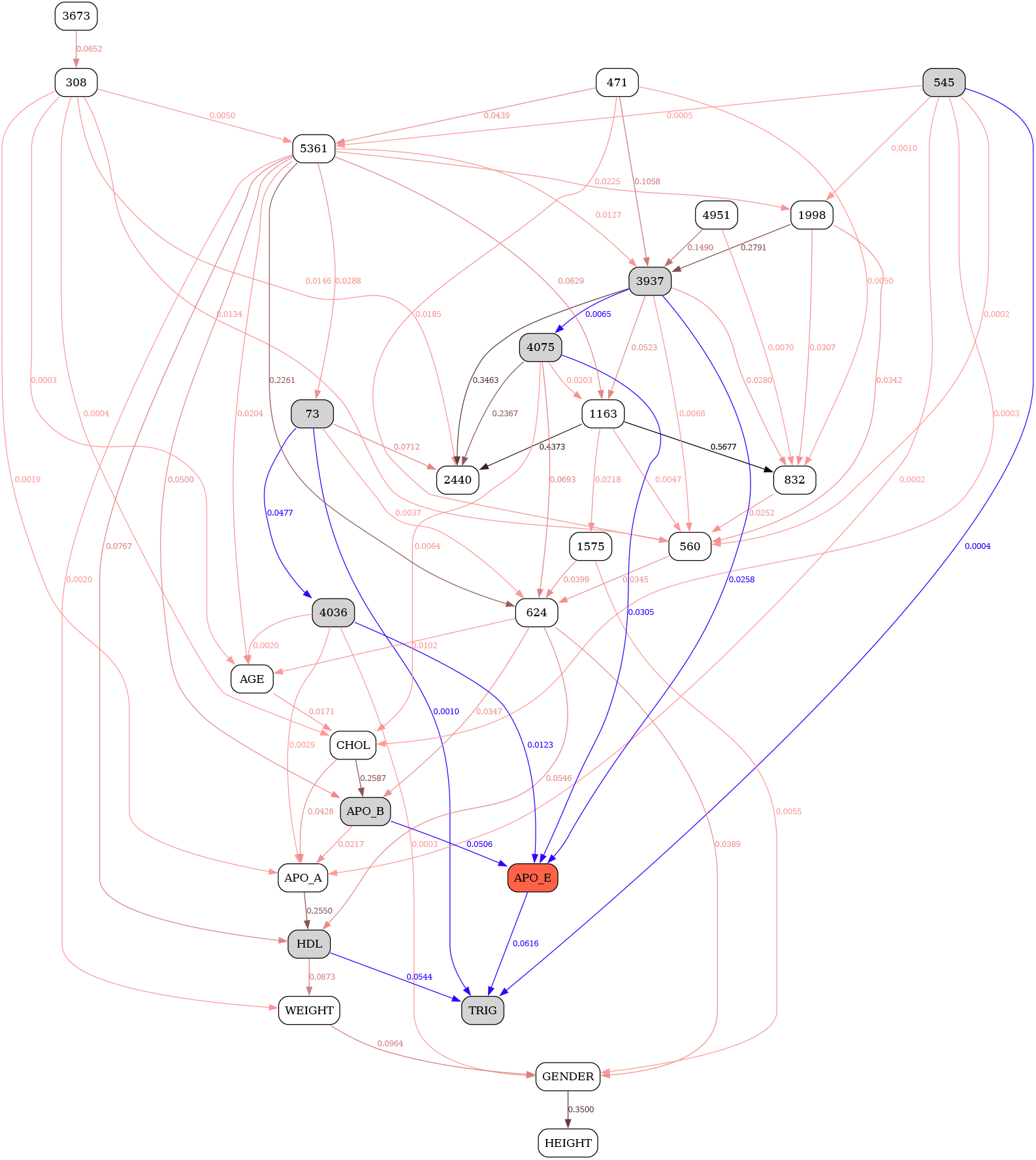
K2 5-state.

**Fig. S7.**
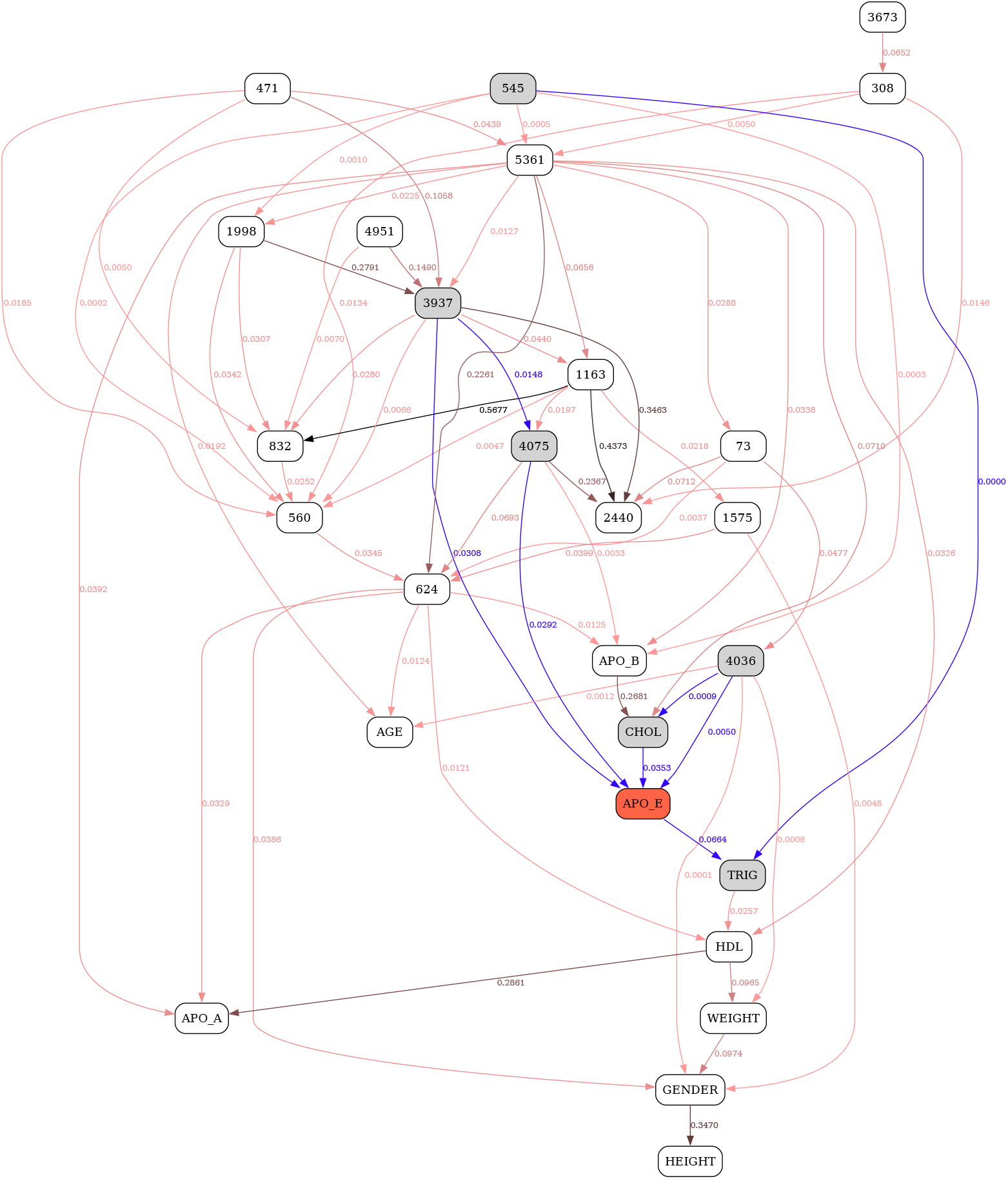
K2 7-state.

**Fig. S8.**
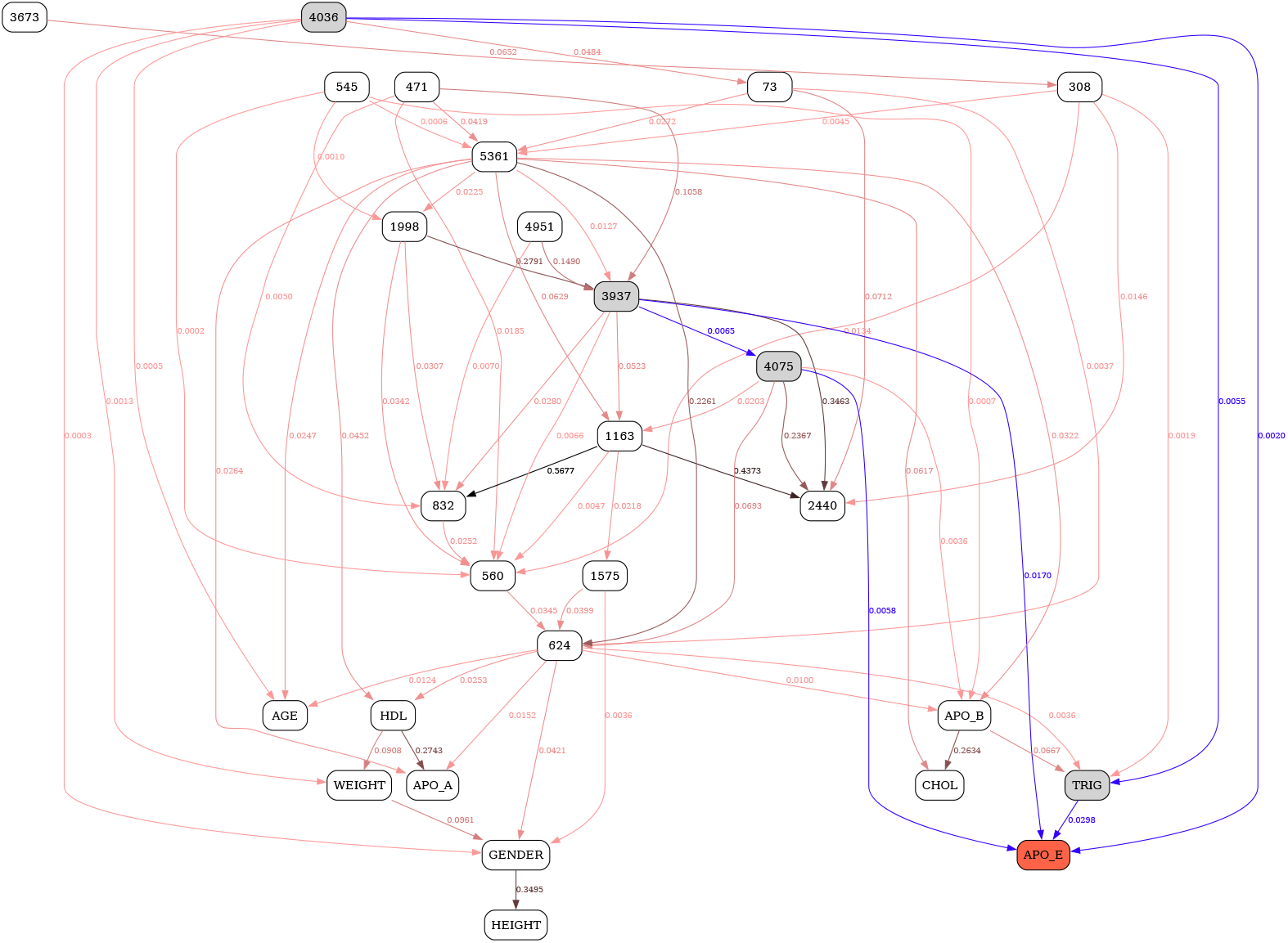
K2 9-state.

**Fig. S9.**
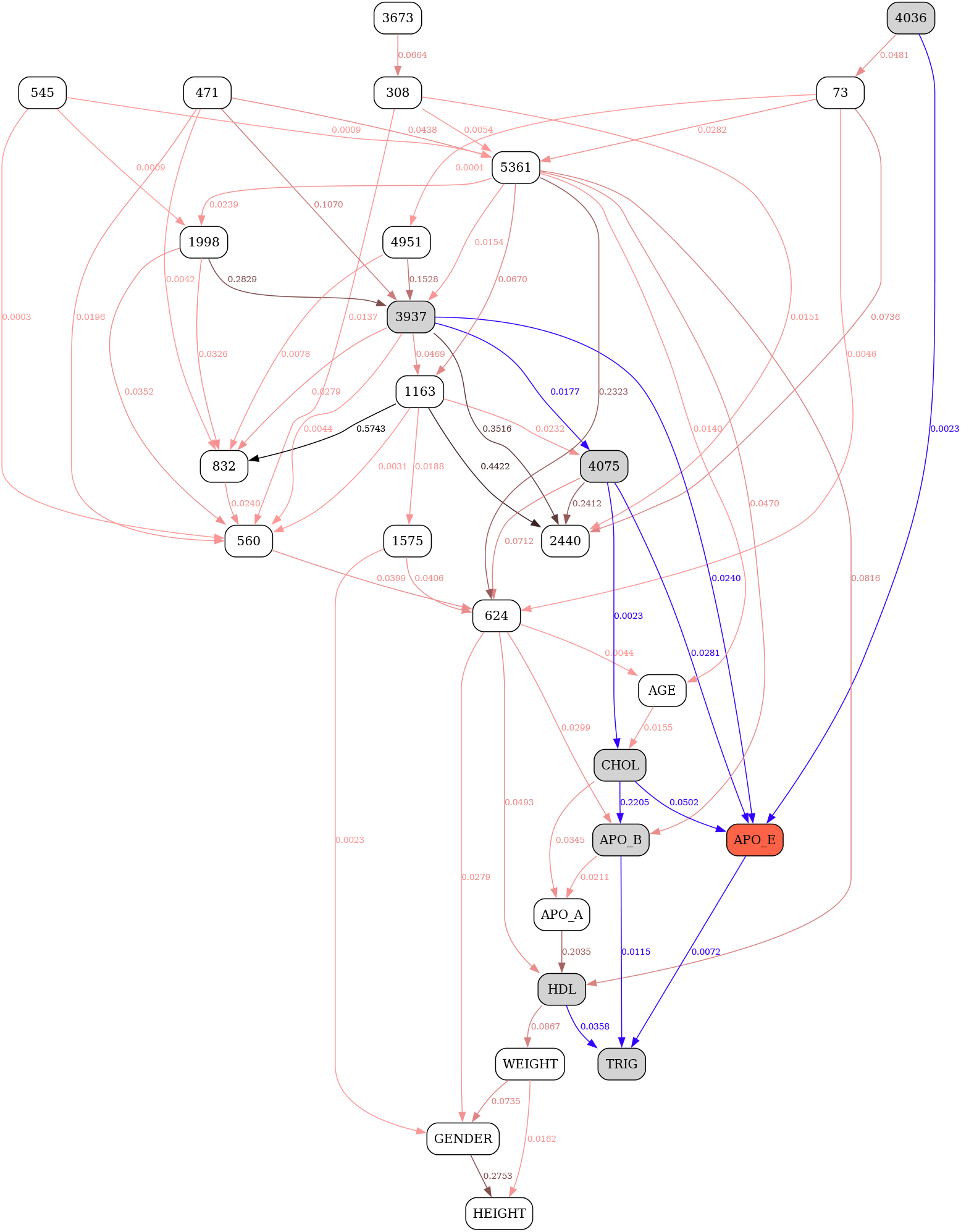
MU 3-state.

**Fig. S10.**
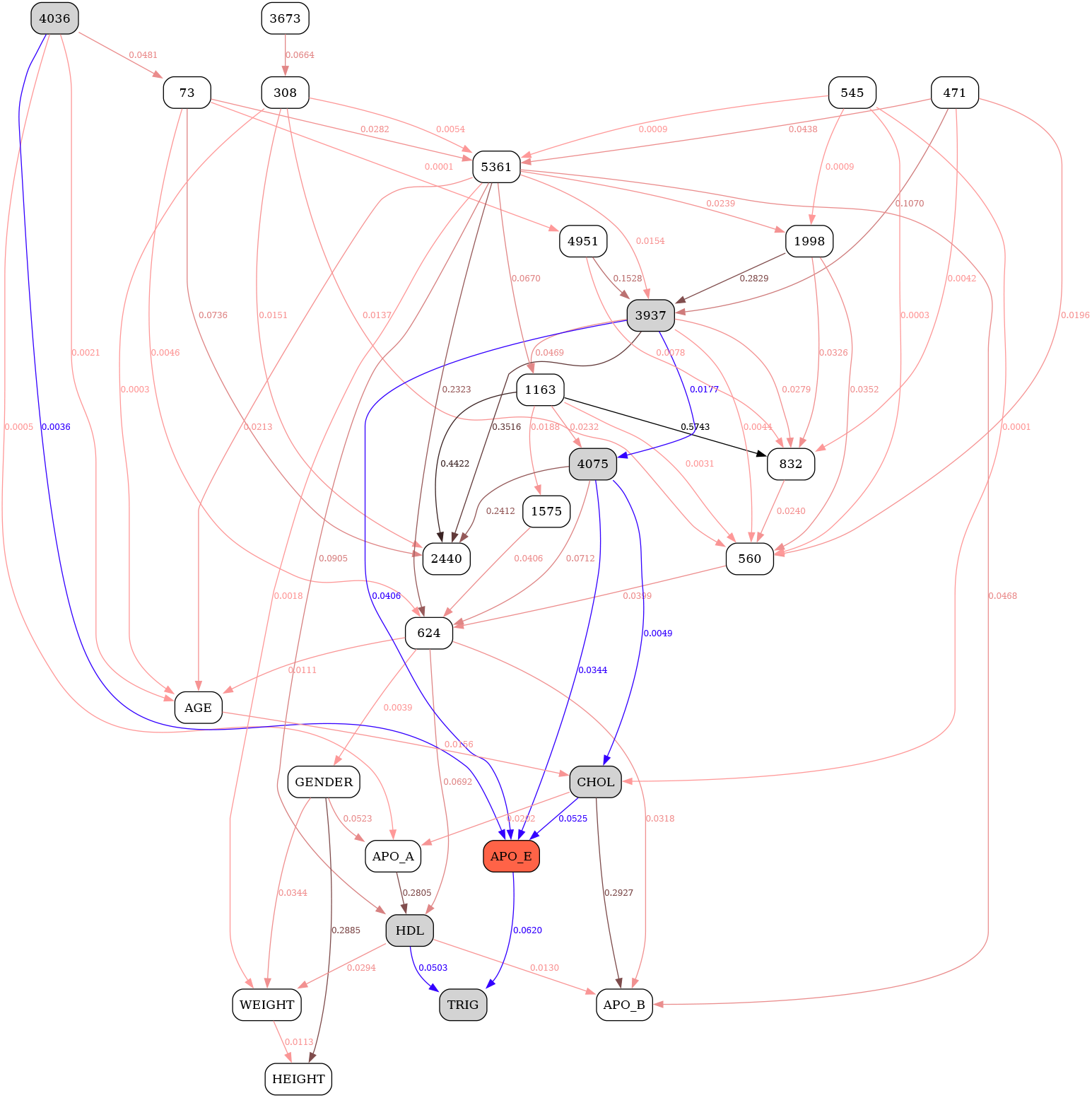
MU 5-state.

**Fig. S11.**
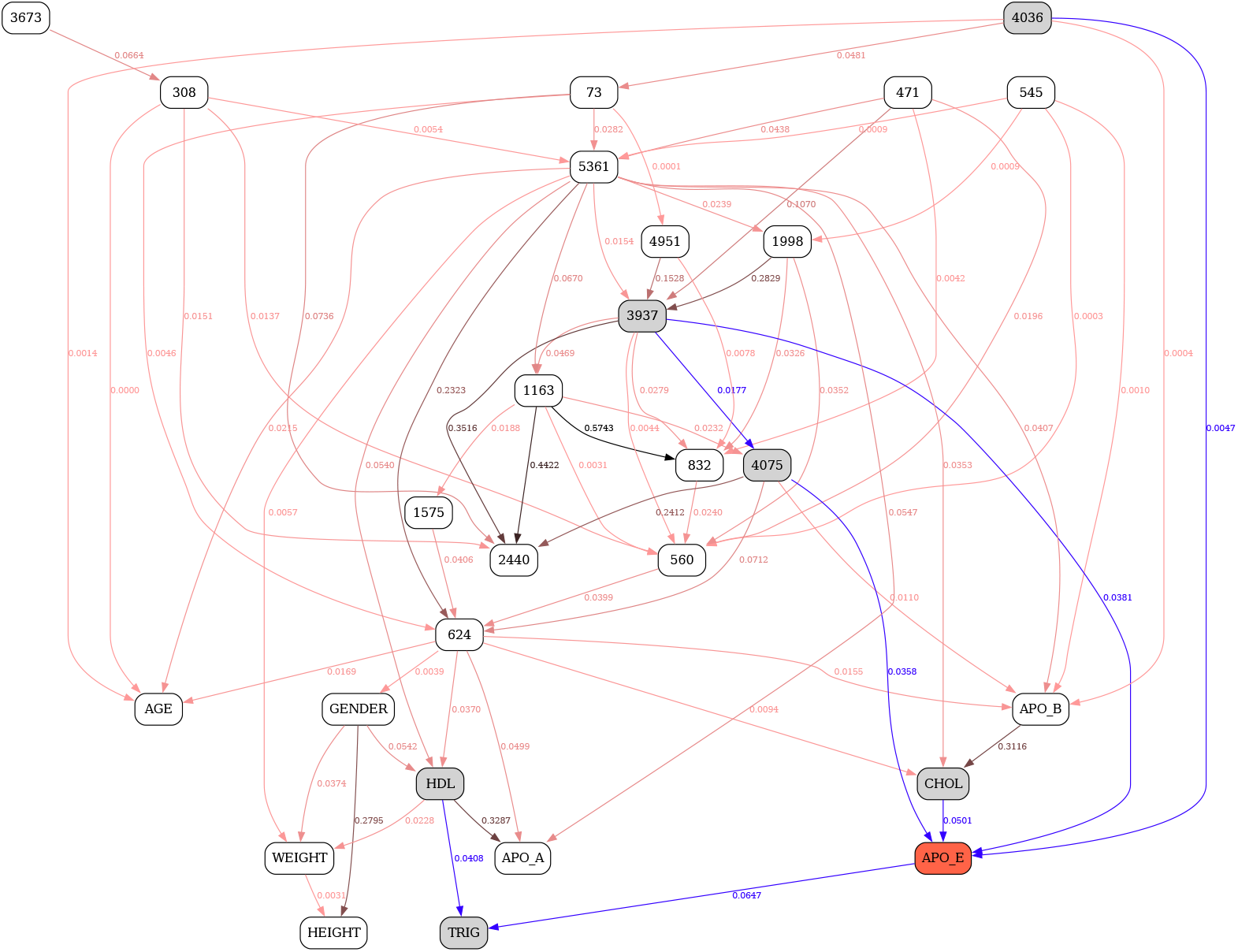
MU 7-state.

**Fig. S12.**
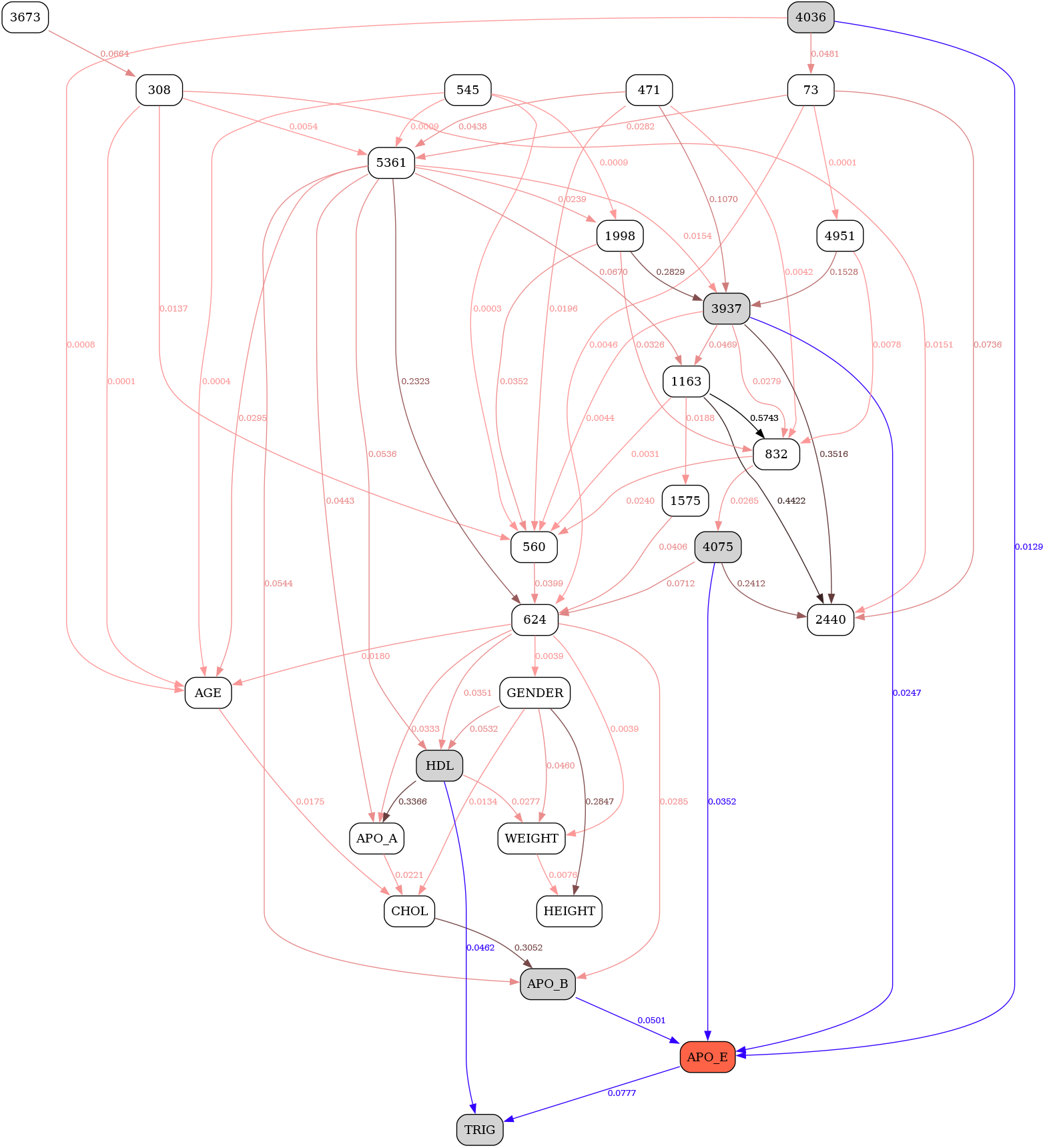
MU 9-state.

## Notes

### Competing Interest Statement

The authors have declared no competing interest.

### Summary of Updates

Inconsistencies and conceptual ambiguities were fixed. Abstract was streamlined into a much more compact form. Motivation was substantially expanded, more details added. New section documenting a concrete example of MDL failure was created. The application of the calibrated MU criterion to the same example, indicating successful attainment of design goals, was added. Section interpreting the new criterion was expanded and refined. The tables were recalculated to remove pasting inconsistencies and moved into the supplement. New tables were added to include inter-BDeu comparison. New figures with full BNs were added to the supplement.

